# FRAME-tags: genetically encoded fluorescent markers for multiplexed barcoding and time-resolved tracking of live cells

**DOI:** 10.1101/2021.04.09.436507

**Authors:** Andrew V. Anzalone, Miguel Jimenez, Virginia W. Cornish

**Author notes:** These authors contributed equally to this work (Andrew V. Anzalone, Miguel Jimenez). Correspondence: Andrew V. Anzalone, Miguel Jimenez, or Virginia W. Cornish.

## Abstract

Cellular barcodes offer critical tools for tracking cellular identity in biological systems. Although genetically encoded fluorescent barcodes are ideal for real-time tracking, their scalability is constrained by the broad, overlapping emission spectra characteristic of fluorescent proteins (FPs). Here, we describe a palette of genetically encoded fluorescent barcodes called FRAME-tags, which break this scalability barrier by encoding barcode identity as unique FP expression ratios. FRAME-tags use −1 programmed ribosomal frameshifting RNA motifs to precisely control the translational output of multiple FPs from a single mRNA, leading to extremely narrow and resolvable ratios of the corresponding cellular fluorescence distributions. With this platform, we constructed 20 resolvable FRAME-tags in yeast using just two FPs, and further demonstrated that 100 or more distinguishable FRAME-tags could be made by the addition of a third FP. We used FRAME-tags to map the dynamic fitness landscape of yeast co-cultures, and to characterize the expression pattern of 20 yeast promoters in multiplex across diverse conditions. FRAME-tags offer a valuable new tool for cellular barcoding that enables time-resolved characterization of complex biological systems using widely available fluorescence detection techniques and a minimal number of spectral channels.

## INTRODUCTION

Biological systems commonly contain multiple distinct but morphologically indistinguishable cell types, challenging our ability to study subsets of cells *in situ* at the individual cell level^1^. Moreover, natural and engineered biological systems are extremely dynamic—their cellular compositions evolve over time, their cells divide at varying rates, and they react to changes in their external environment. As increasingly complex biological systems are studied and engineered^2–7^, new methods are required that allow for cellular subpopulations to be uniquely identified in their native context and easily tracked over time.

Cell identification and tracking can be achieved using cellular barcodes, which link the identity of a cell to an observable readout. Ideally, such barcodes can be scaled indefinitely, contain completely orthogonal and distinguishable identifiers, and can be conveniently analyzed in a non-destructive manner using widely accessible laboratory techniques. A number of methods exist for cellular barcoding, such as those based on DNA sequences^8–10^ or exogenously applied labels^11–13^. While DNA sequence-based tags are highly scalable and capable of capturing system-wide snapshots of large cell populations, they are not easily adapted for closely tracking populations of cells over time, or in real time. Moreover, sequencing-based methods require cells to be taken out of their natural context or destroyed prior to analysis^14–16^. Therefore, alternative cell barcoding tools are needed for time-resolved cellular identification, lineage tracking, and phenotypic reporting in multiplex from intact biological systems.

Fluorescent proteins (FPs) are attractive alternatives for constructing cellular barcodes because they are genetically encoded and therefore do not dilute over multiple cell generations, they emit fluorescence signals that can be easily measured directly from samples using microscopy or flow cytometry, and they come in a wide variety of colors^17^. However, despite these advantages, FPs have broad absorption and fluorescence emission spectra that limit the number of variants that can be resolved simultaneously. This restricts single-FP-based barcoding to experiments that contain only a small number of cell types (typically three to five), and it necessitates the use of advanced instrumentation to fully resolve all barcodes^18^.

To bypass this FP scalability challenge, elegant combinatorial approaches have been devised that utilize expression of multiple FPs, thereby mixing several base colors to generate a larger number of resolvable colors^19–21^. Of note, the Brainbow method allows from three to up to approximately 100 theoretically unique FP combinations to be generated from a single genetic construct through stochastic recombination events^22^. While broadly useful, these combinatorial methods have several limitations. First, they exhaust most available fluorescence channels because of their dependence on three or more FP colors, limiting their use alongside other fluorescent reporters. Second, discrimination between barcodes of similar hues becomes increasingly difficult with increasing barcode numbers, requiring the use of additional spatial information to identify specific cells. Finally, these methods rely on the stochastic integration of multiple copies of a genetic construct, making the total barcode diversity variable across experiments and precluding *a priori* barcode assignment to a specific subset of cells.

We sought to overcome the limitations of previous FP barcoding systems by using −1 programmed ribosomal frameshifting to produce cell-identifying ratios of fluorescence proteins. Our approach preserves all of the advantages of FP-based barcoding while using a minimal number of fluorescence channels and single-copy integrated genetic constructs to create a large set of robust and resolvable tags. To this end, we developed a palette of ratiometric fluorescent barcodes that we call FRAME-tags (<u>F</u>rameshift-controlled <u>RA</u>tiometric <u>M</u>ulti-fluorescent protein <u>E</u>xpression tags) that can achieve 20 or more unique tags in yeast using just two fluorescent proteins. Here, we demonstrate that FRAME-tags are small, portable genetic constructs that can be robustly identified in real-time under varying conditions with just two-color microscopes and flow cytometers.

## RESULTS

### Building the FRAME-tag palette

We hypothesized that a large number of unique fluorescent barcodes could be generated from a minimal set of FPs if their absolute and relative expression levels could be precisely set within each cell. We further hypothesized that a co-translational mechanism could offer the highest degree of precision for regulating protein synthesis ratios. In particular, we identified −1 programmed ribosomal frameshifting (−1 PRF) as a promising mechanism to encode FP production ratios since it uses self-contained RNA elements, it is active in a wide variety of organisms, and its activity can be tuned by the choice of a frameshift stimulating RNA motif ^23^. Previously, we reported a large collection of −1 PRF signals that possess frameshifting efficiencies spanning two orders of magnitude in yeast (~0.2% to ~30%) ^24^. In the current work, we designed discrete frameshift modules (*fs*) that incorporate these −1 PRF signals for modular assembly of a FRAME-tag palette (**Fig. 1a** and **Supplementary Fig. 1**). Our *fs* modules were designed so that a failure to frameshift would result in the immediate termination of translation at a proximal stop codon, whereas successful frameshifting would allow for continued translation in the −1 reading frame. This, in principle, leads to a precise synthesis ratio of the proteins upstream and downstream of the *fs* module, determined by the overall strength of the frameshift signal (**Fig. 1b**).

**Figure 1.**
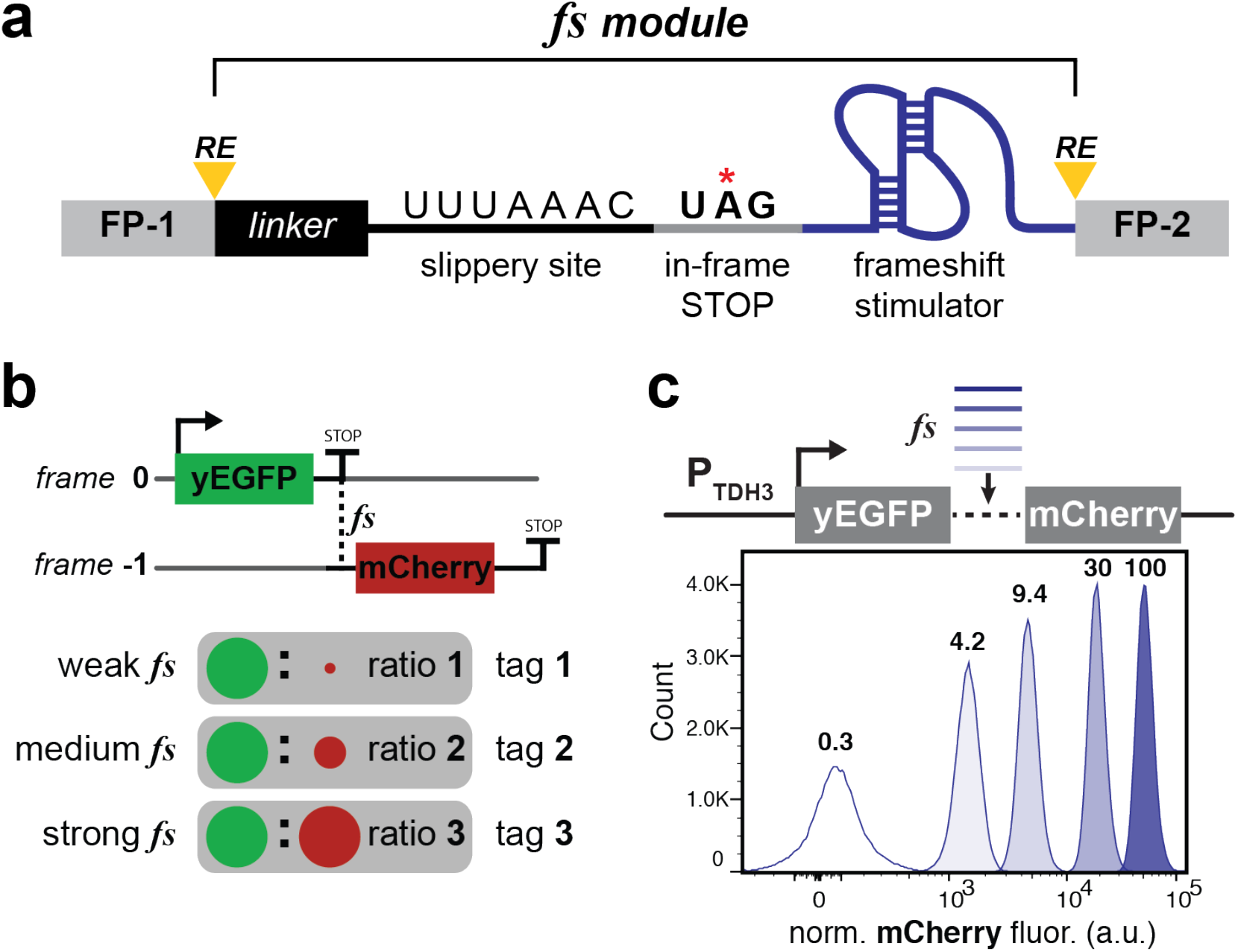
Frameshift modules encode precise FP ratios. **(a)** Frameshift (*fs*) modules are designed to control the stoichiometry of upstream and downstream open reading frames (FP-1 and FP-2) via −1 PRF. They contain a peptide linker, a heptanucleotide tRNA slippery site, an in-frame stop codon, and a custom frameshift stimulatory RNA signal. *fs* modules are flanked by restriction sites (RE) for convenient cloning. (**b**) At the *fs* modules, translation either terminates or continues in the −1 reading frame. Distinct ratios of the upstream and downstream proteins (yEGFP and mCherry) are produced based on *fs* module frameshift efficiency. These ratios can be used to uniquely tag cells with specific ratios of FP-based fluorescence signals. (**c**) Selected *fs* modules were inserted between yEGFP and mCherry, then chromosomally integrated into yeast; mCherry fluorescence was evaluated by flow cytometry and normalized by side scatter; values written above the distribution curves indicate frameshift efficiency determined by comparison of mCherry/yEGFP ratios to the 100% yEGFP-mCherry fusion (see **Supplementary Fig. 2**).

To begin constructing and validating the FRAME-tag palette, *fs* modules^24^ were first screened in a yeast dual-FP reporter assay wherein *fs* modules were inserted between an upstream yEGFP encoded in the +1 reading frame and a downstream mCherry encoded in the −1 reading frame (**Supplementary Fig. 2** and **Supplementary Table 1**). Each translation event produces a yEGFP protein, but only those translation events that frameshift at the *fs* module result in mCherry production. From this screen, we identified a set of five *fs* modules whose mCherry fluorescence distributions were highly resolvable, displaying less than 1% overlap between adjacent populations (**Fig. 1c**). In addition, flow cytometry analysis revealed that these *fs* modules produced distributions of mCherry:yEGFP fluorescence ratios that had even greater resolution. (**Supplementary Fig. 3**). We reasoned that these narrow distributions of fluorescence ratios were due to co-translation of the fluorescent proteins from a single mRNA, making the relative ratio of FPs robust to intrinsic biological noise originating from variability in transcription, translation initiation, and mRNA stability^25,26^. Moreover, we found that absolute fluorescence could be made robust to extrinsic biological noise by single-copy chromosomal integration into the same genomic locus in yeast and expression from the TDH3 promoter (**Supplementary Fig. 3**).

Next, using these five validated *fs* modules, we designed two-FP FRAME-tag constructs that contain yEGFP and mCherry reading frames, wherein each FP is controlled by an upstream *fs* module (**Fig. 2a** and **Supplementary Table 2**). In this design, translation of each FP requires successful frameshifting at all upstream *fs* modules, so the expected FP yield is predicted to be the product of all preceding frameshifting efficiencies. We constructed a set of 20 FRAME-tags variants with combinations of *fs* modules predicted to yield resolvable FP ratios and integrated TDH3-controlled FRAME-tags at the *Leu2* locus in yeast. When analyzed by flow cytometry, these 20 FRAME-tags were found to be highly resolvable (**Fig. 2a** and **Supplementary Fig. 4**). The observed FP signals detected from each FRAME-tag construct were in good agreement with the predicted expression levels based on *fs* module strengths, with some deviation that depended on the location and identity of the RNA signals (**Supplementary Fig. 5**).

**Figure 2.**
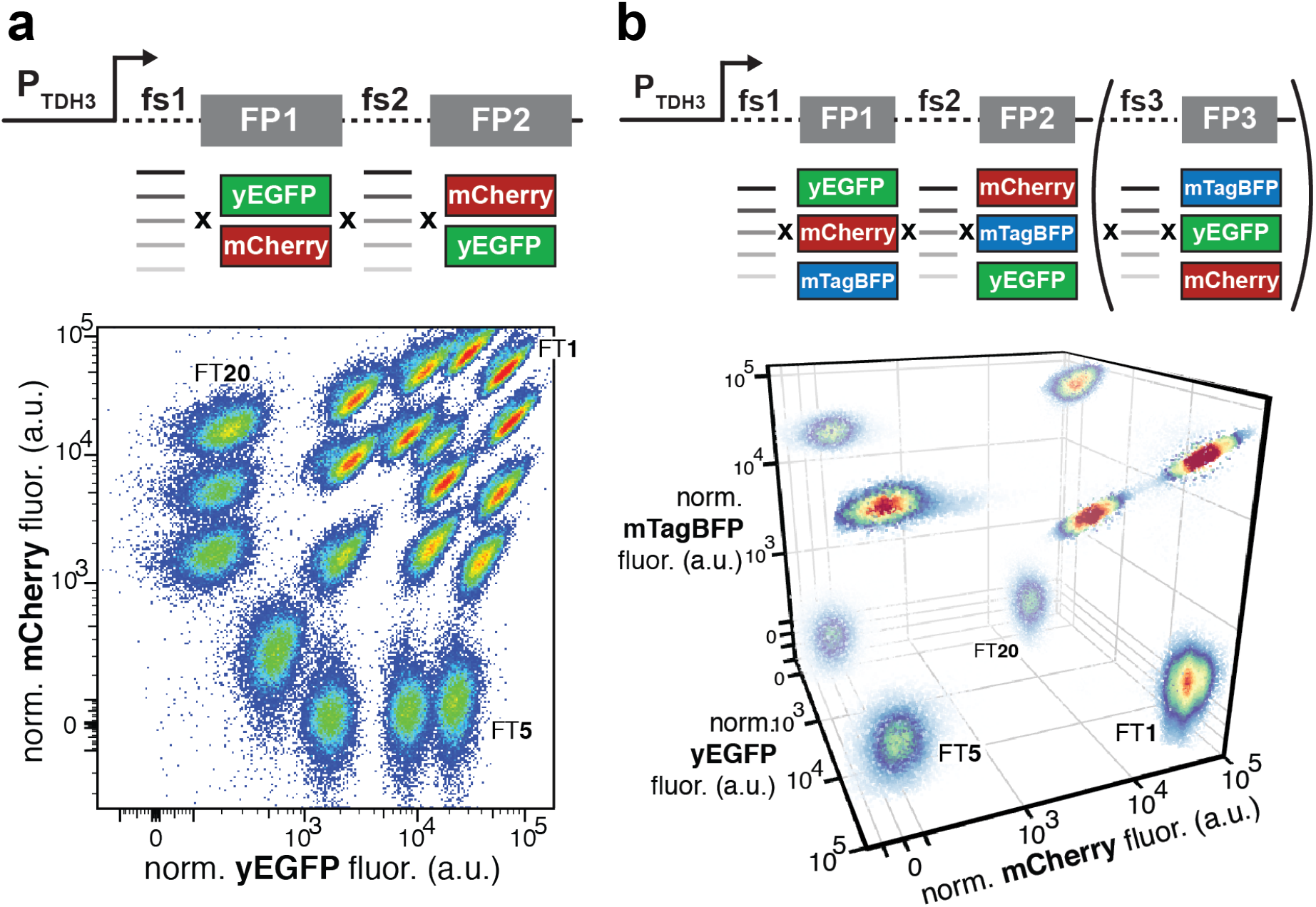
Two-color and three-color FRAME-tags generate highly resolvable cell populations. (**a**) Two-color FRAME-tags were constructed by assembling two FPs (yEGFP and mCherry) in either orientation, each paired with one of the five *fs* modules shown in **Fig. 1c**. FRAME-tag constructs were then chromosomally integrated at the *Leu2* locus in yeast, and their fluorescence was analyzed by flow cytometry (fluorescence signal normalized by side scatter); data displayed as a scatter plot from twenty FRAME-tag strains grown in a mixed culture, pseudo-colored by event density. (**b**) The FRAME-tag design was expanded to three FPs (yEGFP, mCherry and mTagBFP2) to generate two-FP and three-FP FRAME-tags (include bracketed module). Three-color FRAME-tags were analyzed as in panel **a** and visualized in a three-dimensional scatter plot pseudo-colored by event density. Some FRAME-tags (FT1, FT5, FT20) are labeled for reference between panels **a** and **b**.

Next, we expanded our FRAME-tag series to a third color dimension by the introduction of an additional FP variant. After screening several FPs, we determined that the blue mTagBFP2^27^ is orthogonal to the yEGFP and mCherry fluorescence channels (**Supplementary Fig. 6**). To establish the scalability of our FRAME-tag palette, we generated three additional two-FP FRAME-tags containing mTagBFP2–yEGFP or mTagBFP2–mCherry pairs. We also designed a FRAME-tag architecture that contains all three FPs regulated by upstream *fs* modules and validated this design by the construction of two additional FRAME-tags (**Fig. 2b** and **Supplementary Fig. 7**). Based on these results demonstrating consistently narrow fluorescence distributions for both two-color and three-color FRAME-tag populations, we estimate that up to 100 unique and resolvable FRAME-tags could fit within this three-color FP space (**Supplementary Note 1**).

To streamline data analysis for FRAME-tag applications, we developed an automated flow cytometry gating and analysis pipeline (FRAME-finder) based on the R package openCyto^28^ (**Supplementary Note 2**). FRAME-finder exploits the characteristically narrow and predictable fluorescence distributions of FRAME-tagged cells to automatically segment mixed populations into unique FRAME-tag bins using simple one-dimensional gates. (**Supplementary Fig. 8**). Statistical analysis of this gating algorithm revealed that it could identify FRAME-tagged populations of cells with gate positive predictive values (PPV, probability that an event falling within the gate is a true event) ranging from 0.99 to >0.9999 (mean gate PPV of 0.9987) at a population gating threshold of 90% (**Supplementary Fig. 9**). By extrapolation of these data, we predict that a large majority of FRAME-tagged strains could be detected with a PPV of greater than 0.9 for dilutions as low as 1 in >10^3^ cells while maintaining a population gating threshold of 50% (**Supplementary Fig. 10**).

Next, we asked if fluorescence microscopy could also be used to identify FRAME-tags. Using two-color microscopy, we imaged samples of 10^4^ yEGFP–mCherry FRAME-tagged yeast cells and determined fluorescence intensity values for each cell by image segmentation (**Fig. 3a**). Using this microscopy-derived data as input, FRAME-finder enabled simultaneous assignment of all 20 yEGFP–mCherry FRAME-tag variants across more than 90% of the population (**Fig. 3b** and **Supplementary Figs. 11-12**). The high fidelity of FRAME-tag identification was supported by independent assignment of the same FRAME-tag barcode to adjacent mother-daughter cell pairs that are expected to be genetically identical (**Fig. 3c**).

**Figure 3.**
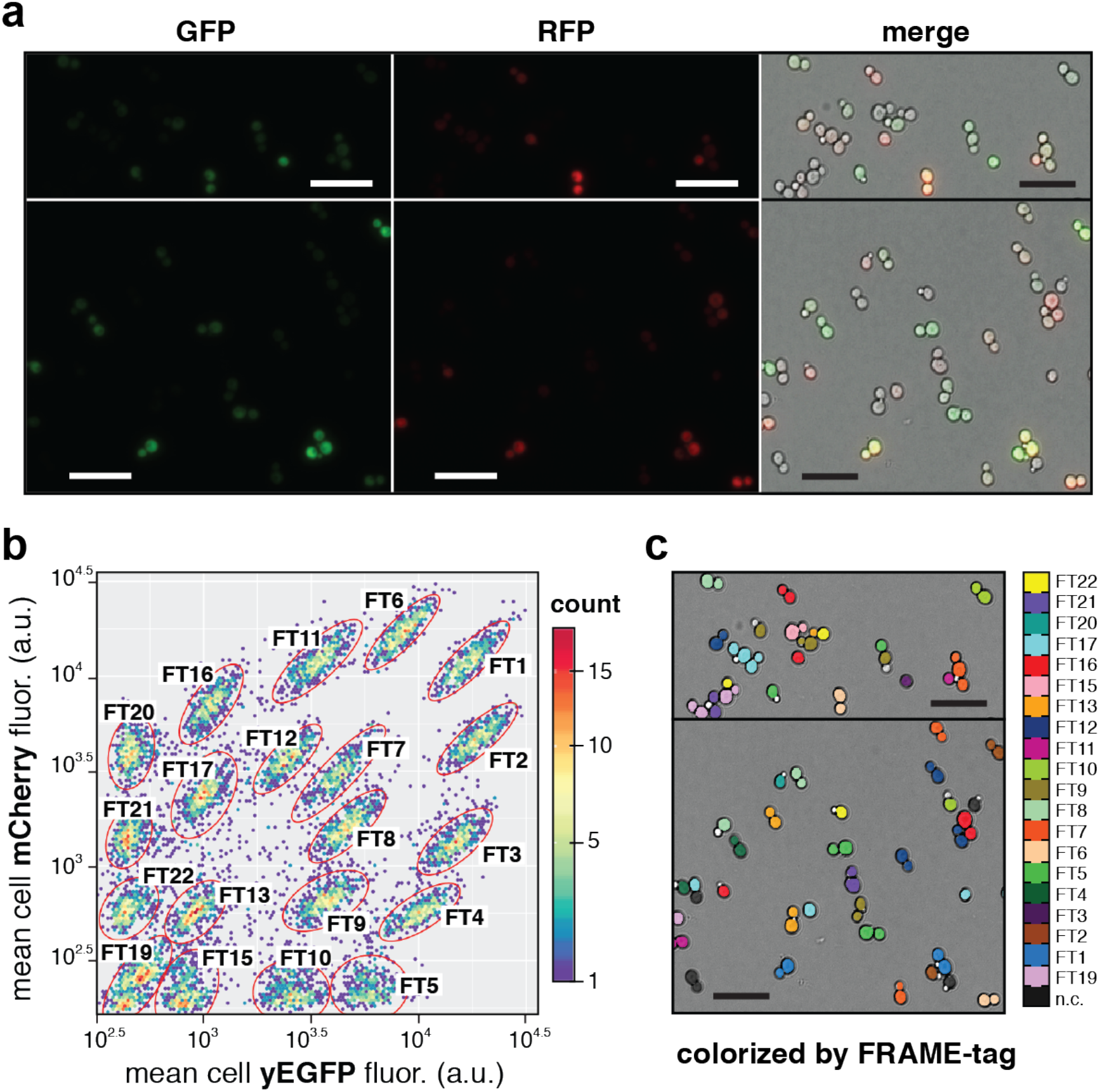
Identification of FRAME-tags using fluorescence microscopy. (**a**) Fluorescence microscopy images of a mixture of live yeast containing all 20 yEGFP-mCherry FRAME-tags. Left panels, GFP channel; middle panels, RFP channel; right panels, merged fluorescence overlay with brightfield channel. Top and bottom panels show two different sub-fields. Scale bar = 20 μm. (**b**) Microscopy images of a 20 FRAME-tag co-culture that collectively captured 10^4^ cells were segmented, then yEGFP and mCherry fluorescence values from each cell were extracted and plotted as a two-dimensional scatter plot, pseudo-colored by event density. Individual FRAME-tag populations were programmatically assigned using FRAME-finder. (**c**) Cells falsely colored by FRAME-tag indices based on barcode identification in panel **b**. (see **Supplementary Fig. 11 and 12**); scale bar = 20 μm. FT# = FRAME-tag-#. n.c.: not classified.

### Tracking a complex microbial co-culture with FRAME-tags

After establishing the FRAME-tag palette and FRAME-finder tool, we sought to evaluate the robustness of FRAME-tags for long-term and continuous tracking of complex microbial co-cultures under a variety of selective environmental pressures. To streamline the generation of multiple FRAME-tag–phenotype pairs, we first validated a workflow for introducing FRAME-tags into new host yeast strains that requires just a single transformation step (**Supplementary Fig. 13**). With this method, we generated a model microbial co-culture containing nine FRAME-tagged yeast strains, each of which displays a distinct phenotype when grown in standard positive and negative selection conditions used in the field of yeast genetics^29,30^. These strains differ in their expression levels of the histidine biosynthetic enzyme His3 and the uracil biosynthesis enzyme Ura3. His3 provides cells with the ability to synthesize the amino acid histidine and therefore grow in media lacking histidine, while expression of Ura3 makes cells sensitive to the growth inhibiting 5-fluoroorotic acid (5-FOA) (**Supplementary Fig. 14**). We verified that when grown independently, each of the nine strains, whose expression of His3 and Ura3 differ, displays a distinct growth phenotype in the two standard selective conditions (**Fig. 4a** and **Supplementary Fig. 14**).

With this complex microbial co-culture in hand, we set out to address how changes in selective pressure influence the co-culture’s composition and growth dynamics. As a first step, we used FRAME-tags to determine the change in strain abundance after growth of the co-culture in various selective media conditions (**Fig. 4b**). Given the speed and ease with which FRAME-tagged strains are analyzed in this manner, we rapidly assessed the co-culture’s overall growth (as measured by OD_600_) and strain composition (by flow cytometry) after culturing in 216 distinct conditions that varied in their concentration of histidine, 5-FOA and 3-aminotriazole (3-AT, an inhibitor of the His3 enzyme). To summarize the observed population distribution for each condition, we devised a “population distribution index” (PD index), which is a value that scores the population distribution based on the underlying population fraction of each strain (see **Fig. 4c** and **Supplementary Fig. 15** for details of the PD calculation). Positive PD index values correspond to conditions that favor the growth of strains that express higher levels of His3 and Ura3, while negative PD index values correspond to conditions that favor the growth of strains that express lower levels of His3 and Ura3. PD indices close to zero indicate a condition with a flat strain distribution (**Fig. 4a** and **Fig. 4b**).

**Figure 4.**
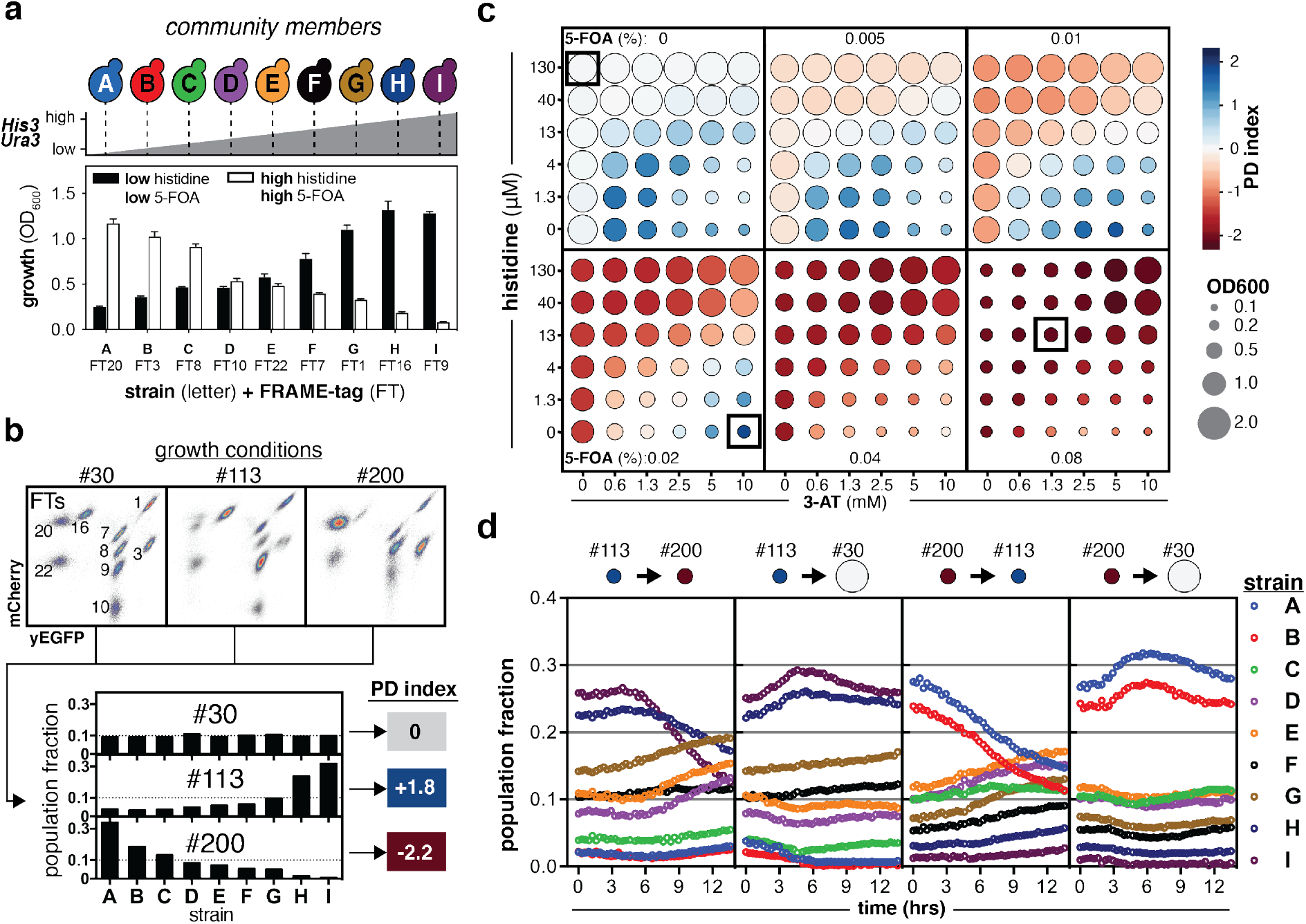
Real-time monitoring of yeast co-culture dynamics with FRAME-tags. (**a**) A synthetic yeast co-culture was made from nine individual strains with varying levels of Gal4 expression leading to distinct growth phenotypes in selective conditions (see **Supplementary Fig. 14**); each strain was barcoded with a unique FRAME-tag (A-I). Error bars represent the range of two technical replicates. (**b**) Flow cytometry analysis of the mixed co-culture is used to extract population fractions of each strain member under various culture conditions (#30, #113, #200) and can be summarized as a Population Distribution (PD) index (see **Supplementary Fig. 15**). (**c**) The 9-strain co-culture was grown in 216 culture conditions for 24 hours, then analyzed by flow cytometry to determine the PD index. The total growth of the co-culture was determined by OD_600_. Boxes highlight conditions #30, #113 and #200. (**d**) Co-culture response to abrupt changes in culture conditions (#30, #113, #200) was evaluated for four distinct transitions. Cultures were first grown in the starting condition for 24 hours, then transitioned to the new culture conditions. Cultures were sampled and quantified at 20-minute intervals by flow cytometry using FRAME-tags to determine the respective population fraction of each strain member.

As anticipated, of the 216 conditions, the two conditions that closely match the standard culture medium used to assign the individual strain phenotypes gave the expected PD index values (i.e., 0 μM His + 0% 5-FOA + 5 mM 3-AT led to a positive PD index of 1.02; 130 μM His + 0.08% 5-FOA + 0 mM 3-AT led to a negative PD index of −1.94) (**Fig. 4c**). However, unexpectedly, these two standard conditions, which were expected to provide the largest selective pressures, in fact did not give rise to the largest PD index values (i.e., the largest skew in population fractions). Instead, we found that non-standard combinations of the three medium components led to the largest effective selective pressure (e.g., 40μM His + 0.08% 5-FOA + 5 mM 3-AT led to a PD index of −2.18) while also resulting in more robust growth (OD_600_ of 0.91 vs 0.25 for the standard condition). This FRAME-tag experiment provides a detailed relational map for choosing improved His3 and Ura3 selection conditions for genetic studies and directed evolution in yeast.

Next, we leveraged our ability to track FRAME-tags in real-time to further dissect the population dynamics of our microbial co-culture when faced with transitions between differing selection conditions. Using the selection map determined above, we chose two conditions (condition #113 and #200 in **Supplementary Fig. 15)** with drastically differing selective pressures on the population structure yet matching absolute culture growth. Growth of the nine-strain co-culture was initiated in one of the two conditions. After 24 hours, cultures were abruptly transitioned to the opposite selective condition (condition #113 to #200, or vice versa) or a neutral condition (condition #30) and then tracked at 20-minute intervals by flow cytometry (**Fig. 4d**). Expectedly, cultures responded to reversals in selective pressure by favoring growth of the previously lower-abundance strains and disfavoring growth of the previously higher-abundance strains. Unexpectedly, however, transition to neutral conditions led to the collapse of low-abundance populations, and a temporary surge in growth for high-abundance strains. This temporally coupled growth surge and strain collapse would have been undetectable by other techniques that only capture endpoint measurement of the population. Overall, these results demonstrate that the FRAME-tag palette can be used to monitor dynamic growth trajectories of individual strains in real-time within complex co-cultures.

### Multiplexed fluorescent reporting with FRAME-tags

A key feature of the two-color FRAME-tag palette is its compatibility with additional fluorescent reporters, which allows for orthogonal measurements of multiple biological signals using just three fluorescence channels. To demonstrate how FRAME-tags could be applied for multiplexed reporting in this manner, we combined a mTagBFP2-based transcriptional reporter construct with the palette of 20 yEGFP–mCherry FRAME-tags. We selected 20 yeast promoters, including 18 promoters from environmentally sensitive genes and two control promoters from housekeeping genes, and cloned these promoters upstream of mTagBFP2 on CEN plasmids (see strains in **Supplementary Table 3**). Each unique mTagBFP2 reporter construct was transformed into one of the 20 yEGFP–mCherry FRAME-tagged strains to barcode the blue fluorescence signal (**Fig. 5a**).

**Figure 5.**
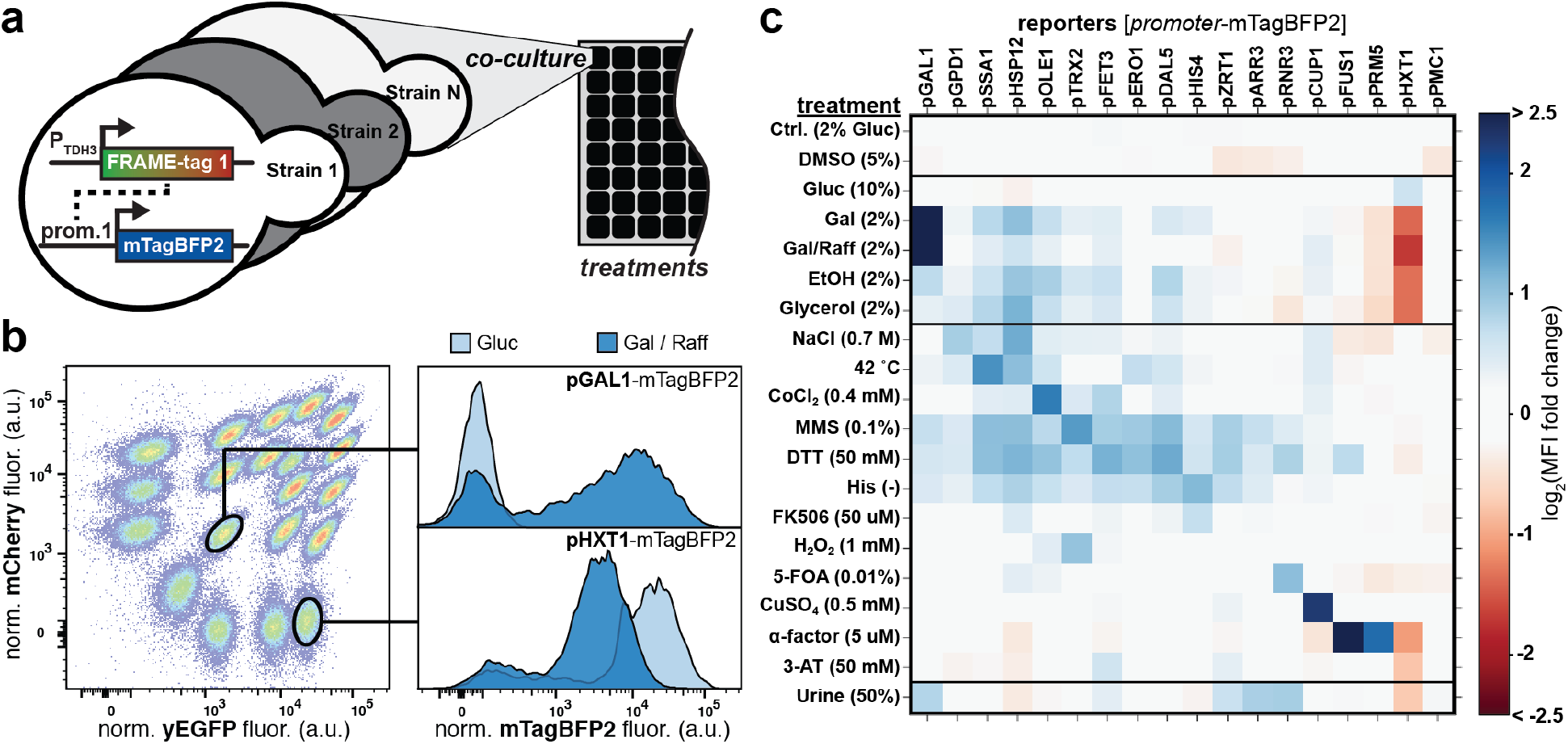
Multiplexed phenotypic profiling with FRAME-tags. (**a**) yEGFP-mCherry FRAME-tags were used to index the identity of 20 distinct promoters that drive the expression of mTagBFP2 for profiling the expression of these promoters in co-culture across 20 treatments. (**b**) FRAME-tags are analyzed by flow cytometry and used for deconvolution of the bulk blue fluorescence using FRAME-finder. Histograms are then assigned to each individual reporter construct. Examples of the deconvolved GAL1 promoter mTagBFP2 signal and HXT1 promoter mTagBFP2 signal in either Glucose (Gluc) or galactose+raffinose (Gal / Raff) conditions are shown. mTagBFP2 fluorescence was normalized by side scatter. (**c**) Expression profiles of 18 yeast promoters from mixed co-cultures following the indicated treatments; heat map represents the log_2_ of the fold change of the mean mTagBFP2 fluorescence intensity (MFI) compared to the control condition (30° C in SC dropout with 2% glucose); promoter expression for each condition was normalized to the two control promoters (pACT1 and pTEF1); cultures were analyzed by flow cytometry after 6 hours in the specified treatment. See **Supplementary Fig. 16** for individual histograms.

Using FRAME-tags, we evaluated the transcriptional responses of all 20 promoters in multiplex from samples containing all 20 FRAME-tagged reporter strains. A specific promoter’s transcriptional response was determined by flow cytometry through deconvolution of the mTagBFP2 fluorescence channel by gating the corresponding FRAME-tag sub-population (**Fig. 5b**). The ability to multiplex transcriptional reporters in this manner allows for rapid assessment of transcriptional responses under a larger number of culture conditions. Therefore, we exposed this reporter co-culture to various conditions, including 5 different carbon sources, 9 cell stress inducers, two heavy metals, a GPCR agonist, and a human biological sample. Analysis by flow cytometry and deconvolution with FRAME-finder yielded histograms corresponding to the activation profiles of individual promoters in each condition (**Fig. 5b** and **Supplementary Fig. 16**). This confirmed many known yeast transcriptional responses (**Fig. 5c**) and uncovered subtle differences in transcriptional responses between similar conditions (**Fig. 5c**, compare Gal vs. Gal/Raff). Importantly, FRAME-tag identification was robust in the face of many harsh conditions (e.g., high temperature, osmotic shock, and genotoxic agents), suggesting that FRAME-tags can be used to characterize heterogeneous populations in complex settings. Overall, these results establish that FRAME-tags can be used for multiplexed cell reporting from a single sample.

## DISCUSSION

Here, we developed FRAME-tags to overcome the scalability limit of FP-based cell barcoding by harnessing −1 programmed ribosomal frameshifting to precisely encode non-overlapping FP expression ratios. We constructed a total of 20 genetically encoded FRAME-tags using just two FPs and demonstrated the potential to scale the FRAME-tag palette to over 100 unique barcodes using just three FPs. We expect that our three-FP FRAME-tags could be scaled further to potentially 1000 tags using combinations of five FP variants^18^. Importantly, our approach achieves scalability purely based on fluorescence signals, unlike other FP barcoding methods that heavily rely on spatial discrimination (subcellular or sub-tissue) to fully distinguish similar FP color combinations^19,20^. As a result, FRAME-tags can be identified by techniques like flow cytometry and microscopy, for which we developed an automated analysis pipeline called FRAME-finder that automates the gating of FRAME-tagged cell populations for downstream analyses. Integration of FRAME-tags with spatial information, such as subcellular localization^20^, could scale the palette even further for imaging-based applications.

Recently developed DNA recording systems provide a way to track cell lineage and expression patterns from complex cellular communities^31^, however these methods rely on statistical inference from large numbers of cells and require multiple labor- and reagent-intensive steps that delay data recovery. In contrast, by exploiting the inherent speed of fluorescence data acquisition, FRAME-tags enable this analysis in real-time from intact biological samples with high time resolution and at the single cell level. We showed that FRAME-tags can indeed be used to track diverse cellular communities in real time and visualize dynamic growth trajectories for all cellular subpopulations simultaneously. This enabled us to exhaustively map the positive and negative selection conditions that could be used to direct the community composition to a target distribution or enrich a target cell type. Beyond mapping culture conditions, FRAME-tags could also be used to analyze the interactions of uncharacterized strains or phenotypes^32,33^ as well as forward-engineer specific population compositions and dynamics via synthetic intercellular communication schemes^34^. Therefore, in combination with automated flow cytometry or microscopy, FRAME-tags should enable biologists to continuously characterize natural microbial communities and synthetic biologist to rapidly iterate towards functional microbial communities.

Since FRAME-tags are pre-defined rather than stochastically defined, they can be used to uniquely link cell identity to a specified barcode. FRAME-tags also minimize the number of fluorescent channels that are required for barcode identification, allowing other orthogonal fluorescent reporters to be used alongside FRAME-tags. These reporters can be analyzed in multiplex through FRAME-tag-indexed deconvolution of the bulk reporter fluorescent signal. Using a promoter-driven orthogonal FP, we profiled expression from 20 yeast promoters across 21 experimental conditions in multiplex using FRAME-tags as promoter barcodes. Beyond promoter activity, FRAME-tags could be used to multiplex other fluorescent reporters such as those based on calcium sensing, protein-protein interaction sensing, and cell signaling^35–37^. By comparison to other multiplexing techniques such as chemical barcoding^11^, genetically-encoded FRAME-tags do not require sample staining, thereby decreasing sample-to-sample variability and preventing barcode signal dilution over time. In conjunction with other fluorescent reporters, FRAME-tags should be applicable to various multiplexed phenotypic screens including microbial expression profiling^38^, cell state reporters^39^, and drug screening^11^.

As established here, FRAME-tags should find use in a wide range of applications due to their modularity, scalability, and convenience of characterization. Our current FRAME-tag palette can be immediately used in yeast for basic biological studies and synthetic biology. In addition, it should be possible to develop FRAME-tags for bacterial and mammalian cells, where −1 ribosomal frameshifting also occurs^40^. As future tools for basic biology, FRAME-tags could find use for studying microbiome dynamics, pathogen engraftment, or lineage tracing in developing organisms and tumors^41^. Furthermore, as synthetic biology tools, FRAME-tags could be used in multicellular community engineering, distributed metabolic engineering, and biosensor arrays^42–44^. With the aid of emerging genome-engineering technologies^45^, automated cytometry^46^, and time-lapse imaging^47^, we anticipate that FRAME-tags and their future variants will find extensive use in the broader scientific community for high-throughput, real-time, multicellular tracking.

## METHODS

### Materials

Polymerases, restriction enzymes and Gibson assembly mix were obtained from New England Biolabs (NEB) (Ipswich, MA, USA). Media components were obtained from BD Bioscience (Franklin Lakes, NJ, USA) and Sigma Aldrich (St. Luis, MO, USA). Oligonucleotides and synthetic DNA constructs were purchased from Integrated DNA Technologies (IDT) (Coralville, Iowa, USA). Plasmids were cloned and amplified in *E. coli* strain TG1 (Lucigen, Madison, WI, USA) or C3040 (NEB). Human urine (Catalog No: IR100007P) was purchased from Innovative Research (Novi, MI, USA). All other commercial chemical reagents were obtained from Sigma Aldrich. Bulk optical density and fluorescence measurements were made using an Infinite M200 plate reader (Tecan).

### Plasmid cloning and genomic integration in yeast

All yeast strains were derived from parental strains Fy251 [American Type Culture Collection (ATCC) 96098] or the two-hybrid strain MaV203^30^ (Invitrogen). Yeast transformations were carried out using the lithium acetate method^48^. All plasmids are derivatives of the pRS series of shuttle plasmids, cloned using standard molecular biology protocols, yeast gap repair, and Gibson assembly. Endogenous yeast promoters were obtained by PCR from genomic DNA of strain Fy251. Genomic integration in yeast was performed by homologous recombination of linearized DNA constructs with homology arms and a selectable marker. See **Supplementary Table 3** for a list of all strains used in this work. See **Supplementary Table 4** for a list of plasmids used in this work. See **Supplementary Table 5** for a list of primers used to clone endogenous promoters. See **Supplementary Table 6** for a list of DNA parts used to construct all FRAME-tags.

### FRAME-tag DNA constructs and strains

The yEGFP DNA sequence and mCherry DNA sequence were amplified from a previously constructed plasmid^24^. mTagBFP2^27^, mKO2^49^, mTurquoise2^50^, and mVenus^51^ were obtained as synthetic DNA fragments that were codon optimized for *S. cerevisiae* with the IDT codon optimization tool (Integrated DNA Technologies) and the JCat Codon Adaptation tool^52^ (see **Supplementary Table 7**). A parent dual-FP integration construct was derived from the pNH600 series of vectors^53^, which harbor integration constructs containing a multiple cloning site, an ADH1 terminator from *Candida albicans*, selectable auxotrophic markers from *Candida glabrata*, and flanking 500 bp homology regions to the target locus (pNH605: LEU2). Full integration constructs were cloned into a pRS416 backbone to allow comparison of plasmid-borne and genome-integrated constructs from the same vector. −1 PRF sequences were amplified as a DNA library from previously reported *in vitro* selection products^24^ and cloned into the parent dual-FP integration vector by gap repair in the yeast strain Fy251. Individual clones were isolated by selection on -Ura and the ratios of yEGFP and mCherry fluorescence were assayed in 96-well plates using an Infinite M200 plate reader (Tecan). Plasmid variants representing a range of fluorescence ratios were sequenced, then linearized and integrated into the LEU2 locus of a fresh Fy251 strain. Transformants were selected on synthetic dropout media (SD) (glucose, -Leu) plates, and proper integration was confirmed by sequencing of locus-specific PCR amplified DNA.

The expanded palette of dual-FP FRAME-tags was generated by individually constructing combinations of the 5 chosen frameshift motifs at early and late positions, with two fluorescent proteins (see **Supplementary Table 2**). Vectors were linearized and integrated independently into Fy251, the resulting strains were pre-cultured as above and characterized by flow cytometry. Third fluorescent proteins were screened for compatibility by cloning into a galactose inducible construct (pRS416-Gal1). Sequence verified plasmids were transformed into yEGFP-mCherry FRAME-tagged yeast strains and grown in SD(2% glucose, -Ura) or SD(2%galactose, 2% raffinose, -Ura) media. The contribution of the third fluorescent protein on both GFP and mCherry signals was evaluated by pre-culturing the strains as above and characterizing the fluorescence by flow cytometry (**Supplementary Fig. 6**). mTagBFP2 was chosen as a compatible third fluorescent protein. A three FP set of FRAME-tags was generated by individually constructing the indicated combinations of FPs and frameshift motifs (see **Supplementary Table 2**). The resulting constructs were integrated into strain Fy251, pre-cultured as above and characterized by flow cytometry.

### Flow cytometry

Characterization of the FY251-based FRAME-tag strains including the multiplex transcriptional profiling was performed on a LSR II (Becton Dickinson) using the following laser/filter sets: 488/525 for yEGFP; 594/620 for mCherry; 405/450 for mTagBFP2. For standard analysis, FRAME-tagged strains were individually pre-cultured overnight in 96-well plates in standard synthetic dropout media (SD) (2% glucose) at 30°C and 800 RPM, then inoculated at an OD_600_ of 0.1 into fresh medium as individual strains or mixtures and grown for a further 10 hours. Cells were harvested by centrifugation, kept as pellets on ice, and analyzed within two hours. High throughput characterization of the MaV203-based FRAME-tag strains, including construction and real-time tracking of the yeast co-culture, was performed on an LSR Fortessa (Becton Dickinson) using the following laser/filter sets: 488/530 for yEGFP; 561/610 for mCherry. Individual strains or mixtures of strains were cultured as described in flat-bottom 96-well plates and directly analyzed on the flow cytometer using a High Throughput Sampler (HTS, Becton Dickinson) in standard mode. All fluorescence signals were normalized by side scatter as a proxy for cell size^46^ and reported as arbitrary normalized fluorescence units scaled by 100,000. Automated gating and data analysis was carried out using custom software (FRAME-finder) based on the R package openCyto (see **Supplemental Note 2** and https://github.com/jmiguelj/FRAMEtags).

### Fluorescence microscopy

FRAME-tagged strains were grown as described above. The mixtures of FRAME-tag strains were imaged on standard microscope slides with coverslips using a Ti-E microscope with Perfect Focus System (Nikon), a CFI Plan Apochromat Lambda 20X objective and a Zyla sCMOS camera (Andor). Excitation/emission (nm) sets used were: 470/525 for yEGFP; 555/620 for mCherry. For each experiment, 10-15 fields were automatically collected representing 8,000 to 12,000 cells. Bright field and fluorescence images were sectioned with FIJI^54^ using a custom script to extract average fluorescence values of individual cells. The resulting data was input into the automated FRAME-tag gating and analysis software in R (see above) to index each cell with its respective FRAME-tag identity. These indices were used in FIJI to colorize the original bright filed images using the ROI Color Coder^55^.

### Multiplexed transcriptional profiling

Native yeast promoters were cloned upstream of an mTagBFP2 expression construct within a pRS416 plasmid backbone (see **Supplementary Table 5** and **Supplementary Table 8**). Sequence confirmed reporter plasmids were then transformed into separate FRAME-tagged Fy251 strains. Strains were individually pre-cultured overnight in 96-well plates in standard synthetic dropout media (2% glucose) at 30°C and 800 RPM. Culture density was measured using the OD_600_, and strains were combined to yield a mixed culture containing an equal proportion of all reporter-FRAME-tag strains. The reporter strain mixture was inoculated in fresh medium to an OD_600_ of 0.1, grown for 10 hours until reaching an OD_600_ of 2.7, and then inoculated into 96-well plates containing the appropriate media condition (see **Supplementary Fig. 16**) to an OD_600_ of 0.3. Cultures were incubated for 6 hours at 30°C and 800 RPM or placed statically at the indicated inducing temperature, then placed on ice and analyzed by flow cytometry within 2 hours. For each treatment, the events were partitioned with FRAME-tag gates (see above) and assigned to the corresponding promoter. The MFI for each promoter was calculated as the median value of the mTagBFP2 fluorescence. Each MFI was normalized by the geometric mean of the two internal controls^56^ (promoters from TEF1 and ACT1):

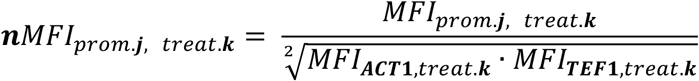

where MFI_prom.j,treat.k_ is the MFI of the j^th^ promoter in the k^th^ treatment. The expression fold-change for each sample was calculated as follows:

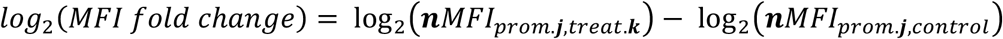

where nMFI_prom.j,control_ is the normalized MFI of the j^th^ promoter in the control treatment of standard synthetic dropout medium (2% glucose) at 30°C.

### Yeast co-culture tracking

A yeast co-culture was designed based on the MaV203 background strain, whose growth phenotype in the presence of varying concentrations of 5-FOA, histidine and 3-AT can be tuned based on Gal4 induction strength (**Supplementary Fig. 14**). This co-culture was constructed using a streamlined workflow that gives FRAME-tag indexed phenotypes with a minimal number of transformations (**Supplementary Fig. 13**). First, parent strain MaV203 was transformed with a DNA library of the 20 dual-FP FRAME-tag integrating constructs. The resulting transformants were pooled, grown overnight and transformed with a library of pRS424-SynGal4 plasmids (previously described pADH-Gal4(BD)-fs-Gal4(AD) construct library)^24^ that possess variable Gal4 transcriptional activity. Individual transformants (harboring a FRAME-tag and SynGal4 plasmid) were screened by flow cytometry to identify a set of strains such that one to three of each FRAME-tag variant was included. The histidine and 5-FOA growth phenotypes of these strains were characterized by pre-culturing overnight in standard synthetic dropout medium (2% glucose) followed by inoculation into 96-well plates containing Low His/Low 5-FOA selective media (2% glucose, 0 μM His, 5 mM 3-AT) or High His/High 5-FOA selective media (2% glucose, 130 μM His, 0.1% 5-FOA) to an OD_600_ of 0.01. Cultures were incubated at 30°C and 800 RPM for 48 hours. Nine strains were selected for the final yeast co-culture so that each member contained a unique FRAME-tag and a unique growth phenotype (see **Supplementary Table 9** for SynGal4 sequences).

For co-culture experiments, each strain was pre-cultured individually overnight in standard synthetic dropout media (2% glucose) at 30°C and 800 RPM. OD_600_ values were measured, and strains were mixed to yield a combined co-culture with an equal proportion of all strains. For the time-course assays displayed in **Supplementary Fig. 14c**, the co-culture was used to inoculate cultures to an OD_600_ of 0.01 in the following media conditions; non-selective: 130 μM His, 0 mM 3-AT, 0% 5-FOA; selection 1: 0 μM His, 5 mM 3-AT, 0% 5-FOA; selection 2: 130 μM His, 0 mM 3-AT, 0.1% 5-FOA. Co-cultures were grown at 30°C and 250 RPM and sampled at the indicated time points. The composition of the co-culture (strains A through I) was determined by flow cytometry and automatic gating using FRAME-finder, with each member’s prevalence calculated as the events assigned to that member divided the total events assigned to all members, excluding unassigned events.

To map the growth phenotype landscape of the co-culture, mixed cultures were inoculated at an OD_600_ of 0.01 into 216 individual wells (in 96-well format) containing unique growth mediums (covering a 6×6×6 matrix representing all combinations of Histidine, 3-AT and 5-FOA concentrations in a base synthetic dropout medium, see **Supplementary Fig. 15**). Cultures were incubated for 24 hours at 30°C and 800 RPM, the OD_600_ was recorded, plates were placed on ice, and cultures were analyzed by flow cytometry within two hours. Co-culture composition was quantified as above. To characterize dynamic co-culture restructuring caused by selective pressure in real time, the yeast co-culture was transitioned between pairs of conditions chosen from the phenotype map as indicated (**Fig. 3** and **Supplementary Fig. 15**). The yeast co-culture was prepared in standard synthetic dropout medium (2% glucose) as described above. Conditions used were as follows; condition #113 (blue): 0 μM His, 10 mM 3-AT, 0.02% 5-FOA); condition #200 (red): 13 μM His, 1.3 mM 3-AT, 0.08% 5-FOA; condition #30 (grey): 130 μM His, 0 mM 3-AT, 0% 5-FOA. The co-culture was inoculated into 10 mL of the initial condition at an OD_600_ of 0.01 and incubated at 30°C and 250 RPM for 24 hours. Each pre-conditioned co-culture was then pelleted (1 min, 15k RPM), washed once in 1.5 mL of the second condition medium then inoculated into 10 mL of the second condition medium at an OD_600_ of 0.03 and incubated at 30°C and 250 RPM. 100 μL samples were analyzed by flow cytometry (see above) every 20 min for a period of 13 hours.

### Statistical analysis

Overlap between mCherry fluorescence distributions (normalized to side scatter) for frameshift (*fs*) modules depicted in **Fig. 1c** and **Supplementary Fig. 2** was determined for all pairwise combinations. For each *fs* module’s distribution, *FS*, an empirical cumulative density function *CDF* was generated using R. To estimate overlap between two distributions, *FS*_*i*_ and *FS*_*j*_, the point of intersection between *CDF*_*i*_ and (1 − *CDF*_*j*_) was approximated using uniroot (in R), and the degree of overlap was taken to be twice the value of the *CDF* at the intersection point. This was performed for all *FS* pairs to generate the matrix depicted in **Supplementary Fig. 2**.

Sensitivity, specificity, positive predictive value (PPV), and negative predictive value (NPV) for FRAME-tag flow cytometry gating algorithms were determined by evaluating data from FRAME-tags captured by flow cytometry individually (~45,000 cells per FT) and combined into a dataset of all events. This allowed comparison of the automatic gating on this combined dataset relative to the known identities of the events. For a given FRAME-tag strain, *FT*_*n*_, the statistical parameters for its gate, *G*_*n*_, were derived as follows. True positives (TP): events from individual analysis of *FT*_*n*_ that fell within *G*_*n*_; false negatives (FN): events from individual analysis of *FT*_*n*_ that fell outside of *G*_*n*_; true negatives (TN): combined events excluding *FT*_*n*_ that fell outside of *G*_*n*_; false positives (FP): combined events excluding *FT*_*n*_ that fell within *G*_*n*_. Further statistics were derived for each gate as follows. Sensitivity: TP/(TP+FN); specificity: TN/(TN+FP); PPV: TP/(TP+FP); NPV: TN/(TN+FN). These parameters were determined using several different gate sets generated at varying thresholds of sensitivity, i.e. the percentage of total cells captured by gates (see **Supplementary Fig. 9**).

Mock dilution studies of the gating pipeline were performed for each FRAME-tag strain by computationally constructing a new distribution using random subsampling from the original distribution, recombining with the other undiluted strain data and reapplying the automated gating pipeline, followed by the above statistical analyses (see **Supplementary Fig. 10**).

## DATA AVAILABILITY

Additional data that support the findings of this study are available upon request. The software and resources for the automated FRAME-tag gating and analysis pipeline (FRAME-finder) can be accessed at: https://github.com/jmiguelj/FRAMEtags

## ACKNOWLEDGMENTS

We would like to thank Tal Danino for generously providing access to his fluorescent microscope, and Zakary Singer for assistance with fluorescence microscopy experiments. We also thank Harris Wang for providing access to his flow cytometer and for helpful comments on this manuscript. This work was supported by funds from the US National Institutes of Health (NIH) (R01 AI110794 to V.W.C.). A.V.A. was supported by a US National Institutes of Health F30 fellowship (F30CA174357). M.J. was supported by a National Science Foundation Graduate Research Fellowship (DGE-11-44155). Research reported in this publication was performed in the Columbia Center for Translational Immunology Flow Cytometry Core, supported in part by US National Institutes of Health award S10RR027050 and the Columbia University Microbiology and Immunology Flow Cytometry Core.

## AUTHOR INFORMATION

### Contributions

A.V.A. and M.J. conceived the project, designed and performed experiments, and analyzed data. A.V.A. developed the frameshift motif library. M.J. developed the flow cytometry and microscopy analysis code (FRAME-finder). A.V.A, M.J., and V.W.C. wrote the manuscript. V.W.C. supervised the research.

