## Supplementary Notes, Figures and Tables for "FRAME-tags: genetically encoded fluorescent markers for multiplexed barcoding and time-resolved tracking of live cells"

**Supplementary Materials Contents**

**Supplementary Note 1**. Theoretical space of a three-color FRAME-tag palette
**Supplementary Note 2**. FRAME-finder: Design and implementation of the automated gating software for FRAME-tags

**Supplementary Figure 1**. Frameshift module architecture and predicted RNA motif structures
**Supplementary Figure 2**. Characterization and selection of frameshift motifs
**Supplementary Figure 3**. FRAME-tags enhanced by chromosomal integration
**Supplementary Figure 4**. Visual index of FRAME-tags
**Supplementary Figure 5**. Secondary effects of frameshift motifs on protein expression levels
**Supplementary Figure 6**. Identification of a compatible third fluorescent protein
**Supplementary Figure 7**. FRAME-tags in RGB space
**Supplementary Figure 8**. Overview of the FRAME-finder automated gating pipeline
**Supplementary Figure 9**. Statistical analysis of the FRAME-finder automated gating pipeline
**Supplementary Figure 10**. Automated gating detection limit of dilute populations
**Supplementary Figure 11**. Fluorescence microscopy with 8 FRAME-tags
**Supplementary Figure 12**. Fluorescence microscopy with 20 FRAME-tags
**Supplementary Figure 13**. Optimized workflow for tagging phenotypes in new strains
**Supplementary Figure 14**. A multi-strain co-culture for dynamic population tracking studies
**Supplementary Figure 15**. High throughput condition profiling of a yeast co-culture
**Supplementary Figure 16**. Multiplexed expression analysis histograms

**Supplementary Table 1**. Sequences of frameshift motifs
**Supplementary Table 2**. FRAME-tag constructs
**Supplementary Table 3**. Strains used in this study
**Supplementary Table 4**. Plasmids used in this study
**Supplementary Table 5**. Primers for promoter cloning
**Supplementary Table 6**. DNA sequences of FRAME-tag parts
**Supplementary Table 7.** DNA sequences of additional fluorescent proteins
**Supplementary Table 8**. DNA sequences of promoters for multiplexed transcriptional profiling
**Supplementary Table 9**. DNA sequences of SynGal4 constructs for co-culture tracking

**Supplementary Data 1**. Example raw flow cytometry data of FRAME-tagged strains.
**Supplementary Data 2**. Example raw microscopy data of FRAM-tagged strains.
**Supplementary Data 3**. Raw data of frameshift efficiency and percent overlap.
**Supplementary Data 4**. Calculated Sensitivity, Specificity, PPV, NPV values of automated gating.
**Supplementary Data 5**. Calculated PPV values of automated gating of mock diluted strains.

**Supplementary References**

**Supplementary Note 1**. Theoretical space of a three-color FRAME-tag palette.

Extrapolating from the FRAME-tags (FTs) that were constructed and tested empirically, we can estimate the expected number of resolvable FT variants in three-color space. Some of the FT constructs would likely require -1 PRF signals that were not implemented for the main two-color system in order to fit additional FTs that produce the desired absolute expression level and ratios of FPs for the three-color system. First, in the vicinity of FT-13, there should be sufficient room to add other FTs, or replace FT-13 with two new FTs. Here, we make the reasonable assumption that such an exchange can be made with available *fs* modules and that an additional FT can be added to the existing two-FP arrangement. Also, we will assume that three-color FP variants will behave in a similar manner to the two-color constructs in terms of absolute protein expression and population distributions, as this was observed for the 3-FP constructs generated.

As shown in **Figure 2b** and **Supplementary Figure S7**, the three-color space can be represented as a three-dimensional cube that contains the FRAME-tag palette. Again, we assume that an additional FT can be added to the two-color set to create a palette of 21 FTs as shown below (replacing FT-13 with FT-13A and FT-13B, red circles). Because of differences in resolution between FTs that have high vs. low overall FP expression, we cannot simply scale in a new dimension by multiplying the existing set by a constant. Therefore, we assume scalability in the third color dimension follows rules of scalability in the two-dimension case. Constructs that express at least one FP at the 30% or 100% absolute level are capable of being scaled across the second color dimension so that a total of 5 FTs is created. Among the set of 21 hypothetical 2-dimensional FTs, the 16 shown in the orange zone (FT-1, FT-2, FT-3, FT-4, FT-5, FT-6, FT-7, FT-8, FT-9, FT-10, FT-11, FT-12, FT-16, FT-17, FT-20, and FT-21) satisfy this condition. So, we predict that each of these 16 FTs can generate 4 additional resolvable FTs each using the new color dimension for a total of 80 three-color FTs (16x5). FTs that do not express at least one FP at the 30% or 100% level (FT-13A, FT-13B, FT-15, FT-19, FT-22) shown in the blue zone can be scaled in the third color dimension to create only 3 additional resolvable FTs each, thus generating a total of 20 total FTs (5x4). Summing these, we predict that a total of ~100 FTs can be generated based on this simple scaling analysis.

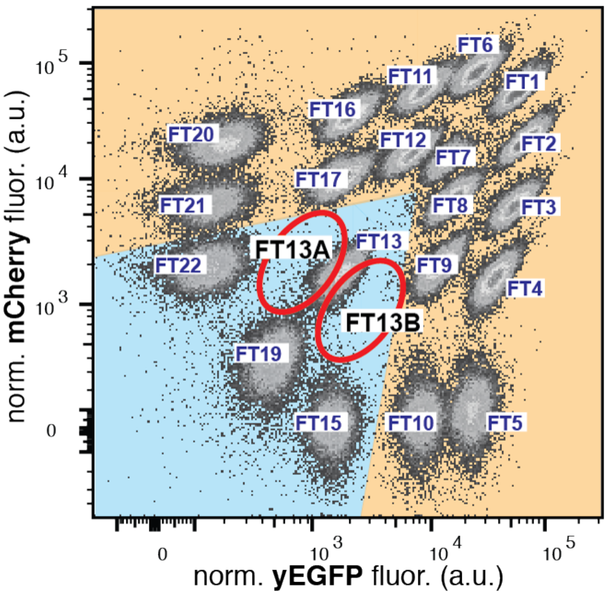

**Supplementary Note 2.** FRAME-finder: Design and implementation of the automated gating software for FRAME-tags.

While the raw cytometry data generated by flow cytometry or microscopy using FRAME-tagged cells (FTs) can be analyzed manually using available software, this process is tedious for samples that include many FT populations and, as with any manually analyzed cytometry data, it is subjective. Therefore, to simplify analysis of FTs and to remove user bias, we wrote FRAME-finder, a custom R package (https://github.com/jmiguelj/FRAMEtags) that functions as a front end for the existing suite of automated flow cytometry processing functions implemented by the R package openCyto.[1]

The existing openCyto pipeline enables automated gating of cytometry data based on a user-defined input file (the "Gating Template") that defines a gating hierarchy and the corresponding automatic gating functions to use at each level of the hierarchy. The FRAME-finder code builds on this pipeline in three ways: (1) it dynamically generates the correct "Gating Template" based on a much simpler input file that need only list the FTs present in the sample; (2) it automatically pre-processes the raw data to generate two derived parameters used in gating (RFP/GFP ratio and RFP+GFP total fluorescence); and (3) it includes an automatic gating function that simplifies valley-specific bisection of 1-dimensional histograms with multiple sharp peaks and valleys characteristic of the FT populations.

The FRAME-finder analysis code also enables robust and efficient assignment of FT indices to each event even for high-throughput and real-time experiments that can generate hundreds of individual data files. Available unsupervised 2-dimensional clustering functions, while powerful and agnostic to cluster structure, were found to be either computationally intensive or unable to consistently assign cluster identities from a predetermined list. To overcome this, the FRAME-finder code leverages the consistent cluster structure of FTs. This cluster structure can be completely predetermined based on the known FTs in the sample and therefore efficiently traversed computationally.

An illustrative example of this automated process is included in **Supplementary Figure 8**. FRAMEtags software and resources for analysis can be accessed at: https://github.com/jmiguelj/FRAMEtags.

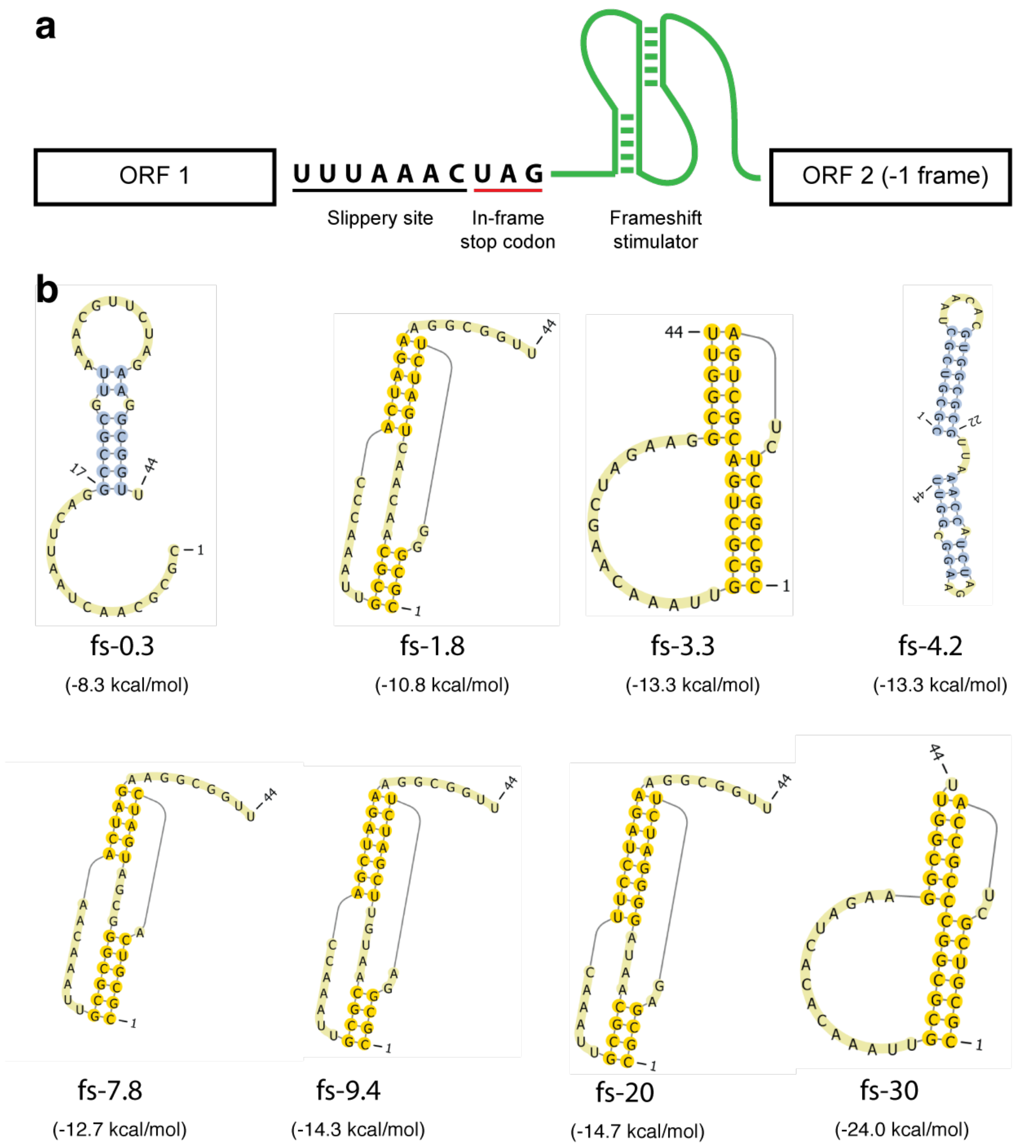


**Supplementary Figure 1**. Frameshift module architecture and predicted RNA motif structures. (**a**) Frameshift module architecture. Open reading frames (ORFs) are separated by frameshift modules that contain a heptanucleotide slippery sequence, an in frame amber stop codon, and a downstream RNA stimulatory element. Frameshifting at the slippery site is stimulated by adjacent RNA secondary structures in *cis*, such as hairpins and pseudoknots (sown in **b**), which divert translation from the original reading frame containing a stop codon (U-UUA-AAC-UAG…) to the -1 reading frame (UUU-AAA-CUA-G...) containing ORF2. (**b**) Frameshift stimulatory structure prediction. Minimum free energy (mfe) RNA secondary structures were predicted computationally with pKiss[2] using the 'mfe' function for non-pseudoknotted structures and the 'local' function for pseudoknotted structures. For each *fs* module, sequences of length 44, starting 8 nucleotides downstream of the slippery site, were evaluated for secondary structure. All calculations were performed at 37 ˚C with Turner 2004 default parameters in mode [P] to ignore kissing hairpins (K-type pseudoknots). Secondary structure diagrams are shown for the mfe of each *fs* module. Structure graphics were generated using PseudoViewer3 (http://pseudoviewer.inha.ac.kr/).

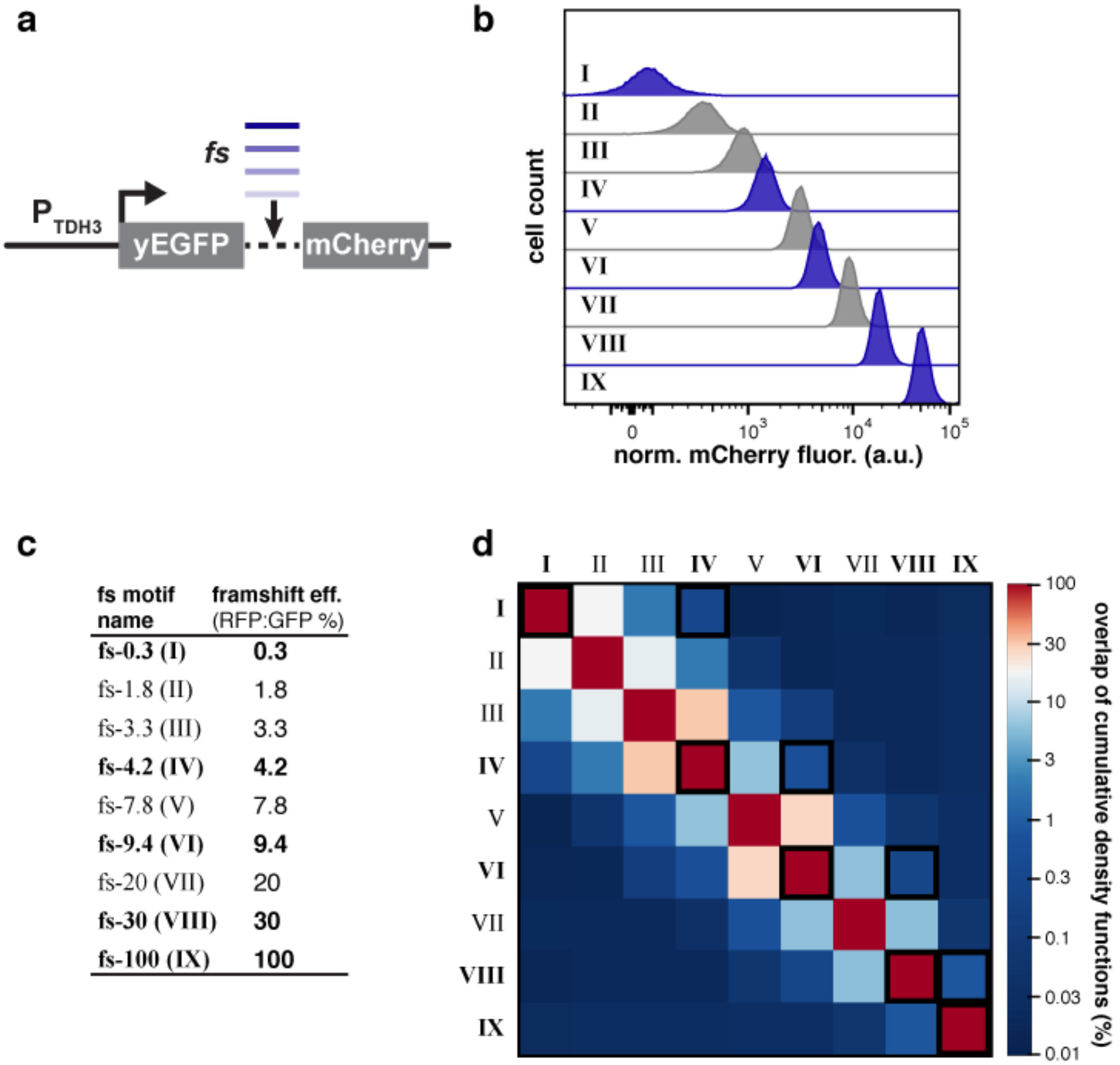


| fs motif name | framshift eff. (RFP:GFP %) |
|---|---|
| **fs-0.3 (I)** | **0.3** |
| fs-1.8 (II) | 1.8 |
| fs-3.3 (III) | 3.3 |
| **fs-4.2 (IV)** | **4.2** |
| fs-7.8 (V) | 7.8 |
| **fs-9.4 (VI)** | **9.4** |
| fs-20 (VII) | 20 |
| **fs-30 (VIII)** | **30** |
| **fs-100 (IX)** | **100** |

**Supplementary Figure 2**. Characterization and selection of frameshift motifs. (**a**) Frameshift (*fs*) motifs were characterized in a dual-fluorescent protein reporter construct. (**b**) Nine distinct frameshift modules were characterized with the dual-fluorescent protein reporter, chromosomally integrated in yeast, and analyzed by flow cytometry. Modules that were selected for FRAME-tags are shaded in blue. mCherry fluorescence was normalized by side scatter. (**c**) Each frameshift motif resulted in a characteristic frameshift efficiency that leads to a unique stoichiometry between mCherry and yEGFP. Motifs selected for FRAME-tags are bolded. Frameshifting efficiency was defined by the ratio of the mCherry to yEGFP fluorescence, by first subtracting the baseline fluorescence of a non-fluorescent strain (FY251) and defining fs-100 (non-frameshift fusion protein) as 100% efficiency. **(d)**. The final set of *fs* motifs was selected by choosing a set with mutually minimal overlaps in their empirical cumulative density functions noted by the squares outlined in black (see Methods for overlap calculation). Frameshift efficiency calculations and raw heatmap data are included in **Supplementary Data 3**.

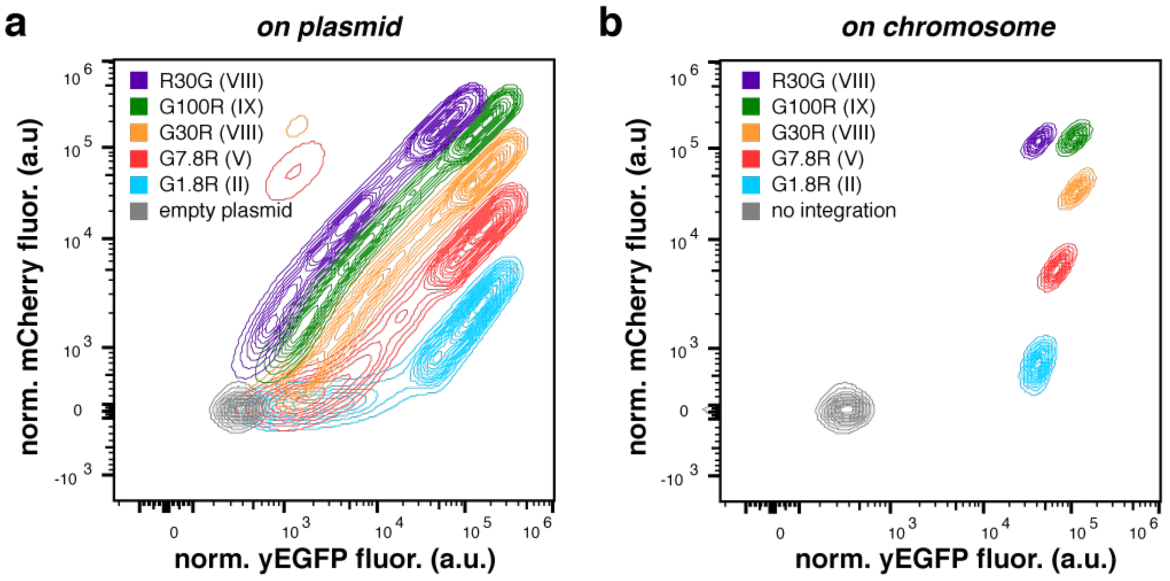


**Supplementary Figure 3**. FRAME-tags enhanced by chromosomal integration. Five FRAME-tags were compared by flow cytometry when (**a**) expressed from a CEN plasmid (low copy) or (**b**) expressed from the chromosome after single-copy integration at the *Leu2* locus. Individual samples overlaid and identified by color. Fluorescence was normalized by side scatter; contours represent 5% quantiles.

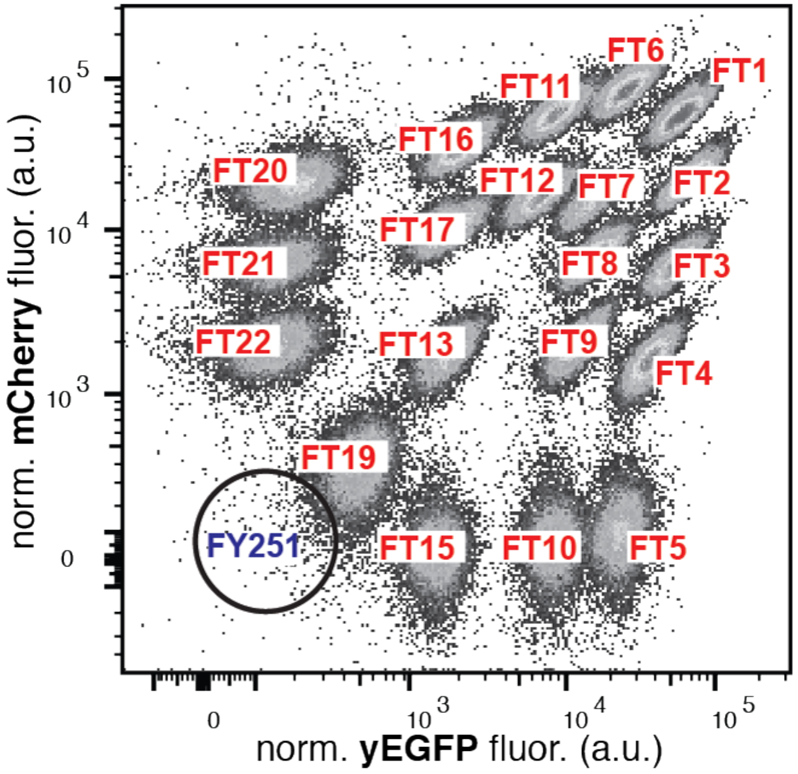


**Supplementary Figure 4**. Visual index of FRAME-tags. The corresponding strain identity of each population was validated with single strain experiments and the FT# names used throughout are labeled in red. The approximate location of a non-fluorescent strain (FY251) is noted by the black outlined circle.

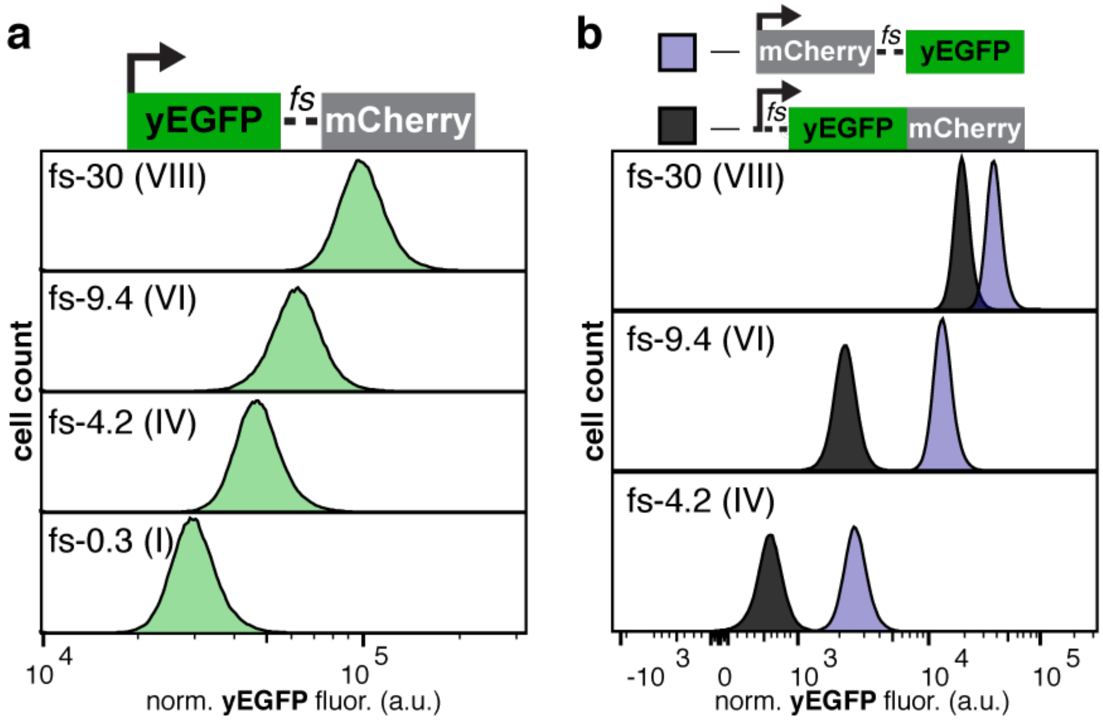


**Supplementary Figure 5**. Secondary effects of frameshift motifs on protein expression levels. (**a**) Fluorescence distributions of yEGFP show that frameshift motifs (*fs*) also influence the expression of the upstream open reading frame. Efficiency of *fs* as in **Supplementary Figure 2.** (**b**) Comparison of yEGFP fluorescence controlled by a late *fs* module (purple) or an early *fs* module (black). The effective yEGFP expression from the same *fs* module depends on its position within the construct (i.e., late or early). The same *fs* motifs are used as in panel **a**, identified by their efficiency.

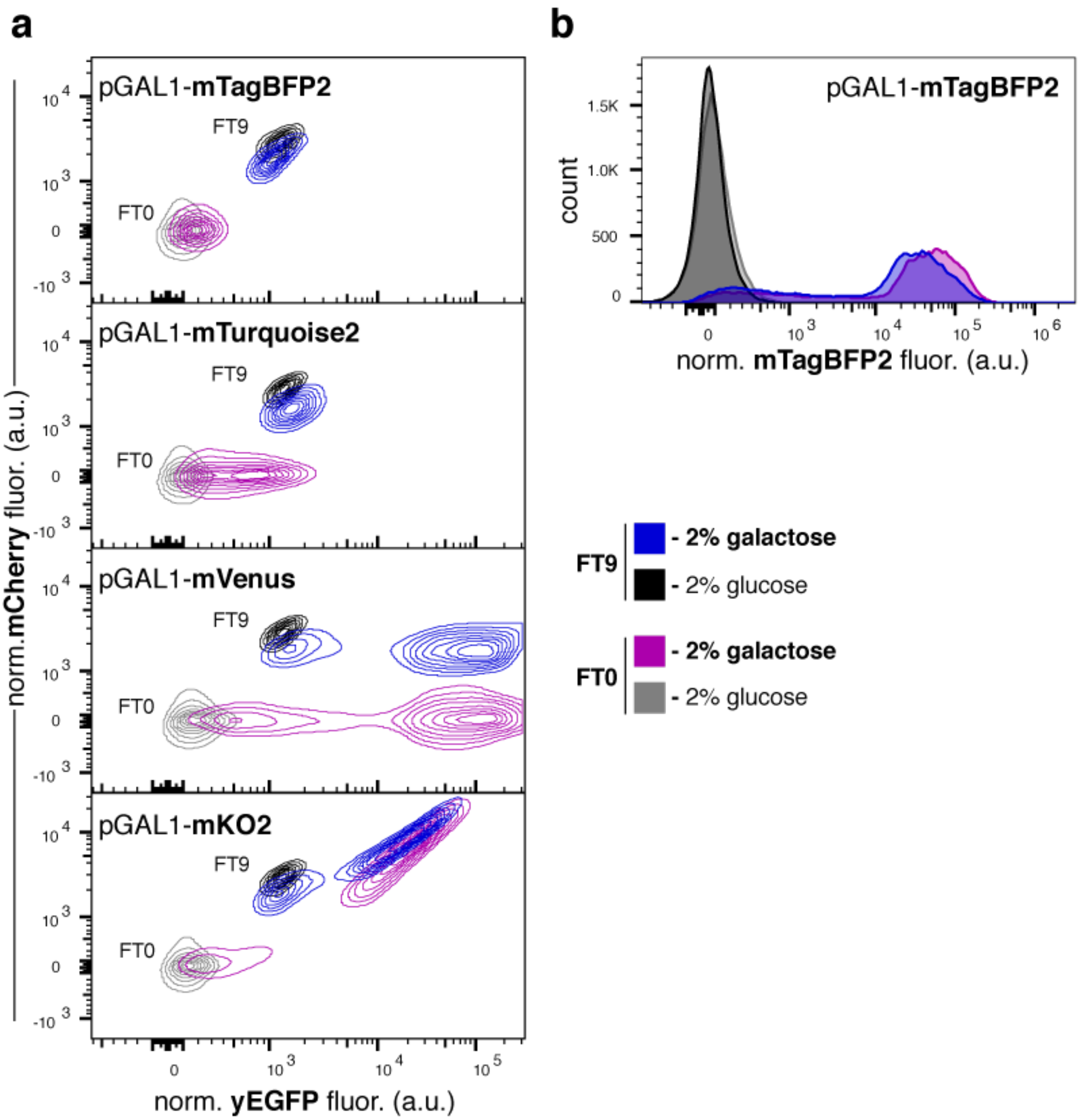


**Supplementary Figure 6**. Identification of a compatible third fluorescent protein. (**a**) yEGFP and mCherry flow cytometry 10% contour plots of FRAME-tagged strains expressing a galactose-inducible reporter. Strains tagged with either FT9 (blue, black) or no tag (FT0, purple, grey) were transformed with a plasmid carrying a third fluorescent protein (mTagBFP2, mTurquoise2, mVenus or mKO2) under the control of a galactose inducible promoter (pGAL1). These strains were grown in either standard synthetic dropout medium (2% glucose, black and grey) or 2% galactose medium (blue, purple). The yEGFP and mCherry signal of the tags was not affected by expression of mTagBFP2 construct (colors v greys). (**b**) Histogram of the of the mTagBFP2 signal from the pGAL1- mTagBFP2 strains in panel **a**. Baseline and induced of expression of mTagBFP2 is not influenced by the presence of a yEGFP/mCherry signal (FT9 v. FT0).

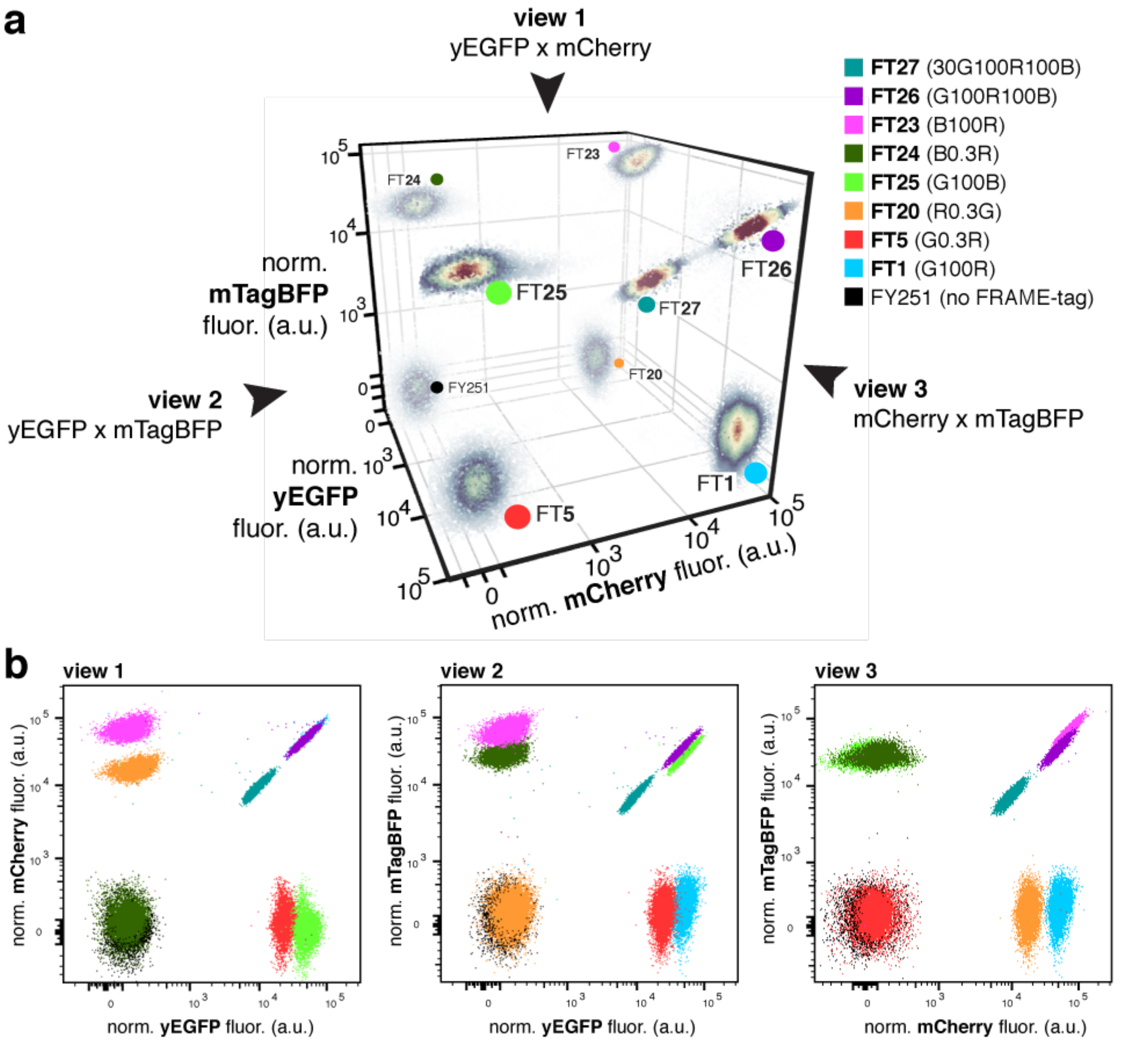


**Supplementary Figure 7**. FRAME-tags in RGB space. **(a)** By including just one additional FP (mTagBFP2) the full combinatorial RGB space becomes available, and up to 100 FRAME-tags could be accessed (see **Supplementary Note 1**). Here, we show three additional dual-FP FRAME-tags that use the blue dimension (FT23, FT24, FT25), and two triple-FP FRAME-tags that use all three dimensions of color (FT26, FT27). Scatterplot of a mixed sample including all noted strains, plotted in 3D using Plotly (https://plot.ly). Events shaded by density (low=dark blue, high=dark red) and populations identified from single strain experiments labeled with corresponding strain identity and a colored circle according to the legend. Legend includes both FT and FP nomenclature from **Supplementary Table 2**. Transparency added to some events to clarify perspective. **(b)** Single strain experiments overlaid and identified by color as in panel **a**. Views correspond to those noted in panel **a**. Fluorescence normalized by side scatter.

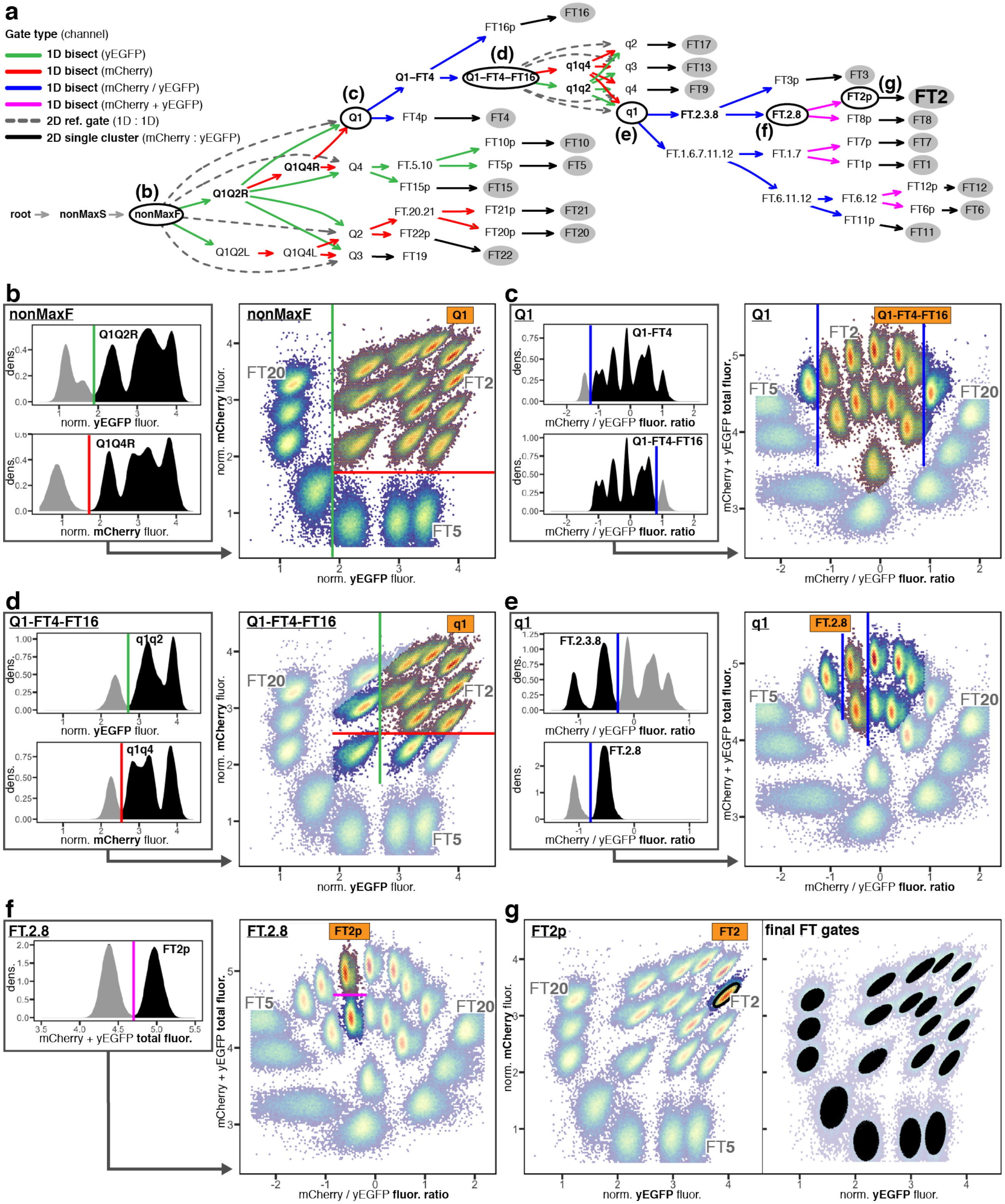


**Supplementary Figure 8.** Overview of the FRAME-finder automated gating pipeline. **(a)** Example gating hierarchy dynamically generated with the FRAME-finder package and implemented with openCyto[1] (https://github.com/jmiguelj/FRAMEtags). This approach primarily uses fast 1D bisecting gates along four derived parameters: normalized yEGFP (green), normalized mCherry (red), fluorescence ratio (blue) and total fluorescence (pink). The final FT populations are gated using a fast 2D single cluster gate (black) that allows specifying the desired sensitivity by choice of the percentile captured. Parent populations illustrated in panels **b-g** are in black outlined ovals and final FT populations in grey ovals. **(b-g)** Automated hierarchical isolation of FT2. Gate colors as in panel **a**. In each panel, left insets show the 1D histogram bisecting gates, and the scatter plots show the resulting gated subpopulation (orange highlight and label). Parent population events highlighted in bright colors (names underlined) and overlaid on all original events in faded colors. Right plot in panel **g**, shows all resulting FT gates derived in the same manner illustrated for the FT2 gate. Populations corresponding to FT2, FT5 and FT20 labeled for reference. Normalized fluorescence values plotted on a logicle scale. Fluorescence ratio and total fluorescence values plotted using arbitrary log units. The raw flow cytometry data of this representative example is included in **Supplementary Data 1**, pre-gated to remove extreme scatter and fluorescent values (nonMaxF) and including the derived parameters (normalized GFP, normalized RFP, fluorescence ratio, fluorescence total). The raw data from a representative microscopy experiment is also included in **Supplementary Data 2**, including sectioned cell areas and ferret in pixel units (1 px = 0.33μm)

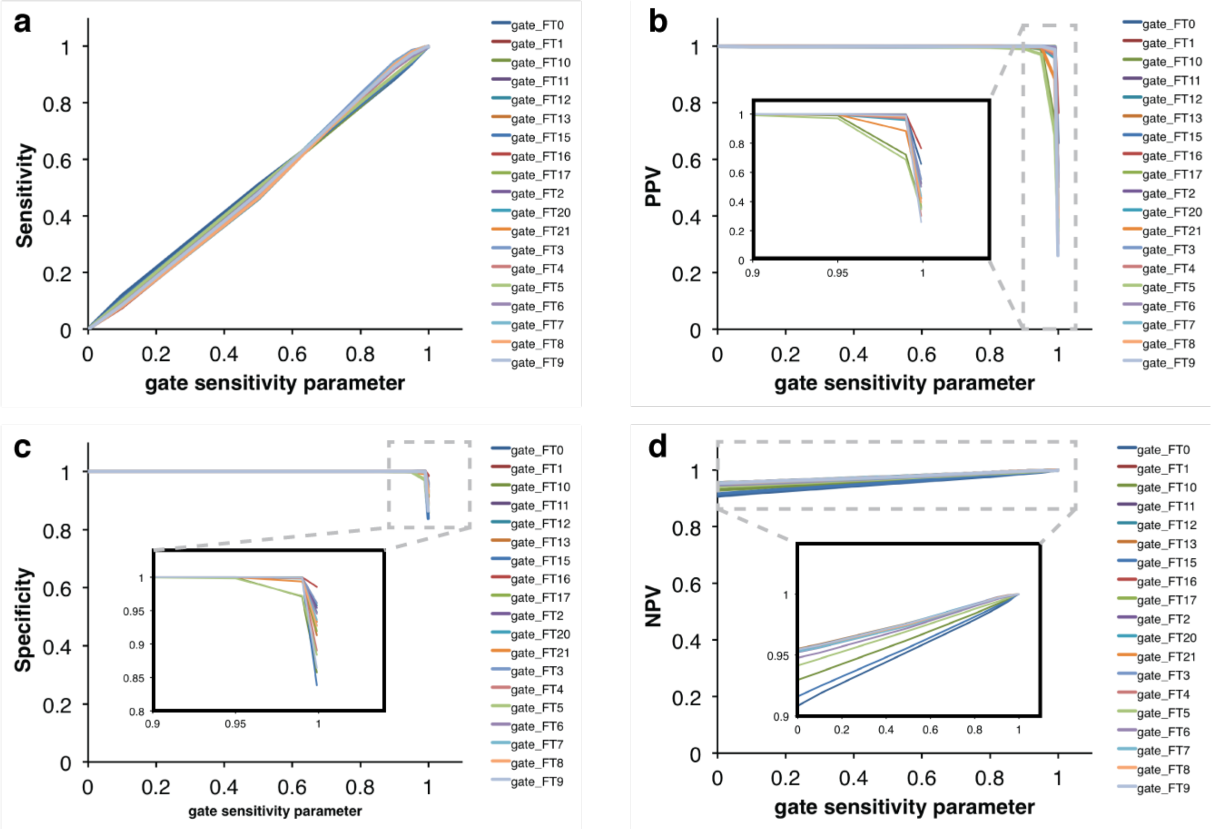


**Supplementary Figure 9**. Statistical analysis of the FRAME-finder automated gating pipeline. (**a**) Sensitivity, (**b**) positive predictive value (PPV), (**c**) specificity, and (**d**) negative predictive value (NPV) were calculated from gating data produced at various gate sensitivity parameters (0.001, 0.01, 0.1, 0.5, 0.8, 0.9, 0.95, 0.99, and 0.999) corresponding to the percentile of cells captured in the final FT gate relative to its immediate parent population (e.g. percentile of events in the FT2 gate relative to events in the FP2p gate, see **Supplementary Figure 8**, black gates). Statistics are shown plotted vs. this gating sensitivity parameter. For details on statistical calculations, see Methods section under the Statistical analysis heading. See **Supplementary Data 4** for the corresponding numeric values.

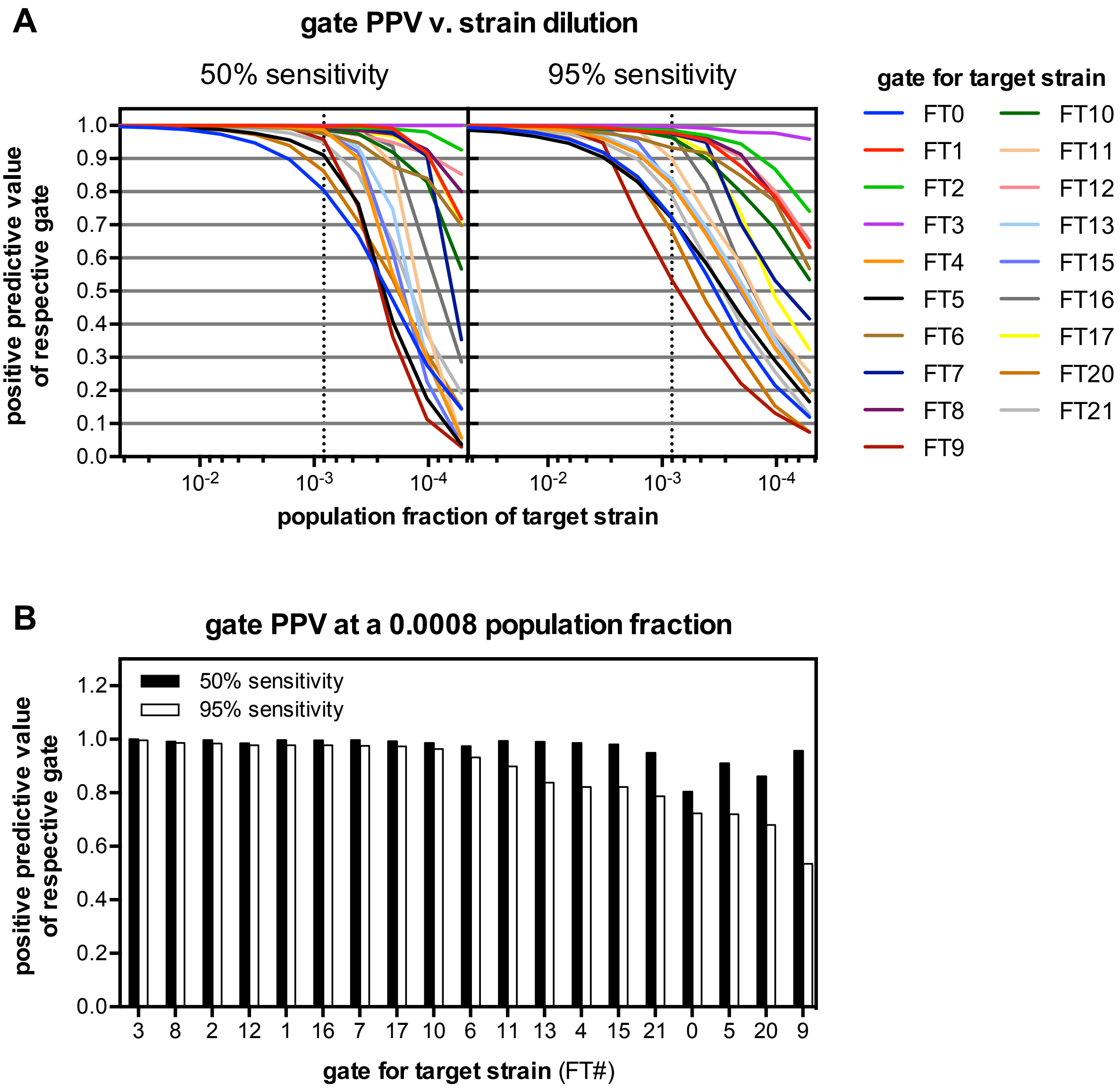


**Supplementary Figure 10**. Automated gating detection limit of dilute populations. **(a)** Flow cytometry experiments for all single strains were run independently and programmatically combined into a mock mixed dataset. This allowed generation of dilution series for each strain within this mixed dataset through subsampling of the events known to belong to that strain, while maintaining all other events unchanged. For each new dilution dataset, the automated gating pipeline was run and the positive predictive values (PPV) of the resulting gates were calculated. This was repeated at a gating sensitivity parameter of 50% and 95% which corresponds directly to real statistical sensitivity (see **Supplementary Figure 9a**). **(b)** Data from panel **a** (at dashed lines), plotted at a single dilution of 0.0008 population fraction of the target strain to be detected, shows that a majority of strains can be correctly gated by the software at this dilution to give a PPV above 0.9 with a sensitivity of 50%. See **Supplementary Data 5** for the corresponding numeric values.

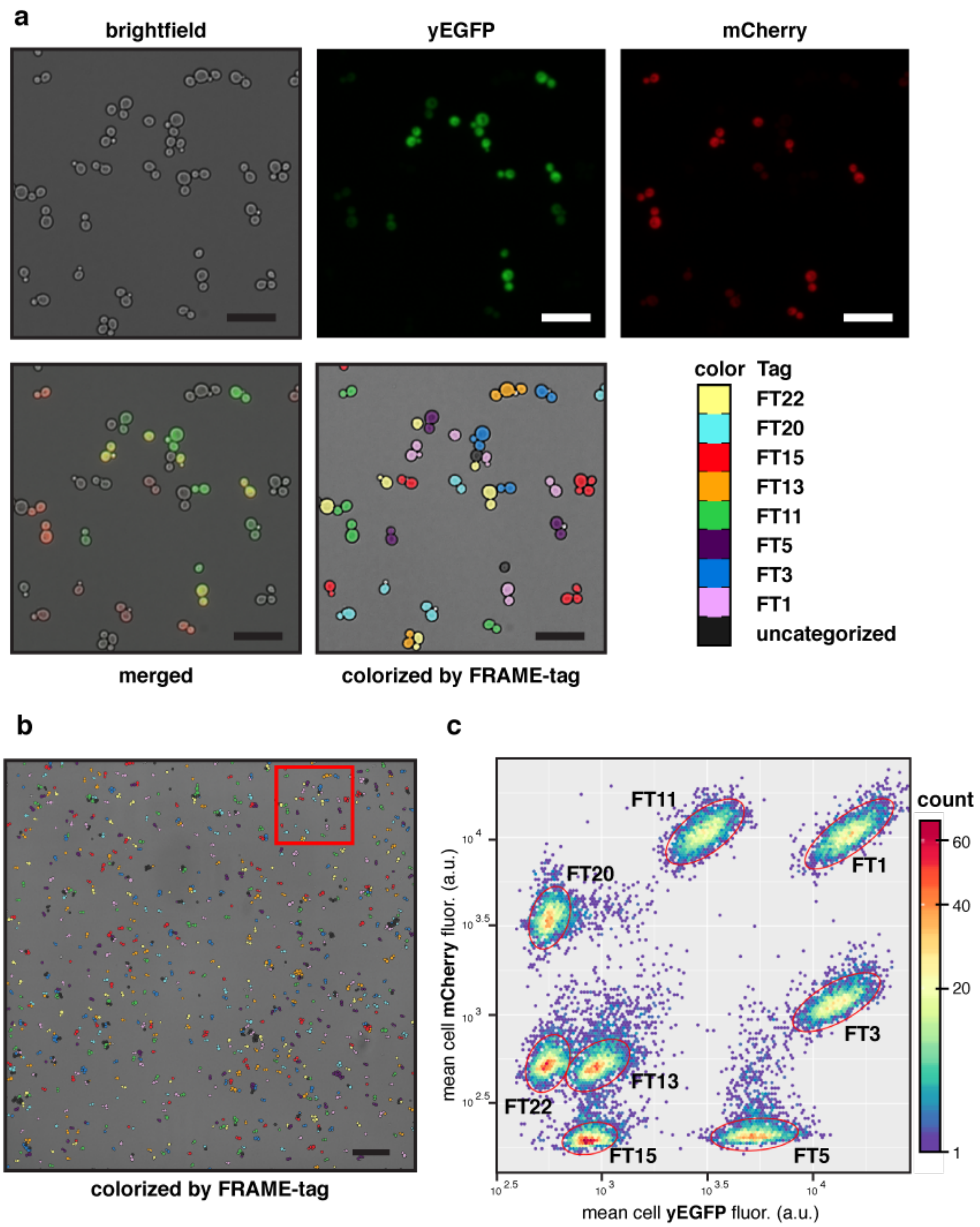


**Supplementary Figure 11**. Fluorescence microscopy with 8 FRAME-tags. (**a**) FRAME-tags imaged by brightfield, yEGFP, and mCherry with merged images. After the analysis as described in the Methods and identified using the automated gating pipeline (**Supplementary Fig. 8**), yeast cells can be false colored. (**b**) One full wide-view image of colorized yeast. (**c**) Grouping and analysis of individual FRAME-tags based on fluorescence microscopy, representing the composite data from 10 fields as shown in panel **b** and used for colorization.

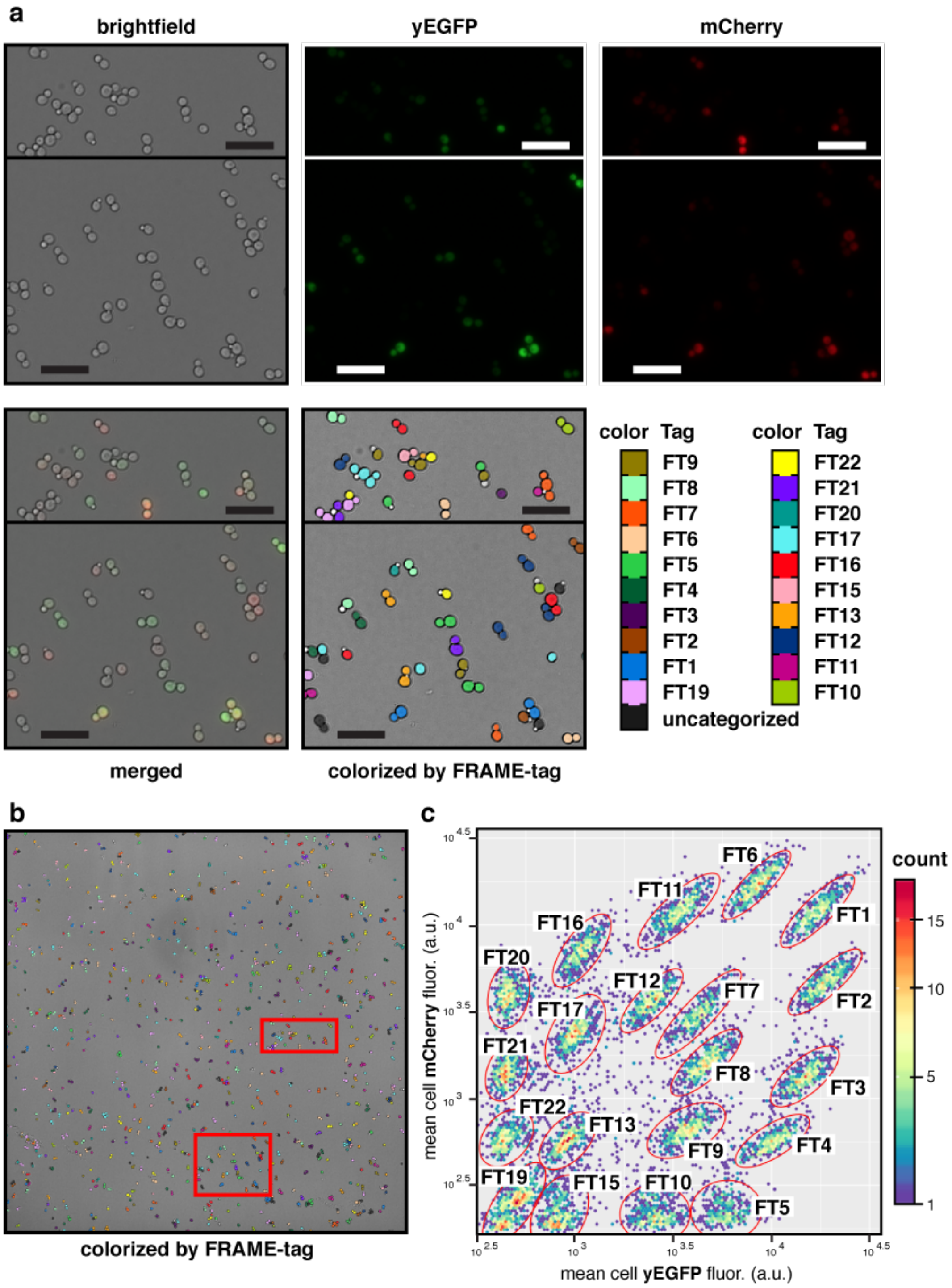


**Supplementary Figure 12**. Fluorescence microscopy with 20 FRAME-tags. (**a**) FRAME-tags imaged by brightfield, yEGFP, and mCherry with merged images. After the analysis as described in the Methods and identified using the automated gating pipeline (**Supplementary Fig. 8**), yeast cells can be false colored. (**b**) One full wide-view image of colorized yeast. (**c**) Grouping and analysis of individual FRAME-tags based on fluorescence microscopy, representing the composite data from 10 fields as shown in panel **b** and used for colorization.

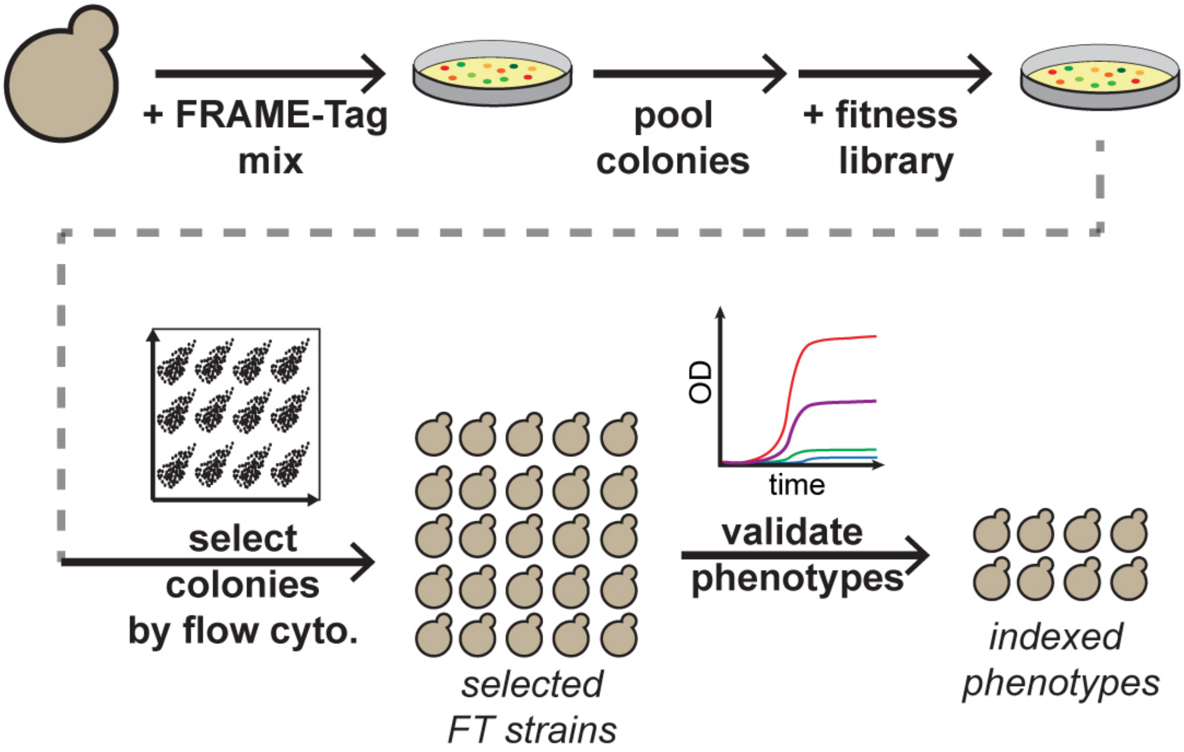


**Supplementary Figure 13**. Optimized workflow for tagging phenotypes in new strains. This workflow is designed to minimize DNA transformation steps. The mixed FRAME-tag DNA constructs can be co-transformed into the desired background strain. Barcoded transformants are directly pooled without tag identification, followed by transformation of pooled constructs or pre-screened libraries of constructs imparting a desired phenotype into the FRAME-tag strain mixture. Analysis is performed on individual colonies to first identify a set of strains with overrepresented FRAME-tag identities. The phenotypes of all members within this overrepresented set are identified and a subset of uniquely indexed phenotypes can be selected and used. Alternatively, pooled FRAME-tag DNA constructs can also be integrated into a preexisting pooled library of phenotypically variable strains followed by phenotype indexing as described.

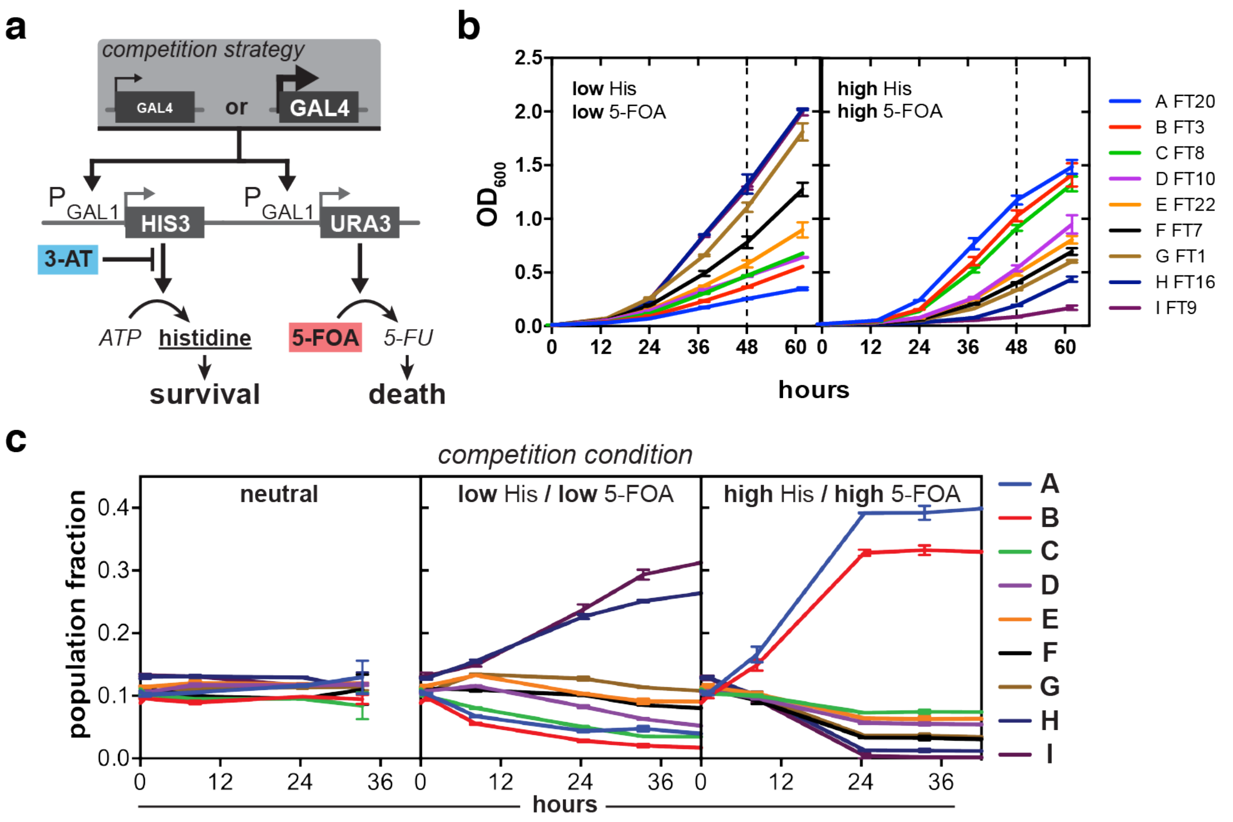


**Supplementary Figure 14**. A multi-strain co-culture for dynamic population tracking studies. (**a**) FRAME-tags were introduced into a background strain of MaV203 (see Methods). The constructs were designed such that growth would be dependent on two factors within the host strain, namely His3 and Ura3 expression. Growth selection was modified with exogenously added factors, namely histidine, 3-aminotriazole (3-AT), and 5-fluoroorotic acid (5-FOA). His3 and Ura3 production are dependent on Gal4 transcriptional activity. High production of His3 allows for cells to grow in the absence of histidine and the presence of 3-AT. High Ura3 expression causes production of the antimetabolite 5-FU in the presence of 5-FOA, which inhibits growth. To impart each strain with a different level of Gal4 activity, previously developed ratiometric Gal4 constructs[3] containing frameshift modules between the N-terminal DNA binding domain and the C-terminal activation domain were implemented (see **Supplementary Table 4** and **Supplementary Table 9**). Overall Gal4 activity is a function of frameshift efficiency. Frameshift sequences were screened in a library fashion on plasmids. Plasmids with desirable phenotypes were transformed into pooled FRAME-tag strains as described in **Supplementary Figure 13**. **(b)** A set of strains with uniquely indexed growth phenotypes was selected and characterized in single-strain experiments. $OD_{600}$ values were measured over 60 hours at 30 °C and 800 RPM; error bars indicate s.d. from 2 replicates. Low His/Low 5-FOA = 2% glucose, -His, 5 mM 3-AT; High His/High 5-FOA = 2% glucose, 130 µM His, 0.1% 5-FOA. Dashed lines mark data used in **Fig. 4a**. (**c**) Time courses of the community grown in non-selective media, or Low His/Low 5-FOA (2% glucose, -His, 5 mM 3-AT) or High His/High 5-FOA (2% glucose, 130 µM His, 0.1% 5-FOA), analyzed by flow cytometry using FRAME-tags to determine the respective population fraction of each community member; error bars indicate s.e.m. from 3 replicates.

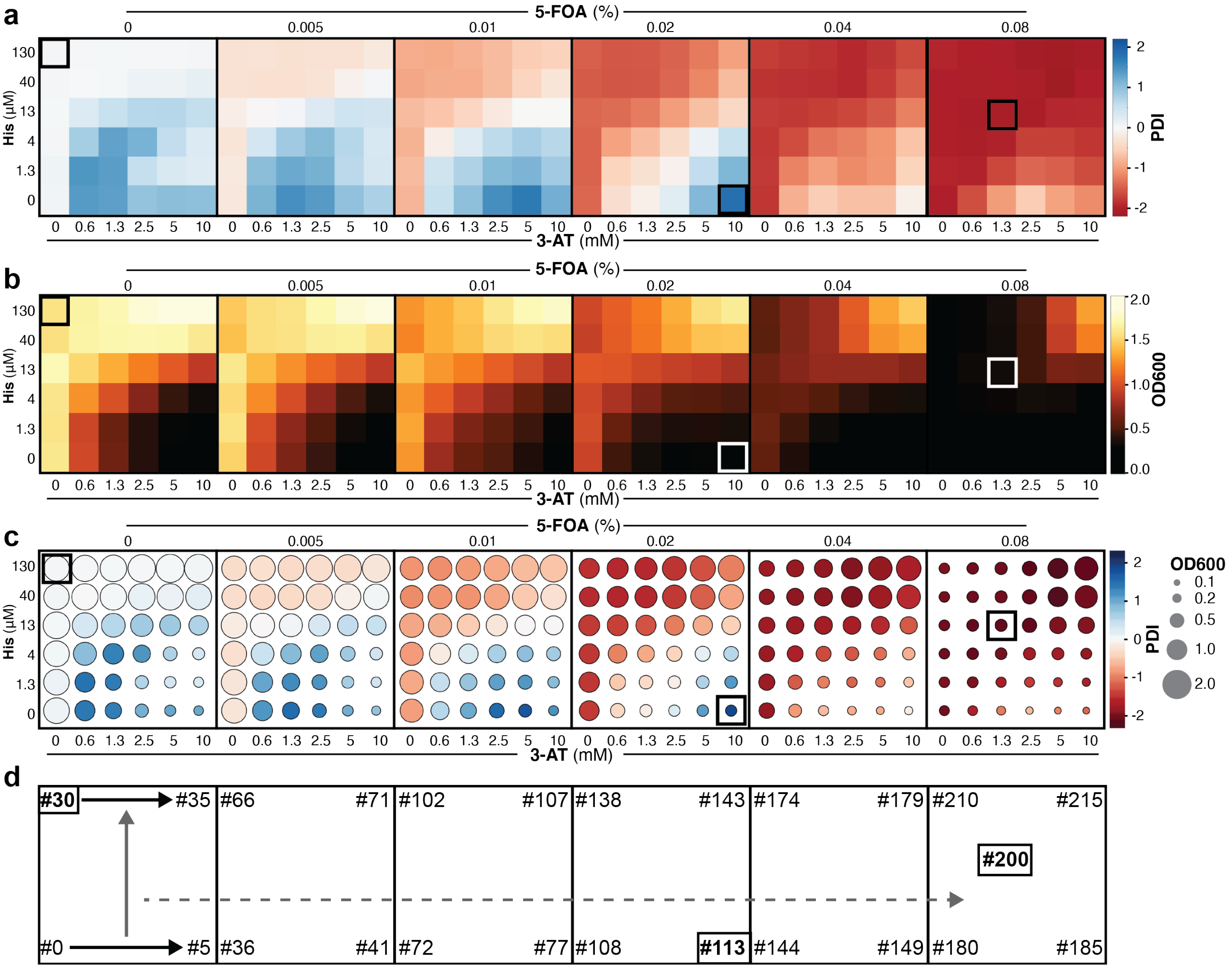


**Supplementary Figure 15**. High throughput condition profiling of a yeast co-culture. **(a-c)** The co-culture composition was assessed after 24 hours of growth in 216 conditions that varied in concentrations of histidine (His), 5-fluoroorotic acid (5-FOA), and 3-aminotriazole (3-AT). FRAME-tags were used to determine all individual strain population fractions (See Methods for experimental details). For visual representation of the selective pressure imparted by conditions, we calculated the Population Distribution Index (PDI) for each condition. The PDI is simply a weighted average (based on population fraction) of assigned phenotype scores that rank order the strains by increasing Gal4, His3 and Ura3 expression that leads to decreasing fitness in the High His/High 5-FOA standard selection condition. PDI can be formally defined by first defining the set containing all nine strains as $S = \{A, B, C, D, E, F, G, H, I\}$, the set $X = \{-4, -3, -2, -1, 0, 1, 2, 3, 4\}$, and a function $f$ that maps $S$ to $X$ such that $\{A, B, \ldots I\} \rightarrow \{-4, -3, \ldots 4\}$. The PDI was determined from the culture composition according to the formula below:

$$PDI = \sum_{s \in S} X_s \cdot P_s$$

where $X_s$ is the mapped integer value for $S_s$ and $P_s$ is the measured population fraction of strain $s$. PDI takes values between -4 and 4. **(a)** Results of 216 growth conditions plotted as a heatmap of

PDI, with red tiles indicating low PDI values and blue tiles indicating high PDI values. In general, low PDI (red) indicate conditions that favor strains with low Gal4 expression, while high PDI (blue) indicate conditions that favor strains with high Gal4 expression. **(b)** Absolute growth for the entire culture (all strains) in each of the 216 growth conditions, represented as a heat map with lighter color representing higher measured $OD_{600}$. **(c)** Combined data from panels **a** and **b** showing overall community state map. Color represents PDI, as in the top panel; circle diameter represents growth and is proportional to the measured final $OD_{600}$ of the bulk culture in that condition. **(d)** Conditions were numbered from 0 to 215 as shown starting from the lower left corner of the array and continuing as shown by the black then grey then dashed arrows. Conditions used in **Fig. 4d** are labeled in all panes with square outlines.

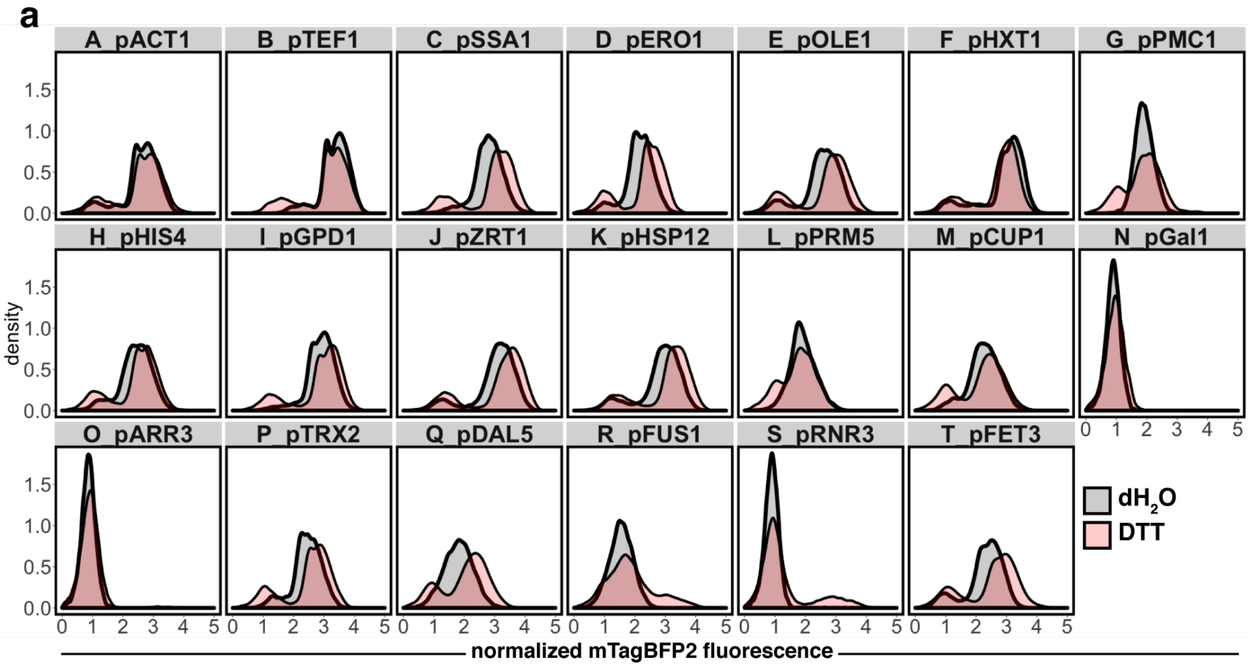


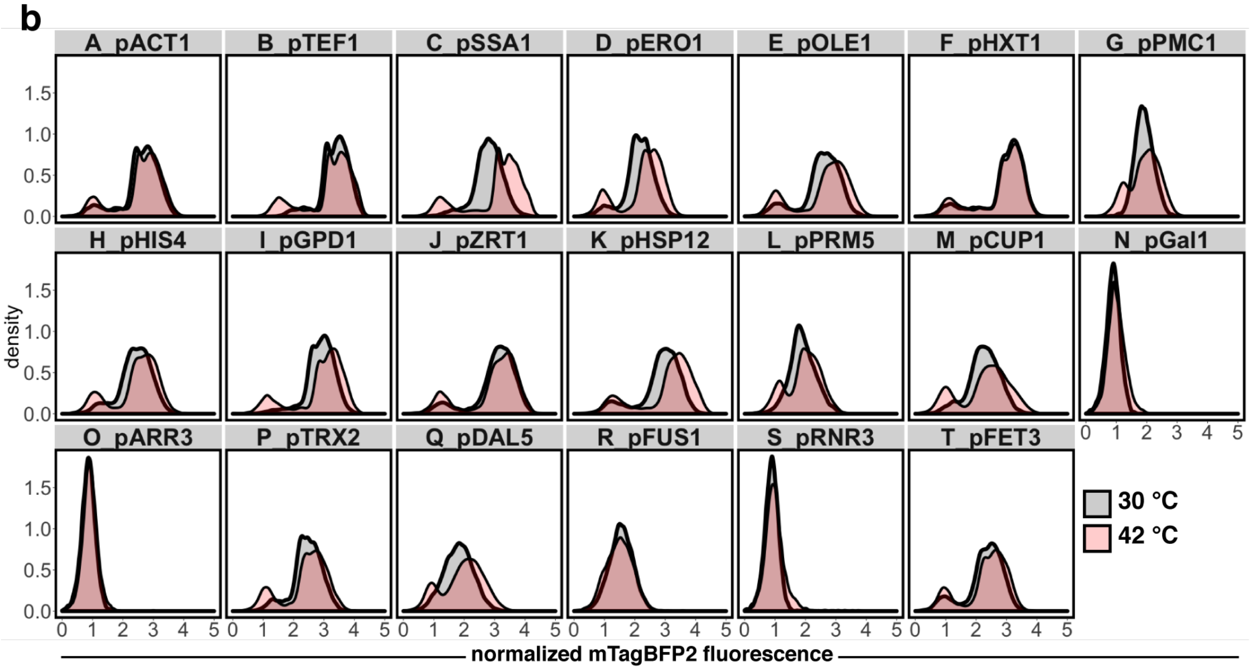


**Supplementary Figure 16a-b**. Multiplexed expression analysis histograms. FRAME-tags were used to profile responses of mixtures of *promoter*-mTagBFP2 reporter strains co-exposed to standard conditions (grey) or **(a)** media with 50 mM dithiothreitol (DTT, red) or **(b)** heat shock at 42 °C (red) for 6 hours. The mTagBFP2 fluorescent signal was normalized by side scatter and plotted on a logicle scale.

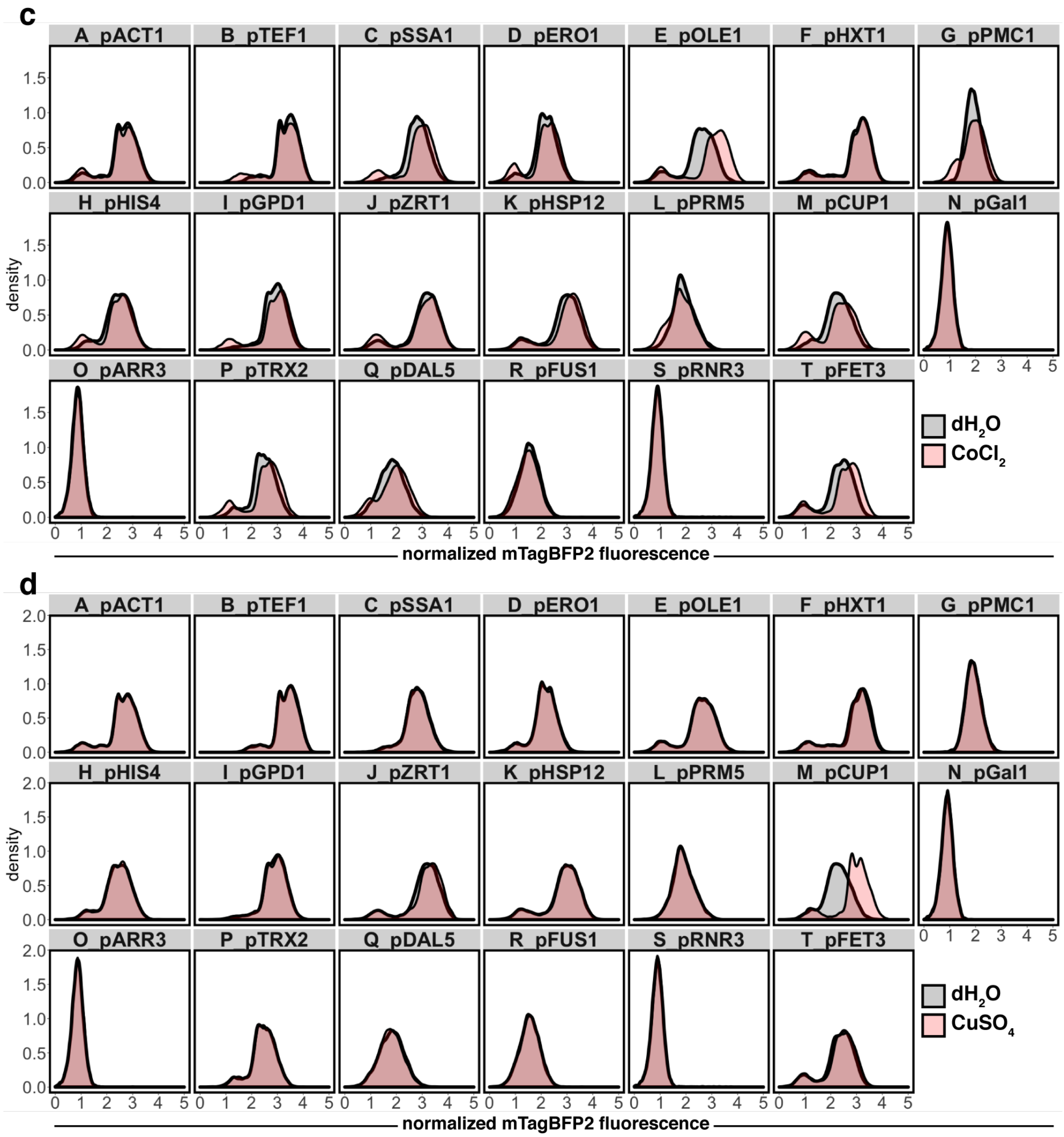


**Supplementary Figure 16c-d**. Multiplexed expression analysis histograms. FRAME-tags were used to profile responses of mixtures of *promoter*-mTagBFP2 reporter strains co-exposed to standard conditions (grey) or **(c)** media with 400 µM cobalt chloride ($CoCl_2$, red) or **(d)** 500 µM copper sulfate ($CuSO_4$, red) for 6 hours. The mTagBFP2 fluorescent signal was normalized by side scatter and plotted on a logicle scale.

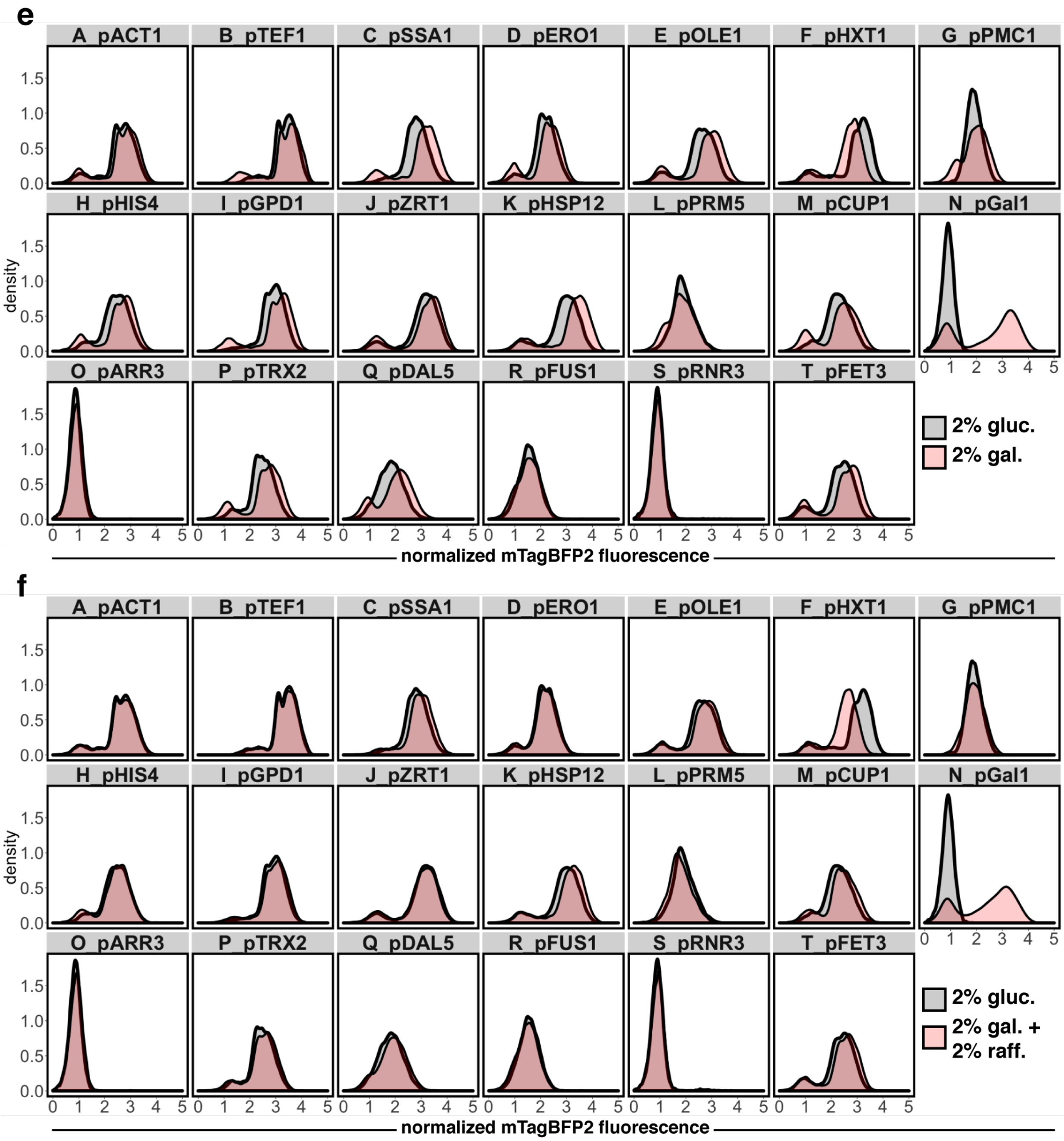


**Supplementary Figure 16e-f**. Multiplexed expression analysis histograms. FRAME-tags were used to profile responses of mixtures of *promoter*-mTagBFP2 reporter strains co-cultured in media with standard carbon source (2% glucose, grey) or **(e)** an alternate carbon source (2% galactose, red) or **(f)** a mixed carbon source (2% galactose / 2% raffinose, red) for 6 hours. The mTagBFP2 fluorescent signal was normalized by side scatter and plotted on a logicle scale.

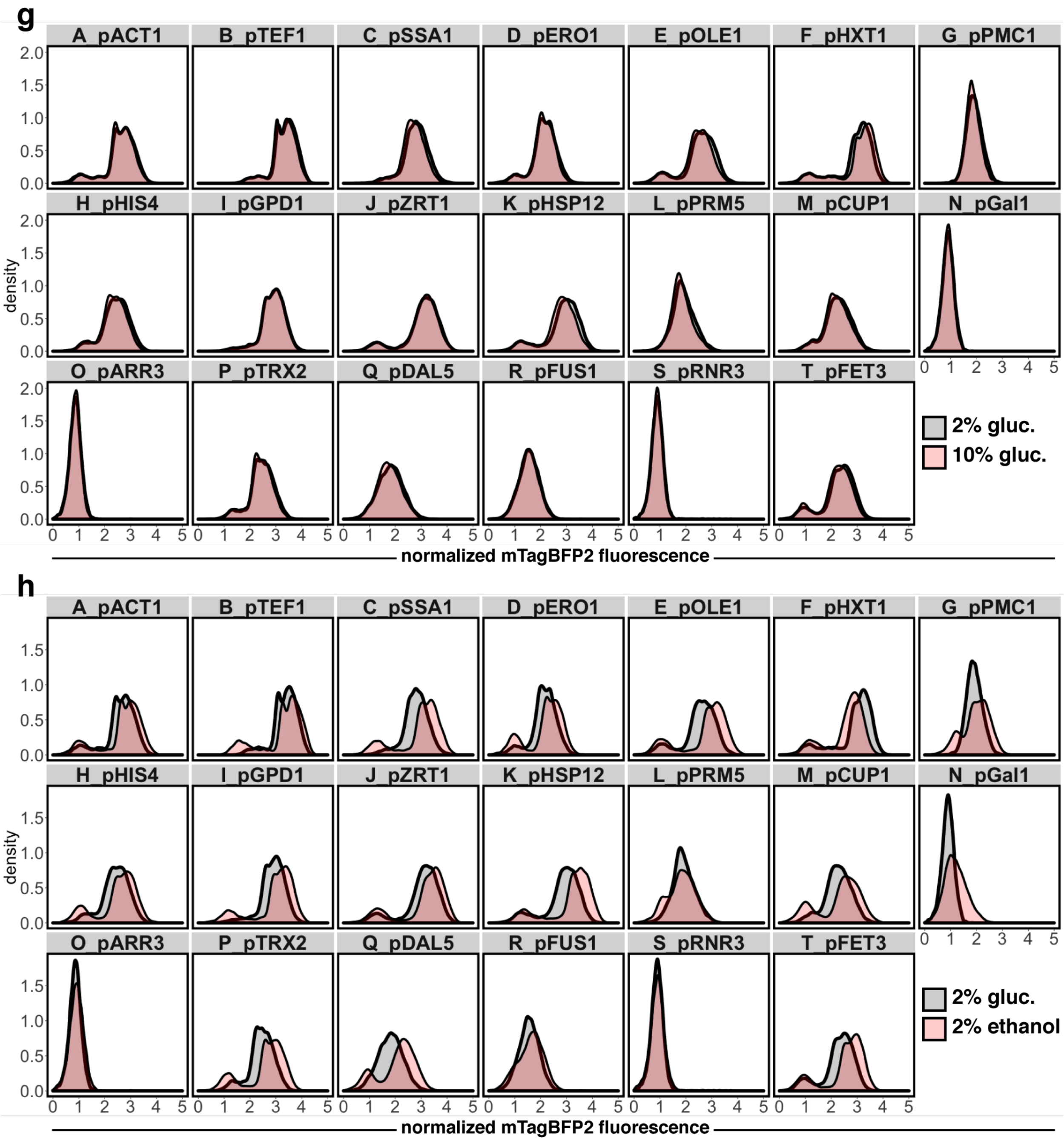


**Supplementary Figure 16g-h**. Multiplexed expression analysis histograms. FRAME-tags were used to profile responses of mixtures of *promoter*-mTagBFP2 reporter strains co-cultured in media with standard carbon source (2% glucose, grey) or **(g)** an increased amount of the carbon source (10% glucose, red) or **(h)** a non-standard carbon source (2% ethanol, red) for 6 hours. The mTagBFP2 fluorescent signal was normalized by side scatter and plotted on a logicle scale.

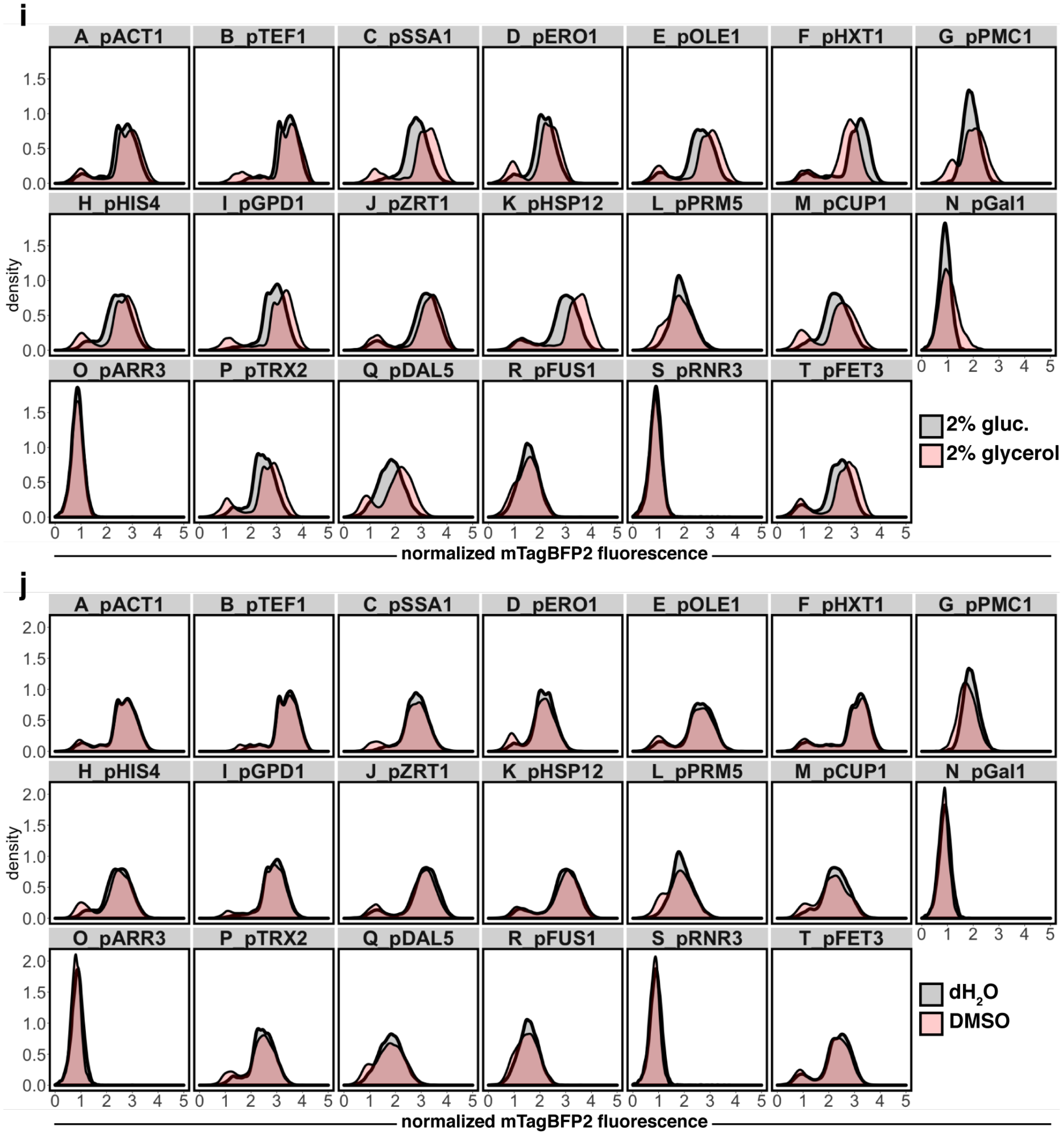


**Supplementary Figure 16i-j**. Multiplexed expression analysis histograms. FRAME-tags were used to profile responses of mixtures of *promoter*-mTagBFP2 reporter strains co-cultured in media with standard carbon source (2% glucose, grey) or **(i)** a non-standard carbon source (2% glucose, red) or **(j)** standard media containing 5% dimethylslfoxide (DMSO, red) for 6 hours. The mTagBFP2 fluorescent signal was normalized by side scatter and plotted on a logicle scale.

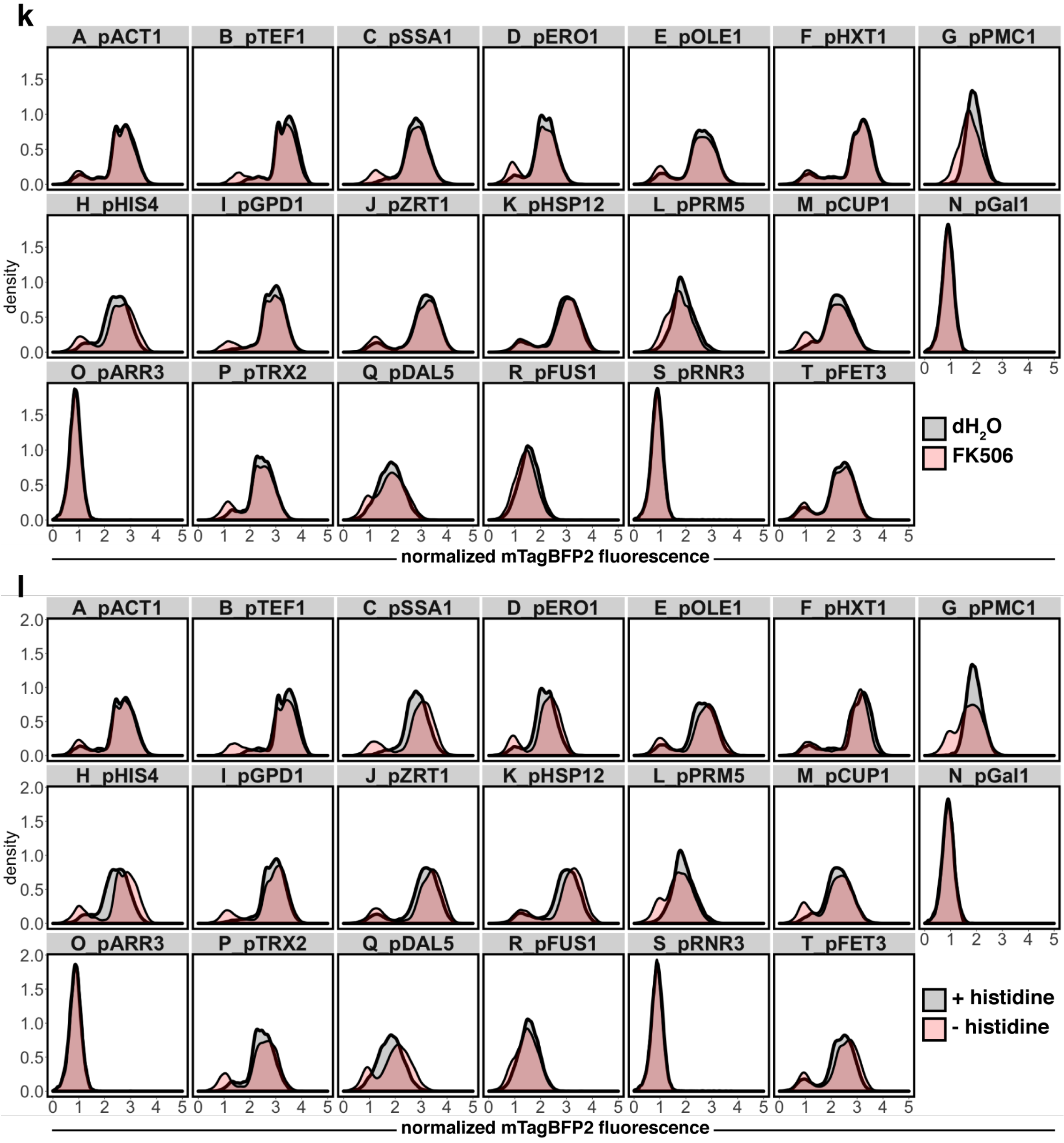


**Supplementary Figure 16k-l**. Multiplexed expression analysis histograms. FRAME-tags were used to profile responses of mixtures of *promoter*-mTagBFP2 reporter strains co-exposed to standard conditions (grey) or **(k)** media with 5 μM FK506 (red) or **(l)** media lacking an essential amino acid (- histidine, red) for 6 hours. The mTagBFP2 fluorescent signal was normalized by side scatter and plotted on a logicle scale.

m

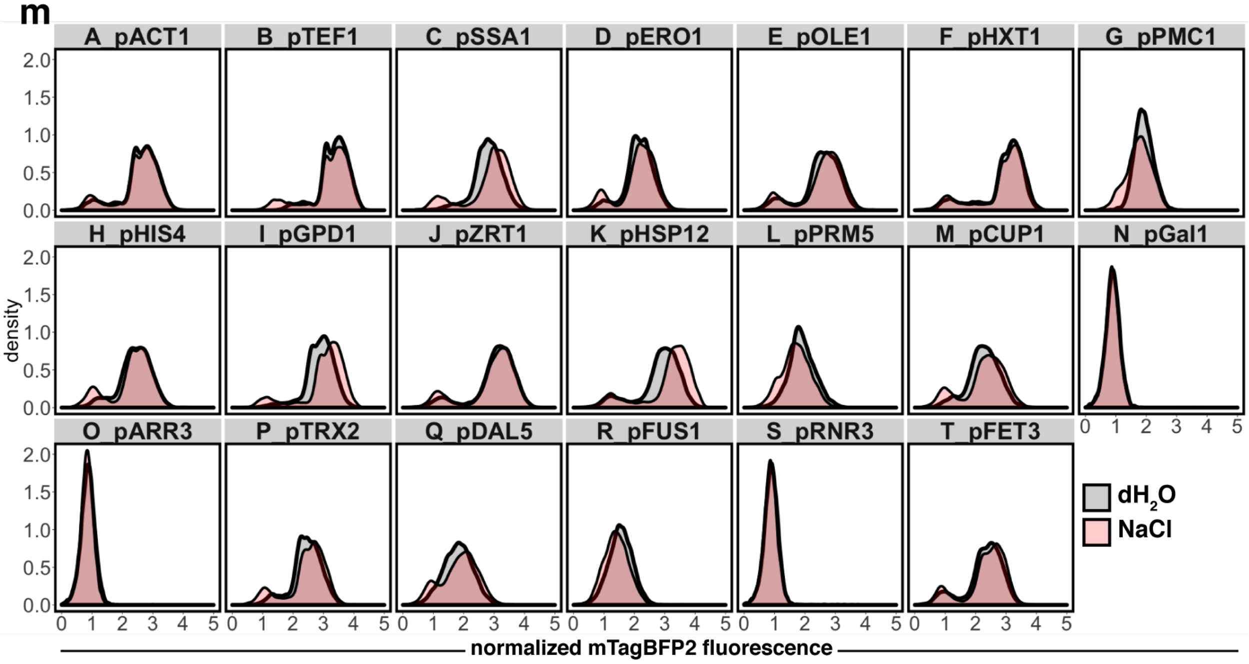


n

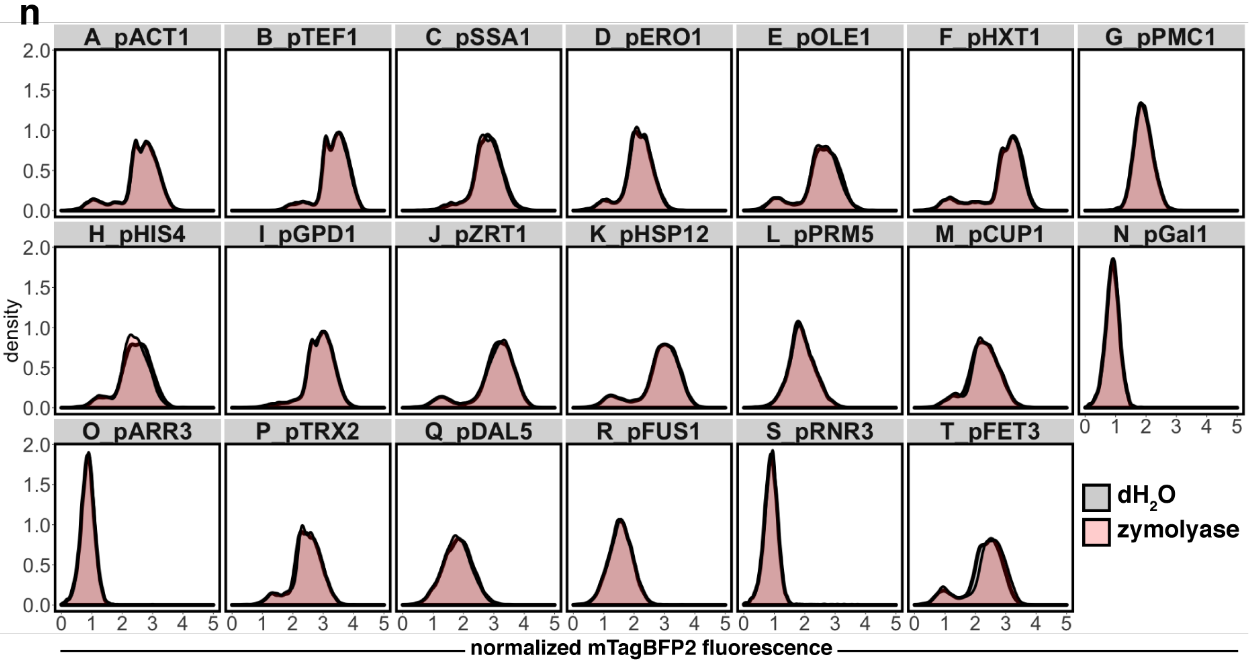


**Supplementary Figure 16m-n**. Multiplexed expression analysis histograms. FRAME-tags were used to profile responses of mixtures of *promoter*-mTagBFP2 reporter strains co-exposed to standard conditions (grey) or **(m)** osmotic shock with in media containing 0.7 M sodium chloride (NaCl, red) or **(n)** media with 5 units zymolyase a cell wall-degrading enzyme (red) for 6 hours. The mTagBFP2 fluorescent signal was normalized by side scatter and plotted on a logicle scale.

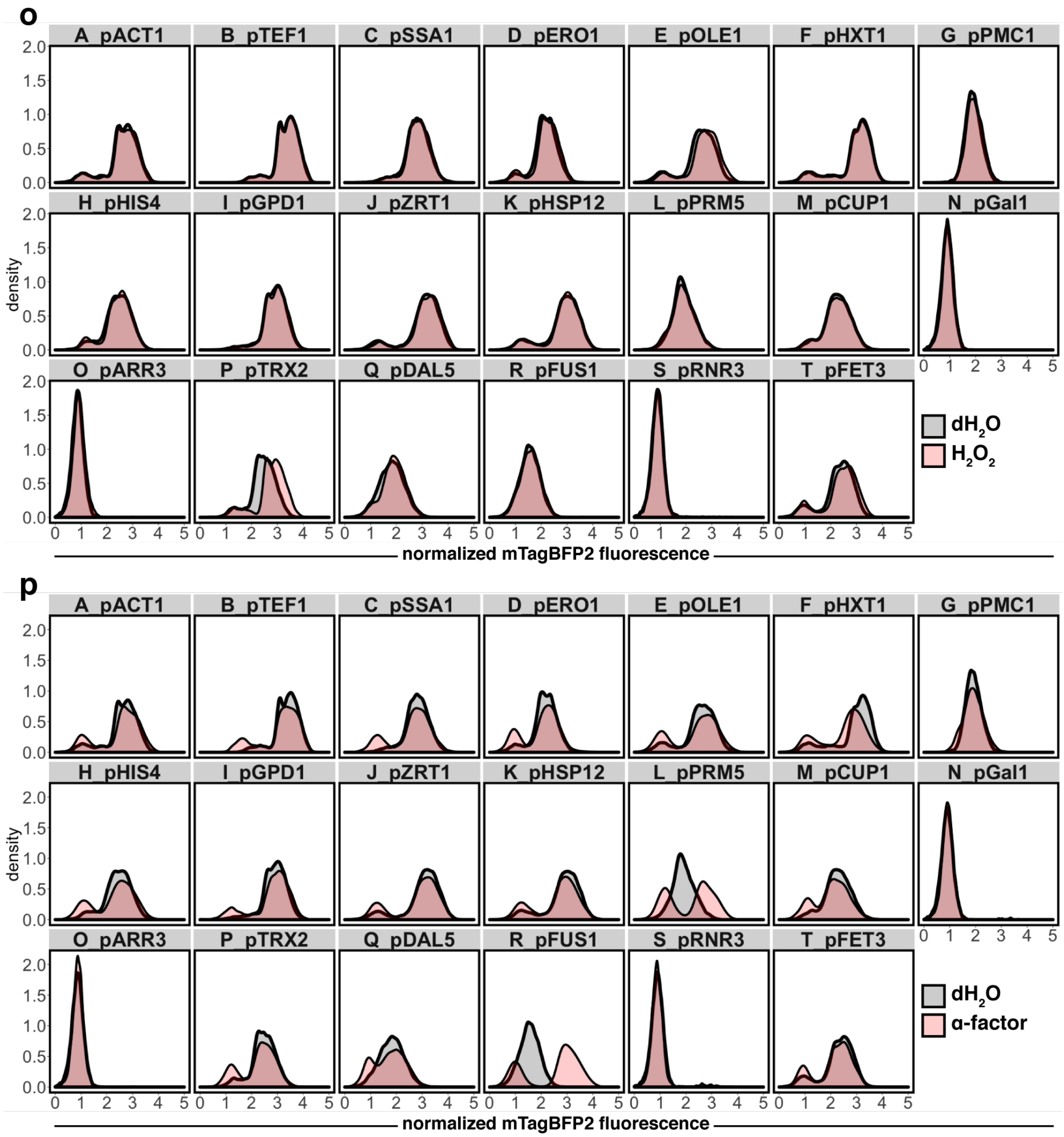


**Supplementary Figure 16o-p**. Multiplexed expression analysis histograms. FRAME-tags were used to profile responses of mixtures of *promoter*-mTagBFP2 reporter strains co-exposed to standard conditions (grey) or **(o)** oxidative shock with in media containing 1 mM hydrogen peroxide ($H_2O_2$, red) or **(p)** media with 5 μM α-factor a yeast peptide pheromone (red) for 6 hours. The mTagBFP2 fluorescent signal was normalized by side scatter and plotted on a logicle scale.

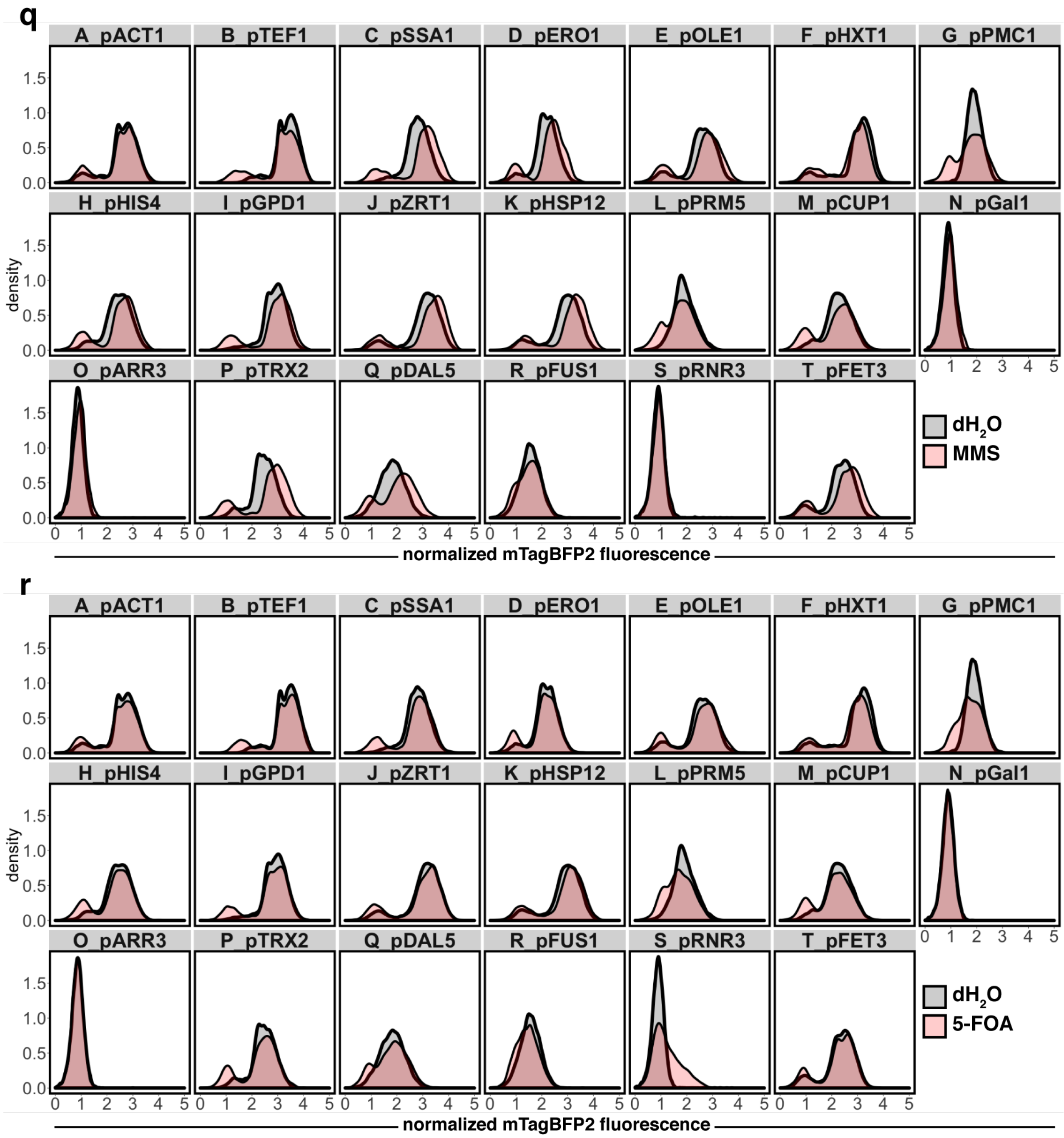


**Supplementary Figure 16q-r**. Multiplexed expression analysis histograms. FRAME-tags were used to profile responses of mixtures of *promoter*-mTagBFP2 reporter strains co-exposed to standard conditions (grey) or **(q)** media containing 0.1% methyl methanesulfonate a genotoxin (MMS, red) or **(r)** media with 0.01% 5-fluoroorotic acid which is converted to the antimetabolite 5-fluorouracil (5-FOA, red) for 6 hours. The mTagBFP2 fluorescent signal was normalized by side scatter and plotted on a logicle scale.

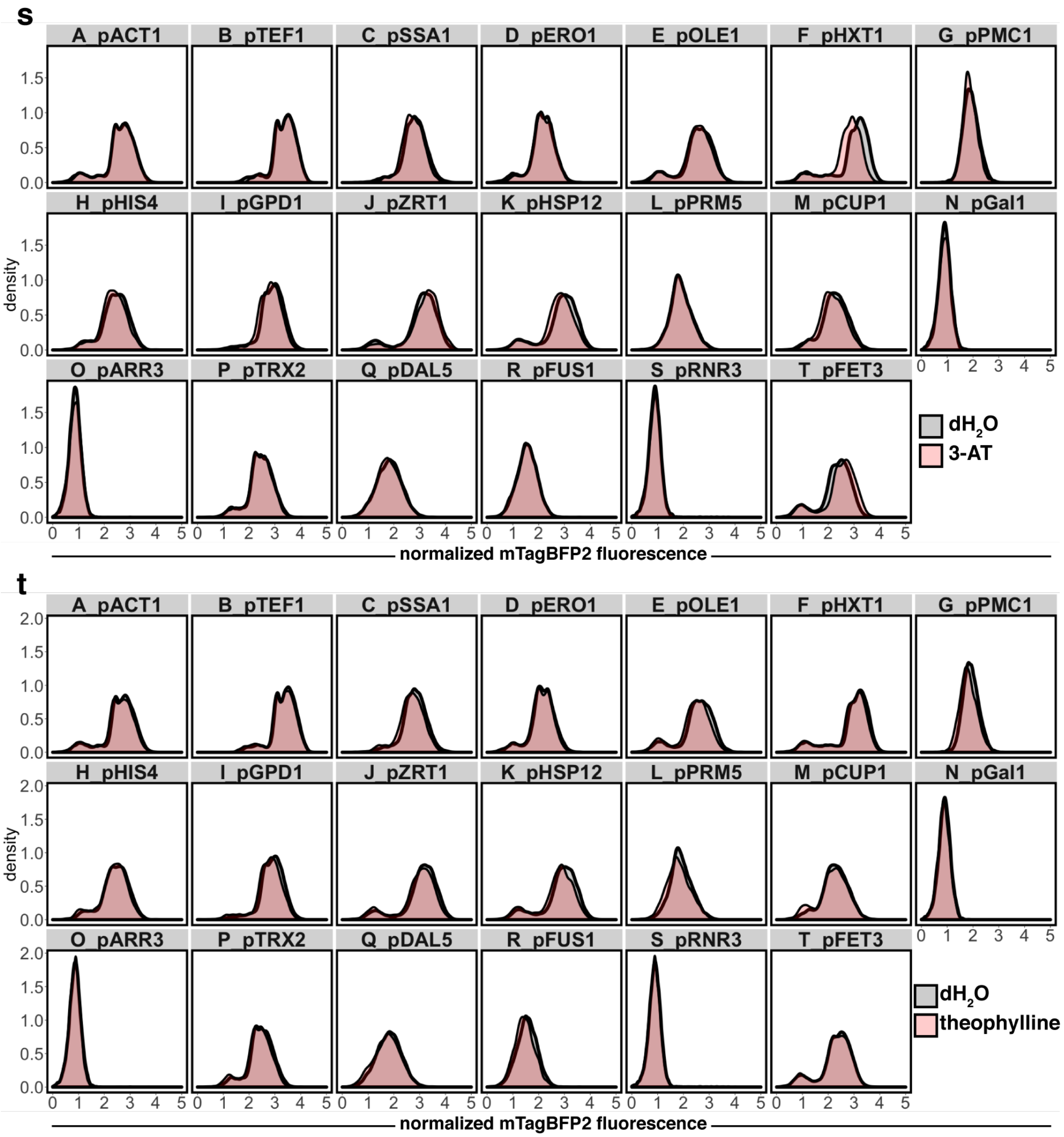


**Supplementary Figure 16s-t**. Multiplexed expression analysis histograms. FRAME-tags were used to profile responses of mixtures of *promoter*-mTagBFP2 reporter strains co-exposed to standard conditions (grey) or **(s)** media containing 50 mM 3-amino-1,2,4-triazole a competitive inhibitor of histidine metabolism (3-AT, red) or **(t)** media with 40 mM theophylline (red) for 6 hours. The mTagBFP2 fluorescent signal was normalized by side scatter and plotted on a logicle scale.

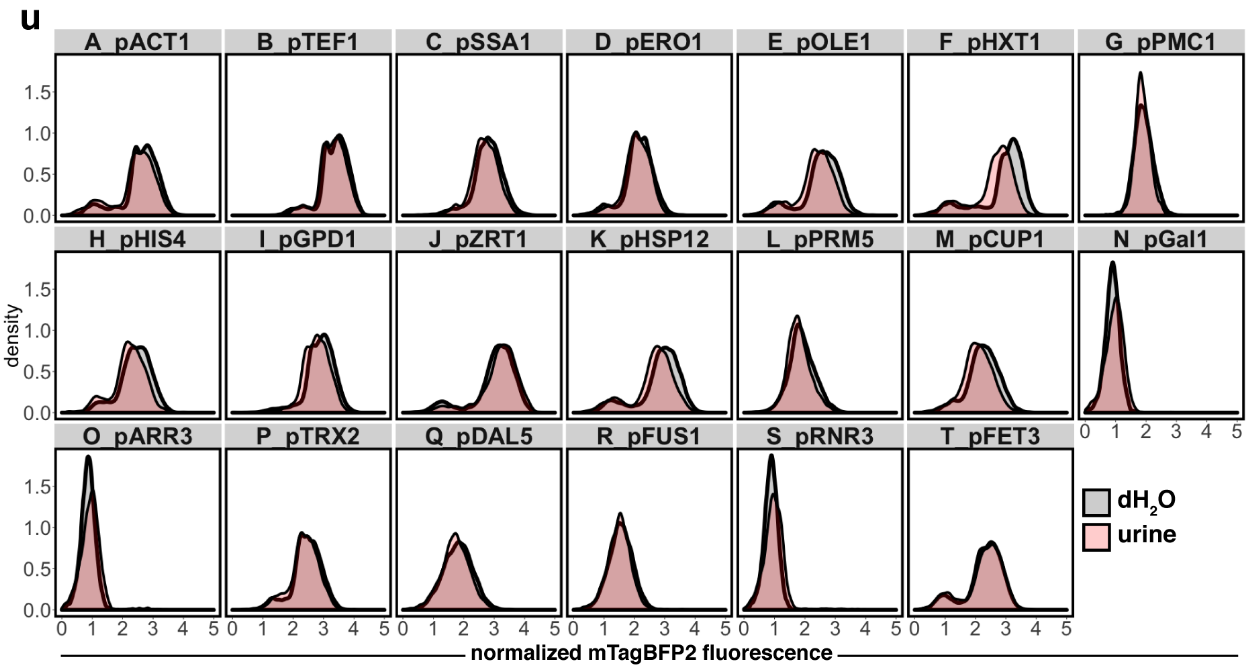


**Supplementary Figure 16u**. Multiplexed expression analysis histograms. FRAME-tags were used to profile responses of mixtures of *promoter*-mTagBFP2 reporter strains co-exposed to standard conditions (grey) or **(s)** media containing 50% human urine (red) for 6 hours. The mTagBFP2 fluorescent signal was normalized by side scatter and plotted on a logicle scale.

**Supplementary Table 1**. Sequences of frameshift motifs

| **Motif** | **Sequence** |
|---|---|
| fs-0.3 (**I**) | AA-GACTTTAAACTAGTTGACGCG**CAA**CTA**ATTCAGGC**CGCGTTAAAC**GTT**CTAGAAGGCGGTTCTATGGGAATGTCTGGA |
| fs-1.8 (**II**) | AA-GACTTTAAACTAGTTGACGCG**GGT**CTA**GTCAACAA**CGCGTTAAAC**CCA**CTAGAAGGCGGTTCTATGGGAATGTCTGGA |
| fs-3.3 (**III**) | AA-GACTTTAAACTAGTTGACGCG**GCT**CTA**GTCGCAGT**CGCGTTAAAC**AAG**CTAGAAGGCGGTTCTATGGGAATGTCTGGA |
| fs-4.2 (**IV**) | AA-GACTTTAAACTAGTTGACGCG**TCG**CTA**ACACGTGG**CGCGTTAAAC**CAT**CTAGAAGGCGGTTCTATGGGAATGTCTGGA |
| fs-7.8 (**V**) | AA-GACTTTAAACTAGTTGACGCG**TCA**CTA**GTAGCGGG**CGCGTTAAAC**AAA**CTAGAAGGCGGTTCTATGGGAATGTCTGGA |
| fs-9.4 (**VI**) | AA-GACTTTAAACTAGTTGACGCG**GAT**CTA**GCTTGTAA**CGCGTTAAAC**CAG**CTAGAAGGCGGTTCTATGGGAATGTCTGGA |
| fs-20 (**VII**) | AA-GACTTTAAACTAGTTGACGCG**AGT**CTA**GGGGATAA**CGCGTTAAAC**TTC**CTAGAAGGCGGTTCTATGGGAATGTCTGGA |
| fs-30 (**VIII**) | AA-GACTTTAAACTAGTTGACGCG**TCG**CTA**CCGCCCGG**CGCGTTAAAC**ACA**CTAGAAGGCGGTTCTATGGGAATGTCTGGA |
| fs-100 (**IX**) | AA**A**GACTTTAAACTAGTTGACGCG**GGT**CTA**GTCAACAA**CGCGTTAAAC**CCA**CTAGAAGGCGGTTCTATGGGAATGTCTGGA |

**Supplementary Table 2**. FRAME-tag constructs

| FT# | FP nom. | Construct assembly |
|---|---|---|
| FT1 | G100R | pTDH3 • **yEGFP** • [*NheI*] • L2 • **fs-100** • [*AatII*] • **mCherry** • tCaADH1 |
| FT2 | G30R | pTDH3 • **yEGFP** • [*NheI*] • L2 • **fs-30** • [*AatII*] • **mCherry** • tCaADH1 |
| FT3 | G9.4R | pTDH3 • **yEGFP** • [*NheI*] • L2 • **fs-9.4** • [*AatII*] • **mCherry** • tCaADH1 |
| FT4 | G4.2R | pTDH3 • **yEGFP** • [*NheI*] • L2 • **fs-4.2** • [*AatII*] • **mCherry** • tCaADH1 |
| FT5 | G0.3R | pTDH3 • **yEGFP** • [*NheI*] • L2 • **fs-0.3** • [*AatII*] • **mCherry** • tCaADH1 |
| FT6 | R30G | pTDH3 • **mCherry** • [*NheI*] • L2 • **fs-30** • [*AatII*] • **yEGFP** • tCaADH1 |
| FT7 | 30G100R | pTDH3 • L1 • **fs-30** • [*SalI*] • **yEGFP** • [*NheI*] • L2 • **fs-100** • [*AatII*] • **mCherry** • tCaADH1 |
| FT8 | 30G30R | pTDH3 • L1 • **fs-30** • [*SalI*] • **yEGFP** • [*NheI*] • L2 • **fs-30** • [*AatII*] • **mCherry** • tCaADH1 |
| FT9 | 30G9.4R | pTDH3 • L1 • **fs-30** • [*SalI*] • **yEGFP** • [*NheI*] • L2 • **fs-9.4** • [*AatII*] • **mCherry** • tCaADH1 |
| FT10 | 30G0.3R | pTDH3 • L1 • **fs-30** • [*SalI*] • **yEGFP** • [*NheI*] • L2 • **fs-0.3** • [*AatII*] • **mCherry** • tCaADH1 |
| FT11 | R9.4G | pTDH3 • **mCherry** • [*NheI*] • L2 • **fs-9.4** • [*AatII*] • **yEGFP** • tCaADH1 |
| FT12 | 30R30G | pTDH3 • L1 • **fs-30** • [*SalI*] • **mCherry** • [*NheI*] • L2 • **fs-30** • [*AatII*] • **yEGFP** • tCaADH1 |
| FT13 | 9.4G100R | pTDH3 • L1 • **fs-9.4** • [*SalI*] • **yEGFP** • [*NheI*] • L2 • **fs-100** • [*AatII*] • **mCherry** • tCaADH1 |
| FT15 | 9.4G0.3R | pTDH3 • L1 • **fs-9.4** • [*SalI*] • **yEGFP** • [*NheI*] • L2 • **fs-0.3** • [*AatII*] • **mCherry** • tCaADH1 |
| FT16 | R4.2G | pTDH3 • **mCherry** • [*NheI*] • L2 • **fs-4.2** • [*AatII*] • **yEGFP** • tCaADH1 |
| FT17 | 30R9.4G | pTDH3 • L1 • **fs-30** • [*SalI*] • **mCherry** • [*NheI*] • L2 • **fs-9.4** • [*AatII*] • **yEGFP** • tCaADH1 |
| FT19 | 4.2G100R | pTDH3 • L1 • **fs-4.2** • [*SalI*] • **yEGFP** • [*NheI*] • L2 • **fs-100** • [*AatII*] • **mCherry** • tCaADH1 |
| FT20 | R0.3G | pTDH3 • **mCherry** • [*NheI*] • L2 • **fs-0.3** • [*AatII*] • **yEGFP** • tCaADH1 |
| FT21 | 30R0.3G | pTDH3 • L1 • **fs-30** • [*SalI*] • **mCherry** • [*NheI*] • L2 • **fs-0.3** • [*AatII*] • **yEGFP** • tCaADH1 |
| FT22 | 9.4R0.3G | pTDH3 • L1 • **fs-9.4** • [*SalI*] • **mCherry** • [*NheI*] • L2 • **fs-0.3** • [*AatII*] • **yEGFP** • tCaADH1 |
| FT23 | B100R | pTDH3 • **mTagBFP2** • [*NheI*] • L2 • **fs-100** • [*AatII*] • **mCherry** • tCaADH1 |
| FT24 | B0.3R | pTDH3 • **mTagBFP2** • [*NheI*] • L2 • **fs-0.3** • [*AatII*] • **mCherry** • tCaADH1 |
| FT25 | G100B | pTDH3 • **yEGFP** • [*NheI*] • L2 • **fs-100** • [*AatII*] • **mTagBFP2** • tCaADH1 |
| FT26 | G100R100B | pTDH3 • **yEGFP** • [*NheI*] • L2 • **fs-100** • [*AatII*] • **mCherry** • [*AfeI*] • L3 • **fs-100** • [*ClaI*] • **mTagBFP2** • tCaADH1 |
| FT27 | 30G100R100B | pTDH3 • L1 • **fs-30** • [*SalI*] • **yEGFP** • [*NheI*] • L2 • **fs-100** • [*AatII*] • **mCherry** • [*AfeI*] • L3 • **fs-100** • [*ClaI*] • **mTagBFP2** • tCaADH1 |

FTs are provided in FP/*fs* nomenclature where G=yEGFP, R=mCherry, B=mTagBFP2 and # = *fs* (e.g. G30R) and full construct sequence layout. See DNA sequences of each part in **Supplementary Table 1** and **Supplementary Table 6**. Restriction sites facilitate replacement of FP and *fs* modules.

**Supplementary Table 3**. Strains used in this study. Strains were generated in this study except where a source is noted. Genotypes of FRAME-tagged strains given in fluorescent protein (FP) nomenclature (e.g. G30R, see Supplementary Table 2)

| Strain | Genotype | Comments |
|---|---|---|
| *Parent strains* | | |
| FY251 | *MAT*a *his3-Δ200 leu2-Δ1 trp1-Δ63 ura3-52* | ATCC 96098 |
| MaV203 | *MATα leu2-3,112 trp1-901 his3Δ200 ade2-101 cyh2R can1R gal4Δ gal80Δ GAL1::lacZ* $HIS3_{UASGAL1}$*::HIS3@LYS2 SPAL10::URA3* | Invitrogen |
| *fs characterization and initial FRAME-tag (FT) strains* | | |
| yFT1 | FY251 *leu2Δ::CgLEU2–G100R* | Also a final dual-FP red/green FT strain |
| yFT2 | FY251 *leu2Δ::CgLEU2–G30R* | Also a final dual-FP red/green FT strain |
| yG20R | FY251 *leu2Δ: CgLEU2–G20R* | |
| yFT3 | FY251 *leu2Δ::CgLEU2–G9.4R* | Also a final dual-FP red/green FT strain |
| yG7.8R | FY251 *leu2Δ::CgLEU2–G7.8R* | |
| yFT4 | FY251 *leu2Δ::CgLEU2–G4.2R* | Also a final dual-FP red/green FT strain |
| yG3.3R | FY251 *leu2Δ::CgLEU2–G3.3R* | |
| yG1.8R | FY251 *leu2Δ::CgLEU2–G1.8R* | |
| yFT5 | FY251 *leu2Δ::CgLEU2–G0.3R* | Also a final dual-FP red/green FT strain |
| *Plasmid characterization strains* | | |
| yPFT1 | FY251 + pFT1 | Used in Supp. Fig. 3 |
| yPFT2 | FY251 + pFT1 | Used in Supp. Fig. 3 |
| yPFT6 | FY251 + pFT6 | Used in Supp. Fig. 3 |
| yPG7.8R | FY251 + pG7.8R | Used in Supp. Fig. 3 |
| yPG1.8R | FY251 + pG1.8R | Used in Supp. Fig. 3 |
| *dual-FP red/green FT strains* | | |
| yFT6 | FY251 *leu2Δ::CgLEU2–R30G* | |
| yFT7 | FY251 *leu2Δ::CgLEU2–30G100R* | |
| yFT8 | FY251 *leu2Δ::CgLEU2–30G30R* | |
| yFT9 | FY251 *leu2Δ::CgLEU2–30G9.4R* | |
| yFT10 | FY251 *leu2Δ::CgLEU2–30G0.3R* | |
| yFT11 | FY251 *leu2Δ::CgLEU2–R9.4G* | |
| yFT12 | FY251 *leu2Δ::CgLEU2–30R30G* | |
| yFT13 | FY251 *leu2Δ::CgLEU2–9.4G100R* | |
| yFT15 | FY251 *leu2Δ::CgLEU2–9.4G0.3R* | |

| | | |
|---|---|---|
| yFT16 | FY251 *leu2Δ::CgLEU2–R4.2G* | |
| yFT17 | FY251 *leu2Δ::CgLEU2–30R9.4G* | |
| yFT19 | FY251 *leu2Δ::CgLEU2–4.2G100R* | |
| yFT20 | FY251 *leu2Δ::CgLEU2–R0.3G* | |
| yFT21 | FY251 *leu2Δ::CgLEU2–30R0.3G* | |
| yFT22 | FY251 *leu2Δ::CgLEU2–9.4R0.3G* | |
| *dual-FP red/green/blue FT strains* | | |
| yFT23 | FY251 *leu2Δ::CgLEU2–B100R* | |
| yFT24 | FY251 *leu2Δ::CgLEU2–B0.3R* | |
| yFT25 | FY251 *leu2Δ::CgLEU2–G100B* | |
| *triple-FP red/green/blue FT strains* | | |
| yFT26 | FY251 *leu2Δ::CgLEU2–G100R100B* | |
| yFT27 | FY251 *leu2Δ::CgLEU2–30G100R100B* | |
| *Strains used to analyze compatibility third FPs* | | |
| yFT0pBFP | FY251 + pGal-BFP | Used in Supp. Fig. 6 |
| yFT0pTurq | FY251 + pGal-Turq | Used in Supp. Fig. 6 |
| yFT0pVenus | FY251 + pGal-Venus | Used in Supp. Fig. 6 |
| yFT0pKO2 | FY251 + pGal-KO2 | Used in Supp. Fig. 6 |
| yFT9pBFP | yFT9 + pGal-BFP | Used in Supp. Fig. 6 |
| yFT9pTurq | yFT9 + pGal-Turq | Used in Supp. Fig. 6 |
| yFT9pVenus | yFT9 + pGal-Venus | Used in Supp. Fig. 6 |
| yFT9pKO2 | yFT9 + pGal-KO2 | Used in Supp. Fig. 6 |
| *Strains used for multiplex transcriptional profiling* | | |
| yFT1p2 | yFT1 + pR2 | reporter: pTEF1 |
| yFT2p4 | yFT2 + pR4 | reporter: pSSA1 |
| yFT3p18 | yFT3 + pR18 | reporter: pERO1 |
| yFT4p16 | yFT4 + pR16 | reporter: pOLE1 |
| yFT5p19 | yFT5 + pR19 | reporter: pHXT1 |
| yFT6p15 | yFT6 + pR15 | reporter: pPMC1 |
| yFT7p13 | yFT7 + pR13 | reporter: pHIS4 |
| yFT8p5 | yFT8 + pR5 | reporter: pGDP1 |
| yFT9p10 | yFT9 + pR10 | reporter: pZRT1 |
| yFT10p3 | yFT10 + pR3 | reporter: pHSP12 |
| yFT11p20 | yFT11 + pR20 | reporter: pPRM5 |
| yFT12p9 | yFT12 + pR9 | reporter: pCUP1 |
| yFT13p22 | yFT13 + pGal-BFP | reporter: pGAL1 |
| yFT15p12 | yFT15 + pR12 | reporter: pARR3 |

| | | |
|---|---|---|
| yFT16p7 | yFT16 + pR7 | reporter: pTRX2 |
| yFT17p14 | yFT17 + pR14 | reporter: pDAL5 |
| yFT19p1 | yFT19 + pR1 | reporter: pACT1 |
| yFT20p6 | yFT20 + pR6 | reporter: pFUS1 |
| yFT21p8 | yFT21 + pR8 | reporter: pRNR3 |
| yFT22p11 | yFT22 + pR11 | reporter: pFET3 |
| *Strains used for community tracking* | | |
| A | MaV203 *leu2Δ::CgLEU2–R0.3G + pSynGal4-A* | tag: FT20 |
| B | MaV203 *leu2Δ::CgLEU2–G9.4R + pSynGal4-B* | tag: FT3 |
| C | MaV203 *leu2Δ::CgLEU2–30G30R + pSynGal4-C* | tag: FT8 |
| D | MaV203 *leu2Δ::CgLEU2–30G0.3R + pSynGal4-D* | tag: FT10 |
| E | MaV203 *leu2Δ::CgLEU2–9.4R0.3G + pSynGal4-E* | tag: FT22 |
| F | MaV203 *leu2Δ::CgLEU2–30G100R + pSynGal4-F* | tag: FT7 |
| G | MaV203 *leu2Δ::CgLEU2–G100R + pSynGal4-G* | tag: FT1 |
| H | MaV203 *leu2Δ::CgLEU2–R4.2G + pSynGal4-H* | tag: FT16 |
| I | MaV203 *leu2Δ::CgLEU2–30G9.4R + pSynGal4-I* | tag: FT9 |

**Supplementary Table 4**. Plasmids used in this study.

| Plasmid | Construct details | Comments |
|---|---|---|
| pRS416 | URA3, CEN6/ARS4, AmpR, ColE1 | ATCC 87521 |
| pRS424 | TRP1, 2μ, AmpR, ColE1 | ATCC 77105 |
| pGal-BFP | pRS416, pGAL1-mTagBFP2 | |
| pGal-Turq | pRS416, pGAL1-mTurquoise2 | |
| pGal-Venus | pRS416, pGAL1-mVenus | |
| pGal-KO2 | pRS416, pGAL1-mKO2 | |
| pG20R | pRS416, CgLEU2-G20R | framed with homology for LEU2 locus |
| pG7.8R | pRS416, CgLEU2-G7.8R | framed with homology for LEU2 locus |
| pG3.3R | pRS416, CgLEU2-G3.3R | framed with homology for LEU2 locus |
| pG1.8R | pRS416, CgLEU2-G1.8R | framed with homology for LEU2 locus |
| pFT1 | pRS416, CgLEU2–G100R | framed with homology for LEU2 locus |
| pFT2 | pRS416, CgLEU2–G30R | framed with homology for LEU2 locus |
| pFT3 | pRS416, CgLEU2–G9.4R | framed with homology for LEU2 locus |
| pFT4 | pRS416, CgLEU2–G4.2R | framed with homology for LEU2 locus |
| pFT5 | pRS416, CgLEU2–G0.3R | framed with homology for LEU2 locus |
| pFT6 | pRS416, CgLEU2–R30G | framed with homology for LEU2 locus |
| pFT7 | pRS416, CgLEU2–30G100R | framed with homology for LEU2 locus |
| pFT8 | pRS416, CgLEU2–30G30R | framed with homology for LEU2 locus |
| pFT9 | pRS416, CgLEU2–30G9.4R | framed with homology for LEU2 locus |
| pFT10 | pRS416, CgLEU2–30G0.3R | framed with homology for LEU2 locus |
| pFT11 | pRS416, CgLEU2–R9.4G | framed with homology for LEU2 locus |
| pFT12 | pRS416, CgLEU2–30R30G | framed with homology for LEU2 locus |
| pFT13 | pRS416, CgLEU2–9.4G100R | framed with homology for LEU2 locus |
| pFT15 | pRS416, CgLEU2–9.4G0.3R | framed with homology for LEU2 locus |
| pFT16 | pRS416, CgLEU2–R4.2G | framed with homology for LEU2 locus |
| pFT17 | pRS416, CgLEU2–30R9.4G | framed with homology for LEU2 locus |
| pFT19 | pRS416, CgLEU2–4.2G100R | framed with homology for LEU2 locus |
| pFT20 | pRS416, CgLEU2–R0.3G | framed with homology for LEU2 locus |
| pFT21 | pRS416, CgLEU2–30R0.3G | framed with homology for LEU2 locus |
| pFT22 | pRS416, CgLEU2–9.4R0.3G | framed with homology for LEU2 locus |
| pFT23 | pRS416, CgLEU2–B100R | framed with homology for LEU2 locus |
| pFT24 | pRS416, CgLEU2–B0.3R | framed with homology for LEU2 locus |
| pFT25 | pRS416, CgLEU2–G100B | framed with homology for LEU2 locus |
| pFT26 | pRS416, CgLEU2–G100R100B | framed with homology for LEU2 locus |
| pFT27 | pRS416, CgLEU2–30G100R100B | framed with homology for LEU2 locus |
| pR1 | pRS416, pACT1-mTagBFP2 | |

| | |
|---|---|
| pR2 | pRS416, pTEF1-mTagBFP2 |
| pR3 | pRS416, pHSP12-mTagBFP2 |
| pR4 | pRS416, pSSA1-mTagBFP2 |
| pR5 | pRS416, pGPD1-mTagBFP2 |
| pR6 | pRS416, pFUS1-mTagBFP2 |
| pR7 | pRS416, pTRX2-mTagBFP2 |
| pR8 | pRS416, pRNR3-mTagBFP2 |
| pR9 | pRS416, pCUP1-mTagBFP2 |
| pR10 | pRS416, pZRT1-mTagBFP2 |
| pR11 | pRS416, pFET3-mTagBFP2 |
| pR12 | pRS416, pARR3-mTagBFP2 |
| pR13 | pRS416, pHIS4-mTagBFP2 |
| pR14 | pRS416, pDAL5-mTagBFP2 |
| pR15 | pRS416, pPMC1-mTagBFP2 |
| pR16 | pRS416, pOLE1-mTagBFP2 |
| pR18 | pRS416, pERO1-mTagBFP2 |
| pR19 | pRS416, pHXT1-mTagBFP2 |
| pR20 | pRS416, pPRM5-mTagBFP2 |
| pSynGal4-A | pRS424, pADH-Gal4(BD)-fsA-Gal4(AD) |
| pSynGal4-B | pRS424, pADH-Gal4(BD)-fsB-Gal4(AD) |
| pSynGal4-C | pRS424, pADH-Gal4(BD)-fsC-Gal4(AD) |
| pSynGal4-D | pRS424, pADH-Gal4(BD)-fsD-Gal4(AD) |
| pSynGal4-E | pRS424, pADH-Gal4(BD)-fsE-Gal4(AD) |
| pSynGal4-F | pRS424, pADH-Gal4(BD)-fsF-Gal4(AD) |
| pSynGal4-G | pRS424, pADH-Gal4(BD)-fsG-Gal4(AD) |
| pSynGal4-H | pRS424, pADH-Gal4(BD)-fsH-Gal4(AD) |
| pSynGal4-I | pRS424, pADH-Gal4(BD)-fsI-Gal4(AD) |

pFT plasmids contain the integrating constructs of the FRAME-tags and were not directly used in experiments except were noted. The integrating construct can be excised with *SwaI*, which cuts these pFT plasmids three times (twice flanking the FT construct and once in the backbone to simplify purification).

**Supplementary Table 5**. Primers for promoter cloning.

| Primer | Sequence |
|---|---|
| pACT_for (MJ615) | ctcactaaagggaacaaaagctggagctctagtCCTTAAAAACATATGCCTCACCCT |
| pACT_rev (MJ616) | gcatattttctttaatcaattcagacattttctagaCAGTAAATTTTCGATCTTGGGAAG |
| pTEF1_for (MJ617) | ctcactaaagggaacaaaagctggagctctagtATAGCTTCAAAATGTTTCTACTCCT |
| pTEF_rev (MJ618) | gcatattttctttaatcaattcagacattttctagaTTAGATTGCTATGCTTTCTTTC |
| pHSP12_for(MJ619) | ctcactaaagggaacaaaagctggagctctagtAGTGAAAATCTCCGGGAGCG |
| pHSP12_rev (MJ620) | gcatattttctttaatcaattcagacattttctagaTGAGTTGTTTGTTTGAGATTATCG |
| pSSA1_for (MJ621) | ctcactaaagggaacaaaagctggagctctagtGGCATTTTCGTTCTTGTGGA |
| pSSA1_rev (MJ622) | gcatattttctttaatcaattcagacattttctagaATTTTTGTTTCTTGTAATACTTGA |
| pGPD1_for (MJ625) | ctcactaaagggaacaaaagctggagctctagtCTGGGGTTTGAGCAAGTCTA |
| pGPD1_rev (MJ626) | gcatattttctttaatcaattcagacattttctagaTTATCAATATTTGTGTTTGTGGAG |
| pFUS1_for (MJ627) | ctcactaaagggaacaaaagctggagctctagtTGCCTCAATCCTTCTTTTGCTT |
| pFUS1_rev (MJ628) | gcatattttctttaatcaattcagacattttctagaCTTGATGGCTTATATCCTGCTCT |
| pTRX2_for (MJ629) | ctcactaaagggaacaaaagctggagctctagtACTTTTACGGGTGGCAACG |
| pTRX2_rev (MJ630) | gcatattttctttaatcaattcagacattttctagaTCGTAGACTCTCGTGTATGTGTGC |
| pRNR3_for (MJ631) | ctcactaaagggaacaaaagctggagctctagtGTAATAACAAGCAGGTGGGCG |
| pRNR3_rev (MJ632) | gcatattttctttaatcaattcagacattttctagaTTATTGCTGCTGCTATTCTTGCTT |
| pCUP1_for (MJ635) | ctcactaaagggaacaaaagctggagctctagtTCACCACCCTTTATTTCAGGC |
| pCUP_rev (MJ636) | gcatattttctttaatcaattcagacattttctagaTGTGATGATTGATTGATTGATTGT |
| pZRT1_for (MJ637) | ctcactaaagggaacaaaagctggagctctagtGGCAAGAGTATTTCAGACTTTCCT |
| pZRT1_rev (MJ638) | gcatattttctttaatcaattcagacattttctagaTTTGTGCTGTTGTTTTATTGTCT |
| pFET3_for (MJ666) | ctcactaaagggaacaaaagctggagctctagtGATAATGCCTTGGCTTGCCT |
| pFET3_rev (MJ640) | gcatattttctttaatcaattcagacattttctagaTACTCTTCCTTACACTGGGGTCC |
| pARR3_for (MJ641) | ctcactaaagggaacaaaagctggagctctagtCACGTGCAAAATCTTCTCTTCG |
| pARR3_rev (MJ642) | gcatattttctttaatcaattcagacattttctagaCCTGATGATTTGTTGGTTGGGT |
| pHIS4_for (MJ643) | ctcactaaagggaacaaaagctggagctctagtAAACCCATGCACAGTGACTC |
| pHIS4_rev (MJ644) | gcatattttctttaatcaattcagacattttctagaATTGTATTACTATTACACAGCGCA |
| pDAL5_for (MJ649) | ctcactaaagggaacaaaagctggagctctagtAGCGTTCTCATCAGTCACTTG |
| pDAL4_rev (MJ650) | gcatattttctttaatcaattcagacattttctagaATCCTTGTTTTGTTGTTTTCTTCA |
| pPMC1_for (MJ651) | ctcactaaagggaacaaaagctggagctctagtGTTTTTACCCGGCAAAGAAGC |
| pPMC1_rev (MJ652) | gcatattttctttaatcaattcagacattttctagaTATTTTTTTTGTTACGCACACAGT |
| pOLE1_for (MJ653) | ctcactaaagggaacaaaagctggagctctagtCATGTCCCGGGGTTAGCG |
| pOLE1_rev (MJ654) | gcatattttctttaatcaattcagacattttctagaTTGTTGTAATGTTTTAGTGCTGT |
| pERO1_for (MJ623) | ctcactaaagggaacaaaagctggagctctagtAAAGAACACGGCGGTAAGAA |
| pERO1_rev (MJ624) | gcatattttctttaatcaattcagacattttctagaTTTACCTGCACGTTACTGTGG |
| pHXT1_for (MJ633) | ctcactaaagggaacaaaagctggagctctagtTGCAAAAAGCTTCCGATCCT |
| pHXT1_rev (MJ634) | gcatattttctttaatcaattcagacattttctagaCGTATATCAACTAGTTGACGATTA |
| pPRM5_for (MJ645) | ctcactaaagggaacaaaagctggagctctagtCTCACCCGGATCGTAGTCAC |
| pPRM5_rev (MJ646) | gcatattttctttaatcaattcagacattttctagaTCTTGCGTTTTGAGTGTCAATTT |

**Supplementary Table 6**. DNA sequences of FRAME-tag parts.

| Part | Sequence |
|---|---|
| pTDH3 | CAGTTCGAGTTTATCATTATCAATACTGCCATTTCAAAGAATACGTAAATAATTAATAGTAGTGATTTTCCTAACTT<br>TATTTAGTCAAAAAATTAGCCTTTTAATTCTGCTGTAACCCGTACATGCCCAAAATAGGGGGCGGGTTACACAGAA<br>TATATAACATCGTAGGTGTCTGGGTGAACAGTTTATTCCTGGCATCCACTAAATATAATGGAGCCCGCTTTTTAAG<br>CTGGCATCCAGAAAAAAAAAGAATCCCAGCACCAAAATATTGTTTTCTTCACCAACCATCAGTTCATAGGTCCATT<br>CTCTTAGCGCAACTACAGAGAACAGGGGCACAAACAGGCAAAAAACGGGCACAACCTCAATGGAGTGATGCAAC<br>CTGCCTGGAGTAAATGATGACACAAGGCAATTGACCCACGCATGTATCTATCTCATTTTCTTACACCTTCTATTACC<br>TTCTGCTCTCTCTGATTTGGAAAAAGCTGAAAAAAAAGGTTGAAACCAGTTCCCTGAAATTATTCCCCTACTTGAC<br>TAATAAGTATATAAAGACGGTAGGTATTGATTGTAATTCTGTAAATCTATTTCTTAAACTTCTTAAATTCTACTTTT<br>ATAGTTAGTCTTTTTTTTAGTTTTAAAACACCAAGAACTTAGTTTCGAATAAACACACATAAACAAACAAA |
| L1 | ATGGGTTCAGGTGAACAATCA |
| L2 | GGCAGCGGCGACTACAAGGACGACGACGAC |
| L3 | GGTGCATCTGGATCAGGACAATCA |
| *NheI* | GCTAGC |
| *AatII* | GACGTC |
| *SalI* | GTCGAC |
| *AfeI* | AGCGCT |
| *ClaI* | ATCGAT |
| yEGFP | ATGTCTAAAGGTGAAGAATTATTCACTGGTGTTGTCCCAATTTTGGTTGAATTAGATGGTGATGTTAATGGTCACA<br>AATTTTCTGTCTCCGGTGAAGGTGAAGGTGATGCTACTTACGGTAAATTGACCTTAAAATTTATTTGTACTACTGGT<br>AAATTGCCAGTTCCATGGCCAACCTTAGTCACTACTTTCGGTTATGGTGTTCAATGTTTTGCTAGATACCCAGATCA<br>TATGAAACAACATGACTTTTTCAAGTCTGCCATGCCAGAAGGTTATGTTCAAGAAAGAACTATTTTTTTCAAAGAT<br>GACGGTAACTACAAGACCAGAGCTGAAGTCAAGTTTGAAGGTGATACCTTAGTTAATAGAATCGAATTAAAAGGT<br>ATTGATTTTAAAGAAGATGGTAACATTTTAGGTCACAAATTGGAATACAACTATAACTCTCACAATGTTTACATCA<br>TGGCTGACAAACAAAAGAATGGTATCAAAGTTAACTTCAAAATTAGACACAACATTGAAGATGGTTCTGTTCAAT<br>TAGCTGACCATTATCAACAAAATACTCCAATTGGTGATGGTCCAGTCTTGTTACCAGACAACCATTACTTATCCAC<br>TCAATCTGCCTTATCCAAAGATCCAAACGAAAAGAGAGACCACATGGTCTTGTTAGAATTTGTTACTGCTGCTGGT<br>ATTACCCATGGTATGGATGAATTGTACAAA |
| mCherry | ATGGTTTCAAAAGGTGAAGAAGATAATATGGCTATTATTAAAGAATTTATGAGATTTAAAGTTCATATGGAAGGTT<br>CAGTTAATGGTCATGAATTTGAAATTGAAGGTGAAGGTGAAGGTAGACCATATGAAGGTACTCAAACTGCTAAAT<br>TGAAAGTTACTAAAGGTGGTCCATTACCATTTGCTTGGGATATTTTGTCACCACAATTTATGTATGGTTCAAAAGCT<br>TATGTTAAACATCCAGCTGATATTCCAGATTATTTAAAATTGTCATTTCCAGAAGGTTTTAAATGGGAAAGAGTTA<br>TGAATTTTGAAGATGGTGGTGTTGTTACTGTTACTCAAGATTCATCATTACAAGATGGTGAATTTATTTATAAAGTT<br>AAATTGAGAGGTACTAATTTTCCATCAGATGGTCCAGTTATGCAAAAAAAAACTATGGGTTGGGAAGCTTCATCA<br>GAAAGAATGTATCCAGAAGATGGTGCTTTAAAAGGTGAAATTAAACAAAGATTGAAATTAAAAGATGGTGGTCAT<br>TATGATGCTGAAGTTAAAACTACTTATAAAGCTAAAAAACCAGTTCAATTACCAGGTGCTTATAATGTTAATATTA<br>AATTGGATATTACTTCACATAATGAAGATTATACTATTGTTGAACAATATGAAAGAGCTGAAGGTAGACATTCAAC<br>TGGTGGTATGGATGAATTATATAAAagcgctGGTGCATCTGGATCAGGACAATCAAAAGACTTTAAACTAGTTGACG<br>CGGGTCTAGTCAACAACGCGTTAAACCCACTAGAAGGCGGTTCTATGGGAATGTCTGGA |
| mTagBFP2 | ATGTCTGAATTGATTAAAGAAAATATGCATATGAAATTGTACATGGAAGGTACTGTTGATAATCATCATTTCAAAT<br>GTACTTCAGAAGGTGAAGGTAAGCCATACGAAGGTACTCAAACTATGAGAATTAAAGTTGTTGAAGGTGGTCCAT<br>TGCCATTCGCTTTTGATATTTTGGCTACTTCATTTTTATATGGTTCAAAGACTTTTATTAATCATACTCAAGGTATTC<br>CAGATTTTTTCAAACAATCATTCCCAGAAGGTTTCACTTGGGAAAGAGTTACTACTTACGAAGATGGTGGTGTTTT<br>AACTGCTACTCAAGATACTTCTTTGCAAGATGGTTGTTTGATTTACAATGTTAAAATTAGAGGTGTTAATTTCACTT<br>CAAATGGTCCAGTTATGCAAAAAAAGACTTTGGGTTGGGAAGCTTTTACTGAAACTTTGTACCCAGCTGATGGTGG<br>TTTGGAAGGTAGAAATGATATGGCTTTGAAGTTGGTTGGTGGTTCTCATTTGATTGCTAATGCTAAGACTACTTAT<br>AGATCAAAGAAGCCAGCTAAGAATTTGAAGATGCCAGGTGTTTATTACGTTGATTATAGATTGGAAAGAATTAAA<br>GAAGCTAATAATGAAACTTACGTTGAACAACATGAAGTTGCTGTTGCTAGATACTGTGATTTGCCATCAAAATTGG<br>GTCATAAATTGAAT |
| tCaADH1 | TAAGCAAATAGCTAAATTATATACGAATTAATATTATGATTAAGTGTTTACGTGAGTGCGATATTTTTATTACTATC<br>TTATACAGTTGTATATACTCTATAAAATGAGTTGTCTATTAATTAACGCGATGAATGCTTTCTGGGTTTACCTCTCC<br>AACAACTCTAGTTTACTTCTCAATACATTCAATTGTATTTGATTTGTCAATACTTCATCATTAATCAATTCTATAGTT<br>TTGTTTTTCTCGTTTATTTCCAAATTTAATGCATCAATTTTATTATTCAATTTGTCGTTGATTTTGGTTAATGATTTTA<br>TGGTTTGATCTCTGGCATTGATTGTTTGTGTTAGTTTTTCATTATTGATAattaaaTTATTTAAGTTAGTTATCAACTCG<br>GTGTTTTCAAGTTTCAAGTTTTCAATTTCTTTAGAGTTTATTAGATTTGTCAAAGTTTCTGAATTGCTTGATTGGTCC<br>TGTAGAAGAGTATTTGTTGTTGTGGATAATTGATTCAATTTTTGAGACAATTGCTGGAAGGCGTTGAAATATCTAG<br>CATCAATCTCATGGTTTTTTTCCCGAGAGTCTCGTAGATTCAATTGTTTTAATATATCTTGGGACCACTCTTGATTTG<br>AACTCATGGAAattaaaCTGGGTGTTGTGTTGTGGTGTAATGATTGTACCCCCTTTGCTTATAATTGTGTGG |

These parts and *fs* modules were assembled into FRAME-tags as shown in **Supplementary Table 2.**

**Supplementary Table 7**. DNA sequences of additional fluorescent proteins.

| Part | Sequence |
|---|---|
| mTurquoise2 | ATGTCTAAAGGTGAAGAATTATTCACTGGTGTTGTTCCAATTTTGGTTGAATTAGATGGTGATGTTAATGGTCAT<br>AAATTCTCTGTTTCTGGTGAAGGTGAAGGTGATGCTACTTATGGTAAGTTGACTTTGAAGTTTATTTGTACTACT<br>GGTAAATTGCCAGTTCCATGGCCAACTTTGGTTACTACTTTGTCTTGGGGTGTTCAATGTTTTGCTAGATATCCA<br>GATCATATGAAACAACATGATTTCTTTAAATCTGCTATGCCAGAAGGTTACGTTCAAGAAAGAACTATTTTCTTT<br>AAAGATGATGGTAATTACAAAACTAGAGCTGAAGTTAAATTCGAAGGTGATACTTTGGTTAATAGAATTGAATT<br>GAAGGGTATTGATTTCAAAGAAGATGGTAATATTTTGGGTCATAAGTTGGAATACAATTACTTCTCTGATAATGT<br>TTACATTACTGCTGATAAGCAAAAGAATGGTATTAAGGCTAATTTCAAGATTAGACATAATATTGAAGATGGTG<br>GTGTTCAATTAGCTGATCATTATCAACAAAATACTCCAATTGGTGATGGTCCAGTTTTGTTGCCAGATAATCATT<br>ATTTGTCTACTCAATCTAAATTGTCTAAAGATCCAAATGAAAAAAGAGATCATATGGTTTTGTTGGAATTTGTTA<br>CTGCTGCTGGTATTACTTTGGGTATGGATGAATTGTACAAA |
| mVenus | ATGTCTAAAGGTGAAGAATTATTCACTGGTGTTGTTCCAATTTTGGTTGAATTAGATGGTGATGTTAATGGTCAT<br>AAATTCTCTGTTTCTGGTGAAGGTGAAGGTGATGCTACTTATGGTAAGTTGACTTTGAAGTTGATTTGTACTACT<br>GGTAAATTGCCAGTTCCATGGCCAACTTTGGTTACTACTTTGGGTTACGGTTTGCAATGTTTTGCTAGATATCCA<br>GATCATATGAAACAACATGATTTCTTTAAATCTGCTATGCCAGAAGGTTACGTTCAAGAAAGAACTATTTTCTTT<br>AAAGATGATGGTAATTACAAAACTAGAGCTGAAGTTAAATTCGAAGGTGATACTTTGGTTAATAGAATTGAATT<br>GAAGGGTATTGATTTCAAAGAAGATGGTAATATTTTGGGTCATAAGTTGGAATACAATTACAATTCTCATAATG<br>TTTACATTACTGCTGATAAGCAAAAGAATGGTATTAAGGCTAATTTCAAGATTAGACATAATATTGAAGATGGT<br>GGTGTTCAATTAGCTGATCATTATCAACAAAATACTCCAATTGGTGATGGTCCAGTTTTGTTGCCAGATAATCAT<br>TATTTGTCTTACCAATCTAAATTGTCTAAAGATCCAAATGAAAAAAGAGATCATATGGTTTTGTTGGAATTTGTT<br>ACTGCTGCTGGTATTACTTTGGGTATGGATGAATTGTACAAA |
| mKO2 | ATGTCTGTTATTAAGCCAGAAATGAAAATGAGATATTATATGGATGGTTCTGTTAATGGTCATGAATTTACTATT<br>GAAGGTGAAGGTACTGGTAGACCATATGAAGGTCATCAAGAAATGACTTTGAGAGTTACTATGGCTGAAGGTG<br>GTCCAATGCCATTCGCTTTTGATTTGGTTTCTCATGTTTTCTGTTACGGTCATAGAGTTTTCACTAAGTACCCAGA<br>AGAAATTCCTGATTATTTTAAGCAAGCTTTTCCAGAAGGTTTATCTTGGGAAAGATCATTGGAATTTGAAGATGG<br>TGGTTCTGCTTCTGTTTCTGCTCATATTTCTTTGAGAGGTAATACTTTCTATCATAAGTCTAAATTTACTGGTGTT<br>AATTTTCCAGCTGATGGTCCAATTATGCAAAATCAATCTGTTGATTGGGAACCATCTACTGAAAAAATTACTGCT<br>TCTGATGGTGTTTTAAAAGGTGATGTTACTATGTATTTGAAATTGGAAGGTGGTGGTAATCATAAATGTCAAATG<br>AAAACTACTTATAAAGCTGCTAAAGAAATTTTGGAAATGCCAGGTGATCATTATATTGGACATAGATTGGTTAG<br>AAAGACTGAAGGTAATATTACTGAACAAGTTGAAGATGCTGTTGCTCATTCT |

These coding DNA sequences were codon-optimized for expression in yeast.

**Supplementary Table 8**. DNA sequences of promoters for multiplexed transcriptional profiling.

| Promoter | Sequence |
|---|---|
| pACT1 | CCTTAAAAACATATGCCTCACCCTAACATATTTTCCAATTAACCCTCAATATTTCTCTGTCACCCGGCCTCTATTTTC<br>CATTTTCTTCTTTACCCGCCACGCGTTTTTTTCTTTCAAATTTTTTTCTTCCTTCTTCTTTTTCTTCCACGTCCTCTTGCA<br>TAAATAAATAAACCGTTTTGAAACCAAACTCGCCTCTCTCTCTCCTTTTTGAAATATTTTTGGGTTTGTTTGATCCTT<br>TCCTTCCCAATCTCTCTTGTTTAATATATATTCATTTATATCACGCTCTCTTTTTATCTTCCTTTTTTTCCTCTCTCTTG<br>TATTCTTCCTTCCCCTTTCTACTCAAACCAAGAAGAAAAAGAAAAGGTCAATCTTTGTTAAAGAATAGGATCTTCTA<br>CTACATCAGCTTTTAGATTTTTCACGCTTACTGCTTTTTTCTTCCCAAGATCGAAAATTTACTGtctagaaa |
| pTEF1 | ATAGCTTCAAAATGTTTCTACTCCTTTTTTACTCTTCCAGATTTTCTCGGACTCCGCGCATCGCCGTACCACTTCAAA<br>ACACCCAAGCACAGCATACTAAATTTCCCCTCTTTCTTCCTCTAGGGTGTCGTTAATTACCCGTACTAAAGGTTTGG<br>AAAAGAAAAAAGAGACCGCCTCGTTTCTTTTTCTTCGTCGAAAAAGGCAATAAAAATTTTTATCACGTTTCTTTTTC<br>TTGAAAATTTTTTTTTTTGATTTTTTTCTCTTTCGATGACCTCCCATTGATATTTAAGTTAATAAACGGTCTTCAATTT<br>CTCAAGTTTCAGTTTCATTTTTCTTGTTCTATTACAACTTTTTTTACTTCTTGCTCATTAGAAAGAAAGCATAGCAATC<br>TAAtctagaaa |
| pGAL1 | ACGGATTAGAAGCCGCCGAGCGGGTGACAGCCCTCCGAAGGAAGACTCTCCTCCGTGCGTCCTCGTCTTCACCGGT<br>CGCGTTCCTGAAACGCAGATGTGCCTCGCGCCGCACTGCTCCGAACAATAAAGATTCTACAATACTAGCTTTTATGG<br>TTATGAAGAGGAAAAATTGGCAGTAACCTGGCCCCACAAACCTTCAAATGAACGAATCAAATTAACAACCATAGG<br>ATGATAATGCGATTAGTTTTTTAGCCTTATTTCTGGGGTAATTAATCAGCGAAGCGATGATTTTTGATCTATTAACA<br>GATATATAAATGCAAAAACTGCATAACCACTTTAACTAATACTTTCAACATTTTCGGTTTGTATTACTTCTTATTCAA<br>ATGTAATAAAAGTATCAACAAAAAATTGTTAATATACCTCTATACTTTAACGTCAAGGAGAAAAAACcccggattctagaaa |
| pGPD1 | CTGGGGTTTGAGCAAGTCTAAGTTTACGTAGCATAAAAATTCTCGGATTGCGTCAAATAATAAAAAAAGTAACCCC<br>ACTTCTACTTCTACATCGGAAAAACATTCCATTCACATATCGTCTTTGGCCTATCTTGTTTTGTCCTCGGTAGATCAG<br>GTCAGTACAAACGCAACACGAAAGAACAAAAAAAGAAGAAAACAGAAGGCCAAGACAGGGTCAATGAGACTGTT<br>GTCCTCCTACTGTCCCTATGTCTCTGGCCGATCACGCGCCATTGTCCCTCAGAAACAAATCAAACACCCACACCCCG<br>GGCACCCAAAGTCCCCACCCACACCACCAATACGTAAACGGGGCGCCCCCTGCAGGCCCTCCTGCGCGCGGCCTCC<br>CGCCTTGCTTCTCTCCCCTTCCTTTTCTTTTTCCAGTTTTCCCTATTTTGTCCCTTTTTCCGCACAACAAGTATCAGAA<br>TGGGTTCATCAAATCTATCCAACCTAATTCGCACGTAGACTGGCTTGGTATTGGCAGTTTCGTAGTTATATATATAC<br>TACCATGAGTGAAACTGTTACGTTACCTTAAATTCTTTCTCCCTTTAATTTTCTTTTATCTTACTCTCCTACATAAGAC<br>ATCAAGAAACAATTGTATATTGTACACCCCCCCCCTCCACAAACACAAATATTGATAAtctagaaa |
| pSSA1 | GGCATTTTCGTTCTTGTGGATTGTTGTAAACTTTCCAGAACATTCTAGAAAGAAAGCACACGGAACGTTTAGAAGCT<br>GTCATTTGCGTTTTTTCTCCAGATTTTAGTTGAGAAAGTAATTAAATTATTCTTCTTTTTCCAGAACGTTCCATCGGC<br>GGCAAAAGGGAGAGAAAGAACCCAAAAAGAAGGGGGGCCATTTAGATTAGCTGATCGTTTCGAGGACTTCAAGGT<br>TATATAAGGGGTGGATTGATGTATCTTCGAGAAGGGATTGAGTTGTAGTTTCGTTTCCCAATTCTTACTTAAGTTGTT<br>TTATTTTCTCTATTTGTAAGATAAGCACATCAAAAGAAAAGTAATCAAGTATTACAAGAAACAAAAATtctagaaa |
| pHSP12 | AGTGAAAATCTCCGGGAGCGGGCGGATCCCACTAACGGCCCAGCCGAAAATGGAAAAAAAGGGTCGGTGATGTGT<br>GGGTGCCAGCTGGCGGTAGCAATGACGACGTGTTGACGGGCCCTTGGCTCTTGGGACAAGGACTAGAAGCCAAAA<br>GCCAGAGGCGGTAAAAATAGCAAGACTAGAATATTGCTGGCATCTGTTAAGGGGATATGTTGCAACTTGCAGGGG<br>GCGGCACAAAATAACATAGAAACGTAGTAAAGAGGGGAAAAGGAAAAGGAAAAGGAAAAGGAAGGAAAAAAAC<br>CCATTGACGTAGAAATTGAAAGAAGGAAAGGTATACGCAAGCATTAATACAACCCACAAACACAGACCAGAAGCA<br>CTCTAGACGGAGAGTAACTAGATCTACAGCCCCTGGAAAATCGTTTGGTCAACTTTGAGGTTCCGGTCGTCCCCCTC<br>TTGATCTGAAAGGTCTTTCTCTAAATCTATATTAAAACGTATAAATAGGACGGTGAATTGCGTTCTACTTCCTCAAT<br>TGCGTTTGATCTTATTTAATCTCTCTCTAATATATAGAAAAAAAAACCATCTGATTATTCGATAATCTCAAACAAAC<br>AACTCAtctagaaa |
| pOLE1 | CATGTCCCGGGGTTAGCGGGCCCAACAAAGGCGCTTATCTGGTGGGCTTCCGTAGAAGAAAAAAAGCTGTTGAGCG<br>AGCTATTTCGGGTATCCCAGCCTTCTCTGCAGACCGCCCCAGTTGGCTTGGCTCTGGTGCTGTTCGTTAGCATCACAT<br>CGCCTGTGACAGGCAGAGGTAATAACGGCTTAAGGTTCTCTTCGCATAGTCGGCAGCTTTCTTTCGGACGTTGAACA<br>CTCAACAAACCTTATCTAGTGCCCAACCAGGTGTGCTTCTACGAGTCTTGCTCACTCAGACACACCTATCCCTATTG<br>TTACGGCTATGGGGATGGCACACAAAGGTGGAAATAATAGTAGTTAACAATATATGCAGCAAATCATCGGCTCCTG<br>GCTCATCGAGTCTTGCAAATCAGCATATACATATATATATGGGGGCAGATCTTGATTCATTTATTGTTCTATTTCCAT<br>CTTTCCTACTTCTGTTTCCGTTTATATTTTGTATTACGTAGAATAGAACATCATAGTAATAGATAGTTGTGGTGATCA<br>TATTATAAACAGCACTAAAACATTACAACAAtctagaaa |
| pTRX2 | ACTTTTACGGGTGGCAACGGAACCAACGTATTTAGAGATTGTTTTTTGGTCAAGCGAGGAACCCCTGTTGGCAAAG<br>TTGCCAGGTATATCATGGGTGGCGAGGTCACCATTGCAAGCATTGAAACCGTTGGCGGCGTGAGAGTCAGTGAAGA<br>AAGTCTTGTTGAGCCCGGTAAGAATGACATACTCGGCTTCAAGATCGCTCCAAGATCAGCATAACTTGAGTGCCAG<br>TGAATATTAAGTAATCATCAAAGTATATGTGTAATTGTTTATACTCTTAGTAAAGGATGCTCCCTACAAGGTGGCTC<br>TTTTCTTACTAAGCGCGTTCAGTTTCCAGCCAGCCGAAAGAGGGATATCAGTATATAAGAAAGCCATTCGGGGGAT<br>GAAAAGCTGACAAGAGAATAACGAGGACCAGTTTTTATTTGTTGTCTAGCAAGAATTATACACGCACACATACACG<br>AGAGTCTACGAtctagaaa |
| pFET3 | GATAATGCCTTGGCTTGCCTATTTCACGGTTACAGGAATAATAACATGTTCATACCGTTTCAGGACCAATAAATGGT<br>CTTCTGCGAGAGAAAAAGGACACTCTCCGTCCGACAGAAATAAGCTTTACTTTCCGGGTGCGAATCAGCCCGTTGC<br>GCCTGGGGTGGTCCCTACAGTACGCTGAGTCGCCGATAAAGACCCTCCGCCTAGAGCTAGGCGAGGCTGACTTAGG<br>CAGGCCCAACAGGCAAGGCCCATCTTCAAAAGTGCACCCATTTGCAGGTGCTCTTATTCTCGCCAATTGCGACAGA<br>AAATGAAGGATGCACTCAAACAGTCGATCCTTCGAGGGAGTATGCCAAGGCCTCGTGCATGTAGTGCGATTATATA<br>TATATATATATATATATATATATGTATGTAAGCAGGCCATGCCCTATAGCTCTTGTTCTATAAGCGATGGATAGGCA<br>TAGGAAACGAAGAGGACCCCAGTGTAAGGAAGAGTAtctagaaa |

pERO1

AAAGAACACGGCGGTAAGAATACGTTCCTTTTTGTGCTGTGTACACCCGTAAAATTGTACATTATTTATTTCAAAAT
ATATAACAGGATCCCTCCAGTGTGTCTGAAATGATGCAACTCTGATACTTCAGAGTCGTACCCTTATTAAATACTAA
TGCCGAATATAGTCATCCAGTAGCCATAGTTCACACACACATTACTTATTCACCACATAAAGAACTAGAAATGCTTG
CTAAATATCGTACCTTGAAGGTAAACATTAAGGCCCCCCAAAAGCATATATACATACTGGCAACACAAAAAAAGTA
AATTGCCAACCACCTACCTTACATATGGTTTGCCCATCCTACATTACCAATACTATCAAGACATTTCTTCTGAAACA
TATTCACAACTGAAACGAGATCATTTTCTTATCTATCTATTGAGTAATGCTTACTTTTCATATTTTCAATGAACAATA
GGATATGTAGGAGAATTGATATATTCACTGCGTATCAGAGAAAAGGTCTACTGACATTTTATGGCAAATGTATTCTA
CACAAATCGAGAATACCACAGACAATGGTACAAGACATACACAAAGAGAAGACTGTTCTAATTAAACAAATAATA
TTGAGCTACCTGCTAAGTATGTCCTTTTCCCTTTGTCCTTTGGTTTCTCTTATAGAAGACCCTGGAAATTTTTCGCATT
TTTCCGGCTTTGGGCGTTAGTAAGAACAAAAAGAAAAGAAGAGAACAAAAAAGAAACGATACGGAGTACGTGTCA
TAAAAACTTGTTCAATCATCCTTGAAGCTAAGTATAAAGAGCTTGAAAAGGTTTACCACTTAAACTGGTTATACTAT
TTCAAGAGTGTAAACATTTTATTGCATATACCACAGTAACGTGCAGGTAAAtctagaaa

pDAL5

AGCGTTCTCATCAGTCACTTGACAAATGCTCGAGGAGCTATCATTTGCTGATAAGGTGCTACAGCGCGCTCCTGCCG
CACGCTTTGTTCCTTTTCGATAAGAGTCCCTCGCGTTAGTCTGAGTGAAGTGCGGAATTCAGCAAACGAATAACAAT
CGACCTTATGATCATGTGGATTATCGGGGCAAAAGATTTGGCCAAGATGTCAGAGAACGTTATCACCAATCACTCA
CACAATTAAGTGGTAGTGTAACTCCGAAGATACGGCTAATACTTATCATTATCTGGTTTTCCGAATATACAGATTGG
ATGAAGTAATATATGTATATAAATGGACCAAGGAAACATCAAATTAGGAGATCATGAGGGAAAGGTTTAACATAA
CAACATTGAAGAAAACAACAAAACAAGGATtctagaaa

pHIS4

AAACCCATGCACAGTGACTCACGTTTTTTTATCAGTCATTCGATATAGAAGGTAAGAAAAGGATATGACTATGAAC
AGTAGTATACTGTGTATATAATAGATATGGAACGTTATATTCACCTCCGATGTGTGTTGTACATACATAAAAATATC
ATAGCACAACTGCGCTGTGTAATAGTAATACAATtctagaaa

pZRT1

GGCAAGAGTATTTCAGACTTTCCTAATATGAAAGGACAAATTGACACTAATGTCTGATTATGGCCAATTCCTGCGGT
AAATTACACGGCGATTACGGCGACATGAGCTCACATTCATCACTCTATGGGACAAATGTTTCCAAACTGGGCGCAA
CAAACACCTGATGTGACTCCTACCCTTTGGACAATGCAGATCCACGCTACGGCAAATTAGTCAAATGCACTAGAAC
ATGGCGCAAGTACTTATTGTGACCTTTGGGGTACCGTTACCGTCAGTTTTCTTCAGCTAAGGCGCGCGCGCCAGATA
ACTAAAAAAAAATATAGTTGCTGCTTAAAAAACAATACACCCGTACTCTCTTGCCTGTAAAAACCTCGAAGGACCA
AAGATACCCTCAAGGTTCTCATCTGTGCGGTATTCTTCAAATTACAATGACATTTCCCAAAATTATCAGATGTGCTC
AGGTATCTTCTCTCCAATGAGATGAGACAGATGAACATATTTGACCTTGAAGGTCATGGAAAGTAGGTTGAGAGCA
AATGTGTAGAACGAAATTAAGAAAAAAAGAAATTACGCACGGCATTAGCTCGATGACTTAGTTATAAATAGAGGC
CTGGTATCGGCTGTCATGATCTCATCTCTTCCCTATTTACAAAAAAACTGCAAGTATAGACAATAAAACAACAGCAC
AAAtctagaaa

pARR3

CACGTGCAAAATCTTCTCTTCGAAGATCAACAACTTGAAAATCCTTCCTCTGATTTTCAATTAGGCCCTTGAGTTGC
CTAGACGTTATGAAACTTACCATTACGCTTGCTGGATTGTCAAGTTTTCCTCAATATTAGTTTGTTATACATTAGTTT
TTATACCGACTTTTCAAAAGTGTTGATTTAAAATCAAATTTTGAATGCTCTTAATTATCTTTTTGTTTGATTAATAAT
CAACTTTAGCGGCAACGCTCCTTACATAATTATAATGTAAACGGAAAAGAATATAAATGAATGCTCTCGTTGTAATT
CAAGAGAACCCAACCAACAAATCATCAGGtctagaaa

pRNR3

GTAATAACAAGCAGGTGGGCGCTTTGAAGAGTATGGTAGAGGATAAGATCCAGAAGGAAACACTCAAGGGTGTTG
TCGTCGCTGGAGGCGTACTAGCCGGCGCTGTGGCCGTGGCTAGTTTCTTCTTAAGAAACAAGAGAAGGTAACAAGC
ACATAAAAAATCAGCACATACGTACATACATAAGAATGAATCGCACGCACGCGTAAACATTTATCATTTAATCTTC
AGTTGTTAGATAAAAAAAAAAAGAAAAGAAAAGAAAGTGAAGGCTTGTTTCAGTTTGAACTAGGTAGCAGAGCAA
GCCCTCGTTCTTGGCTGCTAATTTTCCTAAAGTAGTAAAAAAAGCCAAGTTATCTGCCTACGGTTGTCACAGCAACA
TTGCGTGCCGTTGTTCTTTTGTTTTTTTTTTTTTTTTTTTTTCGTGGTTGTCGCAGCAACGACACCTAGGCGCTGCTCA
AAGGGGCAAAAACCCGGTTGCCATGGCGAGGACCAAACGACAAGATGGGAAAAAAACAATAGTCTATTGTTAAAT
CGTAATACTGTATTGTGAGATGCTGACGCGTTTCGTTTTTCGTGTCAGCGTTCTTTATATTGTTTCGTGTTCTGCTGC
AAAACGTATATAAACGCACTGCTATTTTGCCTTCTTTTGCCTTCTTCCTTGCCTTTCTCTCATCTCATATCCAAGTTGA
AATAAATATGACAAGCAAGAATAGCAGCAGCAATAAtctagaaa

pCUP1

TCACCACCCTTTATTTCAGGCTGATATCTTAGCCTTGTTACTAGTTAGAAAAAGACATTTTTGCTGTCAGTCACTGTC
AAGAGATTCTTTTGCTGGCATTTCTTCTAGAAGCAAAAAGAGCGATGCGTCTTTTCCGCTGAACCGTTCCAGCAAAA
AAGACTACCAACGCAATATGGATTGTCAGAATCATATAAAAGAGAAGCAAATAACTCCTTGTCTTGTATCAATTGC
ATTATAATATCTTCTTGTTAGTGCAATATCATATAGAAGTCATCGAAATAGATATTAAGAAAAACAAACTGTACAAT
CAATCAATCAATCATCACAtctagaaa

pFUS1

TGCCTCAATCCTTCTTTTGCTTCCATATTTACCATGTGGACCCTTTCAAAACAGAGTTGTATCTCTGCAGGATGCCCT
TTTTGACGTATTGAATGGCATAATTGCACTGTCACTTTTCGCGCTGTCTCATTTTGGTGCGATGATGAAACAAACAT
GAAACGTCTGTAATTTGAAACAAATAACGTAATTCTCGGGATTGGTTTTATTTAAATGACAATGTAAGAGTGGCTTT
GTAAGGTATGTGTTGCTCTTAAAATATTTGGATACGACATCCTTTATCTTTTTTCCTTTAAGAGCAGGATATAAGCCA
TCAAGtctagaaa

pPRM5

CTCACCCGGATCGTAGTCACATGATCAAATAAATTATTGCATTACCAATGGCTTCTGTATTAGTTACTGTCCAGGAA
AGGTCTCAATATAACCGGTCACCTTATTTATGATAACAATTTTTAACCATTTACCCTTTATTTTTGCAAAGTTATGAC
CTTTGGAATGCGGCAGAAAAAAAAAAAAATTGATGAAGTAGTCATCAAACAGGTTTCGGCGAAAGACAGTACAAGA
ATTGCCAGCTAAAGCTTTTCTAATATGTTATTCCATCAAATATTCACGCTATTAATGCTATCTGATCGATTACTTGGT
AAAAATAACCAAATGAGTATTGGGTCATATTTTGGAGAAGCGAAAAGTGCACGGCATACTTAAAGAAGAGAAAAA
AATTCTTAATCAAAGCTTAAGCGGCATCGCATAACTCTGACAAAAGTATAATCATAGTATGTGGTTTGAGTAAACA
ACAAGACTGGCAGGTAAGTTTTTTAAATGAATTTTCTTAGCAGGGTTATTGCGGCGCACGTCTATAGGTGATTTACC
TTTGTTCCTGAATATACATGATATTCTTAATGCGACAGCGCCCAGGGAAGCAAAGAACGCCGACGAAAGAGCGGCG
GAGCAAAAGGCGCAAAGTGACACCTTCCGATCCTGGGAAGTGATCGGTTTCGCTGTTCGGGCTTTTCTTAGACAAA
TAAAGTTTAATTTTCTTGCTTTCTCTTTTTGGCTAAGAATTCTATATTGTTCTTGAAACAACTTTGTCACACGTTCTAA

| | |
|---|---|
| | AAATAAAATGGAAAATGGAAAATGAAAAATTAGAGAGAAAATGTATTACTGAAGAATGTGGTAAAGACATTTGAA<br>AAATATTTAATATATTACCTGAGATCATTCAAAGGAACGCATCGGTGCAAAGAAACAGTTATAGCATTAGAGTTAA<br>GTCAAGAGCACTCATTATTACTTTAAAAGGTCGTTGAAAATAGAATAAATTGACACTCAAAACGCAAGAtctagaaa |
| pHXT1 | TGCAAAAAGCTTCCGATCCTCAAATACAGTGAGAGAAAAAGCAAACTTGTCTTCATTATTTTTCTGTTTGGCCCTGT<br>CACGGTTCTTTTTATGGCCTTCTCGAAGGATATCCGTAGTCAATTATATTCACGTAGTTGCCAAAAGTAATTTTTGGA<br>AAACTATTATTCCTCCGAGAAAACCTCACACAGAAATCCTTGCAGGTCTCATCTGGAATATAATTCCCCCCTCCTGA<br>AGCAAATTTTTCCTTTGAGCCGGAATTTTTGATATTCCGAGTTCTTTTTTTCCATTCGCGGAGGTTATTCCATTCCTA<br>AACGAGTGGCCACAATGAAACTTCAATTCATATCGACCGACTATTTTTCTCCGAACCAAAAAAATAGCAGGGCGAG<br>ATTGGAGCTGCGGAAAAAAGAGGAAAAAATTTTTTCGTAGTTTTCTTGTGCAAATTAGGGTGTAAGGTTTCTAGGG<br>CTTATTGGTTCAAGCAGAAGAGACAACAATTGTAGGTCCTAAATTCAAGGCGGATGTAAGGAGTATTGGTTTCGAA<br>AGTTTTTCCGAAGCGGCATGGCAGGGACTACTTGCGCATGCGCTCGGATTATCTTCATTTTTGCTTGCAAAAACGTA<br>GAATCATGGTAAATTACATGAAGAATTCTCTTTTTTTTTTTTTTTTTTTTTTTTTTTACCTCTAAAGAGTGTTGACCAAC<br>TGAAAAAACCCTTCTTCAAGAGAGTTAAACTAAGACTAACCATCATAACTTCCAAGGAATTAATCGATATCTTGCA<br>CTCCTGATTTTTCTTCAAAGAGACAGCGCAAAGGATTATGACACTGTTGCATTGAGTCAAAAGTTTTTCCGAAGTGA<br>CCCAGTGCTCTTTTTTTTTTTCCGTGAAGGACTGACAAATATGCGCACAAGATCCAATACGTAATGGAAATTCGGAA<br>AAACTAGGAAGAAATGCTGCAGGGCATTGCCGTGCCGATCTTTTGTCTTTCAGATATATGAGAAAAAGAATATTCA<br>TCAAGTGCTGATAGAAGAATACCACTCATATGACGTGGGCAGAAGACAGCAAACGTAAACATGAGCTGCTGCGAC<br>ATTTGATGGCTTTTATCCGACAAGCCAGGAAACTCCACCATTATCTAATGTAGCAAAATATTTCTTAACACCCGAAG<br>TTGCGTGTCCCCCTCACGTTTTTAATCATTTGAATTAGTATATTGAAATTATATATAAAGGCAACAATGTCCCCATA<br>ATCAATTCCATCTGGGGTCTCATGTTCTTTCCCCACCTTAAAATCTATAAAGATATCATAATCGTCAACTAGTTGATA<br>TACGtctagaaa |
| pPMC1 | GTTTTTACCCGGCAAAGAAGCTTCTCCATCATTTGTCAGGGGGAAAAGAAAATTAGGAGGCTCAGGCCCTAGAAGC<br>GCCCTCCACGCTTTTTAAACAAATGCTAATCTTCATAATTCATTATCATCGCCACTTGGCATTGCATTTAATGGGCGC<br>CTTCCAAAAACATTCAAATAGTATCAGCCCCCCCCCAAGGAAAGGCTTTATTTGAATTGCGTTAAATATTCGATTTC<br>TTTCTAGCAATTATTACCAAAATAAATTGCAGGCTTAACGAAAAATTAAATAATCTCTTCAGATTTTTTCACTTCCCC<br>AAACTACATTTTAGTCAGGTTGGCACGATTCAATTGAGAAGGCTTAGGTTAAGAATAAACATTTTTTCATAAATTTG<br>AGGAAGAAGGGGCTTTTCTGGTAATTTTCATTCAATAGAAAGGCTTAAGAAAACGTCAAAAAACTCATGCGCCTCC<br>AGTAAAAACTAAGTGTATTAAGAGAATTGTATCACATATATACGACATTGAATTAAAGAAATAAAACTTGAACAAA<br>TAAAGTTTAGAAAAGTGGTTCTAAAAAAAAAAAAAACTGTGTGCGTAACAAAAAAAAATAtctagaaa |

All promoters were cloned to include a short non-native sequence placed immediately upstream of start codon (underlined and lowercase), which includes an *XbaI* site.

**Supplementary Table 9**. DNA sequences of SynGal4 constructs for co-culture tracking.

| Part | Sequence |
|---|---|
| Gal4-BD | ATGAAGCTACTGTCTTCTATCGAACAAGCATGCGATATTTGCCGACTTAAAAAGCTCA<br>AGTGCTCCAAAGAAAAACCGAAGTGCGCCAAGTGTCTGAAGAACAACTGGGAGTGTC<br>GCTACTCTCCCAAAACCAAAAGGTCTCCGCTGACTAGGGCACATCTGACAGAAGTGGA<br>ATCAAGGCTAGAAAGACTGGAACAGCTATTTCTACTGATTTTTCCTCGAGAAGACCTT<br>GACATGATTTTGAAAATGGATTCTTTACAGGATATAAAAGCATTGTTAACAGGATTAT<br>TTGTACAAGATAATGTGAATAAAGATGCCGTCACAGATAGATTGGCTTCAGTGGAGAC<br>TGATATGCCTCTAACATTGAGACAGCATAGAATAAGTGCGACATCATCATCGGAAGAG<br>AGTAGTAACAAAGGTCAAAGACAGTTGACTGTATCG |
| Gal4-AD | GCCAATTTTAATCAAAGTGGGAATATTGCTGATAGCTCATTGTCCTTCACTTTCACTAA<br>CAGTAGCAACGGTCCGAACCTCATAACAACTCAAACAAATTCTCAAGCGCTTTCACAA<br>CCAATTGCCTCCTCTAACGTTCATGATAACTTCATGAATAATGAAATCACGGCTAGTA<br>AAATTGATGATGGTAATAATTCAAAACCACTGTCACCTGGTTGGACGGACCAAACTGC<br>GTATAACGCGTTTGGAATCACTACAGGGATGTTTAATACCACTACAATGGATGATGTA<br>TATAACTATCTATTCGATGATGAAGATACCCCACCAAACCCAAAAAAAGAG**TAA** |
| fsA | GGATCCGGCAGTGGTTCTGGCGCACTACAAGGACGACGACACAAGACTTTAAACTAGT<br>TGACGCGGGTCTAGCCGTTCCCGCGTTAAACGTACTAGAAGGCGGTTCTATGGGAATG<br>TCTGGAAAGCTT |
| fsB | GGATCCGGCAGTGGTTCTGGCGCACTACAAGGACGACGACACAAGACTTTAAACTAGT<br>TGACGCGGGTTTAAGTACGGTCGCGTTAAACTCGCTAGAAGGCGGTTCTATGGGAAAT<br>GTCTGGAAAGCTTGCCAATTT**TAA***|
| fsC | GGATCCGGCAGTGGTTCTGGCGCACTACAAGGACGACGACACAAGACTTTAAACTAGT<br>TGACGCGAGTCTAATGTCTGCCGCGTTAAACTCGCTAGAAGGCGGTTCTATGGGAATG<br>TCTGGAAAGCTT |
| fsD | GGATCCGGCAGTGGTTCTGGCGCACTACAAGGACGACGACATAAGACTTTAAACTAGT<br>TGACGCGGTTCTAGGTCAACCCGCGTTAAACATCCTAGAAGGCGGTTCTATGGGAATG<br>TCTGGAAAGCTT |
| fsE | GGATCCGGCAGTGGTTCTGGCGCACTACAAGGACGACGACACAAGACTTTAAACTAGT<br>TGACGCGGATCTAGGAGGTATCGCGTTAAACAGCCTAGAAGGCGGTTCTATGGGAATG<br>TCTGGAAAGCTT |
| fsF | GGATCCGGCAGTGGTTCTGGCGCACTACAAGGACGACGACACAAGACTTTAAACTAGT<br>TGACGCGTCGCTACCAGTGGGCGCGTTAAATGATCTAGAAGGCGGTTCTATGGGAATG<br>TCTGGAAAGCTT |
| fsG | GGATCCGGCAGTGGTTCTGGCGCACTACAAGGACGACGACACAAGACTTTAAACTAGT<br>TGACGCGGGACTAGGCTTTGCCACGTTAAACAGCCTAGAAGGCGGTTCTATGGGAATG<br>TCTGGAAAGCTT |
| fsH | GGATCCGGCAGTGGTTCTGGCGCACTACAAGGACGACGACACAAGACTTTAAACTAGT<br>TGACGCGGGTCTAGCACCTCGCGCGTTAAACGTGCTAGAAGGCGGTTCTATGGGAATG<br>TCTGGAAAGCTT |
| fsI | GGATCCGGCAGTGGTTCTGGCGCACTACAAGGACGACGACACAAGACTTTAAACTAGT<br>TGACGCGGGACTAGGCATATGCGCGTTAAACAACCTAGAAGGCGGTTCTATGGGAAT<br>GTCTGGAAAGCTT |

SynGal4 constructs are assembled as Gal4-BD • fs • Gal4-AD. Frameshift slippery sites are shown underlined.
*This sequence is mutated and contains a stop codon (bold) in the -1 frame shortly following the canonical fs module sequence.

**Supplementary Data 1**. Example raw flow cytometry data of FRAME-tagged strains.
File name: **FTs_supp-data_1.xlsx**
Description: Raw flow cytometry event data (channels FSC.A, SSC.A, GFP.A, RFP.A) pre-gated to remove events with extreme scatter and fluorescent values (nonMaxF gate in **Supplementary Figure 8**). The derived parameters normalized GFP (normGFP), normalized RFP (normRFP), GFP+RFP fluorescence total (fluorTotal), RFP/GFP fluorescence ratio (fluorRatio) as well the FRAME-tag identity (FT) assigned by the FRAME-finder code are also included for each event.

**Supplementary Data 2**. Example raw microscopy data of FRAME-tagged strains.
File name: **FTs_supp-data_2.xlsx**
Description: Raw microscopy cytometry data based on image segmentation of fields captured on a Ti-E microscope as described in the Methods. Each row includes the fluorescence (GFP and RFP) and geometric (Area and Feret) values of a sectioned area corresponding to a cell. Geometric measurements are given in pixel units (1 px = 0.33μm). For each sectioned area, the FRAME-tag identity (FT) assigned by the FRAME-finder code is also included.

**Supplementary Data 3**. Raw data of frameshift efficiency and percent overlap.
File name: **FTs_supp-data_3.xlsx**
Description: Raw fluorescence data used to calculate frameshift efficiency in **Supplementary Figure 2c** as well as the *fs* distribution overlap data used to plot the heatmap in **Supplementary Figure 2d**.

**Supplementary Data 4**. Calculated Sensitivity, Specificity, PPV, NPV values of automated gating.
File name: **FTs_supp-data_4.xlsx**
Description: Calculated numeric values used to generate the Sensitivity, Specificity, PPV and NPV plots in **Supplementary Figure 9** as described in the Methods (Statistical Analysis).

**Supplementary Data 5**. Calculated PPV values of automated gating of mock diluted strains.
File name: **FTs_supp-data_5.xlsx**
Description: Calculated numeric values used to generate the PPV plots of mock diluted strains in **Supplementary Figure 10** as described in the Methods (Statistical Analysis).
